# HSD3B1 is an Oxysterol 3β-Hydroxysteroid Dehydrogenase in Human Placenta

**DOI:** 10.1101/2022.04.01.486576

**Authors:** Alison Dickson, Eylan Yutuc, Catherine A Thornton, James E Dunford, Udo Oppermann, Yuqin Wang, William J Griffiths

## Abstract

Most biologically active oxysterols have a 3β-hydroxy-5-ene function in the ring system with an additional site of oxidation at C-7 or on the side-chain. In blood plasma oxysterols with a 7α-hydroxy group are also observed with the alternative 3-oxo-4-ene function in the ring system formed by ubiquitously expressed 3β-hydroxy-Δ^5^-C_27_-steroid oxidoreductase Δ^5^-isomerase, HSD3B7. However, oxysterols without a 7α-hydroxy group are not substrates for HSD3B7 and are not usually observed with the 3-oxo-4-ene function. Here we report the unexpected identification of oxysterols in plasma derived from umbilical cord blood and blood from pregnant women taken before delivery at 37+ weeks of gestation, of side-chain oxysterols with a 3-oxo-4-ene function but no 7α-hydroxy group. These 3-oxo-4-ene oxysterols were also identified in placenta, leading to the hypothesis that they may be formed by a previously unrecognised 3β-hydroxy-Δ^5^-C_27_-steroid oxidoreductase Δ^5^-isomerase activity of HSD3B1, an enzyme which is highly expressed in placenta. Proof of principle experiments confirmed that HSD3B1 has this activity. We speculate that HSD3B1 in placenta is the source of the unexpected 3-oxo-4-ene oxysterols in cord and pregnant women’s plasma and may have a role in controlling the abundance of biologically active oxysterols delivered to the fetus.

## Introduction

Oxysterols are oxidised forms of cholesterol or of its precursors (1). The primary routes of oxysterol metabolism are 7α-hydroxylation catalysed by cytochrome P450 (CYP) 7B1 (2), or in the specific case of 24S-hydroxycholesterol (24S-HC) by CYP39A1 (3), and (25R)26-hydroxylation or carboxylation catalysed by CYP27A1 (Figure 1) (4-6). Once 7α-hydroxylated, oxysterols become substrates for the ubiquitous hydroxysteroid dehydrogenase (HSD) 3B7 (7-9), which oxidises the 3β-hydroxy group to a 3-ketone and isomerises the double bond from Δ^5^ to Δ^4^ (Figure 1), a key reaction in bile acid biosynthesis necessary for conversion of initial 3β-hydroxy stereochemistry, as in the cholesterol structure, to the 3α-hydroxy stereochemistry in primary bile acids (5, 6, 10, 11). Cholesterol itself is 7α-hydroxylated by CYP7A1 to 7α-hydroxycholesterol (7α-HC) (12), which like other oxysterols with a 7α-hydroxy group, is a substrate for HSD3B7 (5). With respect to oxysterols, including down-stream sterol-acids, oxidation at C-3 with accompanying Δ^5^-Δ^4^-isomerisation can be regarded as a deactivation mechanism eliminating many of the biological activities of the substrate oxysterol. This is the case for the chemoattractant oxysterols 7α,25-dihydroxycholesterol (7α,25-diHC) and 7α,(25R)26-dihydroxycholesterol (7α,26-diHC, also known as 7α,27-dihydroxycholesterol) (13), and the liver X receptor (LXRα and LXRβ) ligand 3β,7α-dihydroxycholest-5-en-(25R)26-oic acid (3β,7α- diHCA) (14). Note, in much of the literature (25R)26-hydroxylation and (25R)26-carboxylation are described according to non-systematic nomenclature as 27-hydroxylation and 27-carboxylation, with the stereochemistry assumed to be 25R (15). Here we prefer to use systematic numbering, and for brevity when hydroxylation or carboxylation is at C-26 the reader can assume that the stereochemistry is 25R unless stated otherwise i.e., 26-HC is used as the abbreviation for (25R)26-hydroxycholesterol. While HSD3B7 has activity towards 7α-hydroxy-oxysterols it has not been reported to oxidise oxysterols lacking the 7α-hydroxy group (8). Never-the-less deficiency in HSD3B7 does not completely eliminate primary bile acid production (16), suggesting that a second hydroxysteroid dehydrogenase may have 3β-hydroxy-Δ^5^-C_27_ steroid oxidoreductase activity in human.

**Figure 1.**
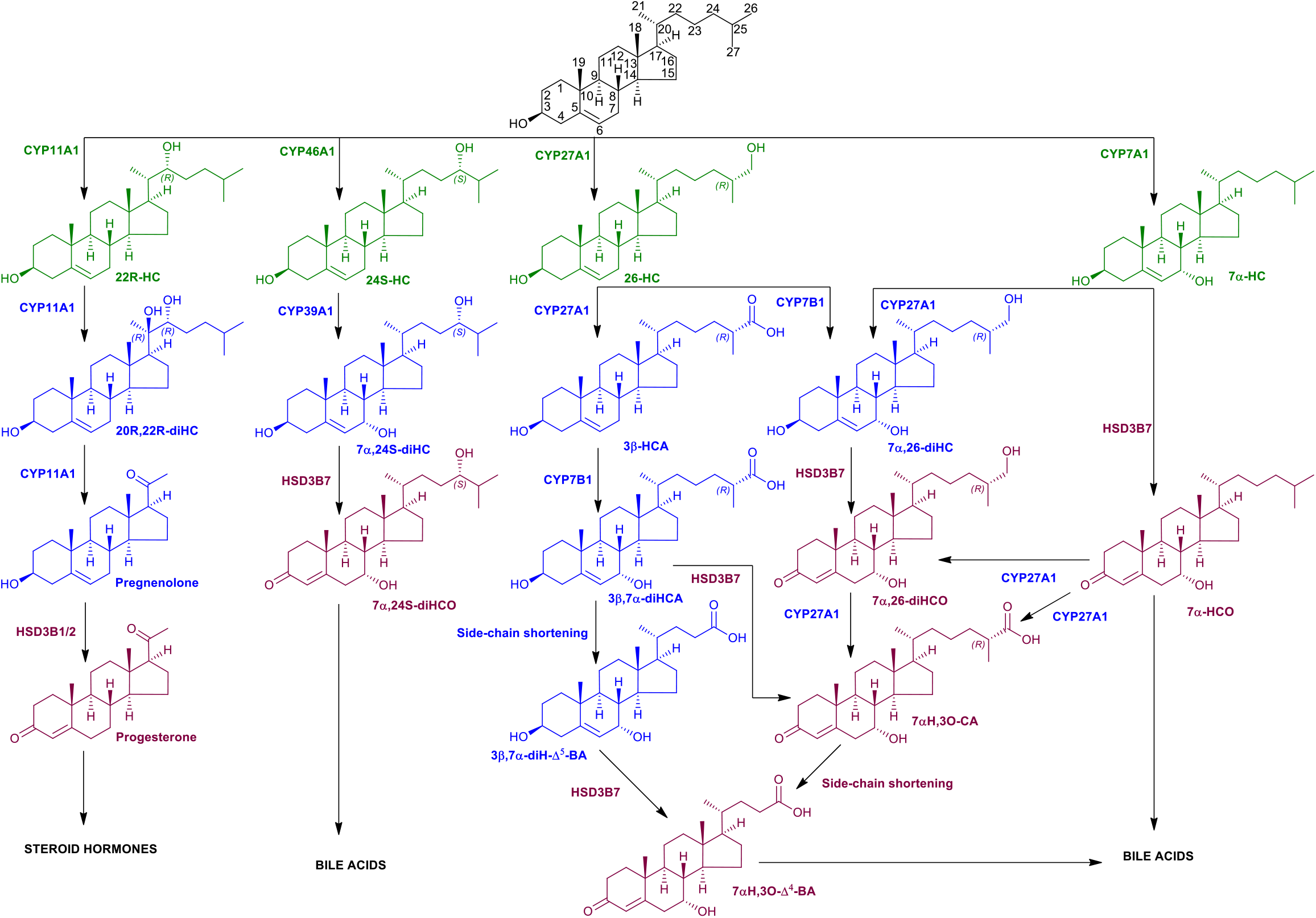
Abbreviated scheme of oxysterol metabolism. Primary oxysterols (hydroxycholesterols, HC) are in green, secondary oxysterols (diHC), including 3β-hydroxycholestenoic acids, are in blue, 3-oxo- 4-ens are in claret.

In humans the HSD3B1 and HSD3B2 enzymes belong to the evolutionarily conserved superfamily of short-chain dehydrogenases/reductases (SDR) (17), with official nomenclature symbols SDR11E1 and SDR11E2, respectively (18). They convert C_19_ and C_21_ steroids with a 3β-hydroxy-5-ene structure to 3-oxo-4-ene products (Figure 1). HSD3B1 is primarily localised to placenta and non-steroidogenic tissue, while HSD3B2 is primarily expressed in the adrenal gland ovary and testis (9, 19). The 3β-hydroxy-5-ene to 3-oxo-4-ene transformation is an essential step in the biosynthesis of all classes of active steroids (20, 21). HSD3B1/2 enzymes are not reported to use C_27_ steroids as substrates although they share about 34% sequence identity to HSD3B7 (SDR11E3) in human, and HSD3B7 does not use C_19_ or C_21_ steroids as substrates suggesting different physiological roles for these HSD enzymes (8).

Two side-chain oxysterols that have not been reported to be 7α-hydroxylated *in vivo* are 22R-hydroxycholesterol (22R-HC) and 20S-hydroxycholesterol (20S-HC). Instead, 22R-HC becomes hydroxylated to 20R,22R-dihydroxycholesterol (20R,22R-diHC) which then undergoes side-chain shortening to pregnenolone in reactions catalysed by CYP11A1 (Figure 1) (22, 23). 20S-HC has also been reported to be converted to pregnenolone (1). Pregnenolone is converted by HSD3B1/2 enzymes to progesterone. 20S-HC, 22R-HC, and 20R,22R-diHC are ligands to LXRs (24-26), while 22R-HC, like many other side-chain hydroxylated 3β-hydroxysterols, is a ligand to INSIG (insulin induced gene), important in the regulation of SREBP-2 (sterol regulatory element-binding protein-2) processing and cholesterol biosynthesis (27). 20S-HC also regulates SREBP-2 processing (28), presumably by binding to INSIG, and is a ligand to the G protein-coupled receptor (GPCR) Smoothened (SMO) important in the Hedgehog (Hh) signalling pathway (29), and has recently been reported to be a ligand to the sigma 2 (σ2) receptor TMEM97 (30). These biological activities are not known to be conveyed to the 3-oxo-4-ene analogues of 20S-HC, 22R-HC or 20S,22R-diHC, again suggesting that oxidation at C-3 with accompanying Δ^5^-Δ^4^ isomerisation may be a deactivation mechanism of oxysterols.

While oxysterols, including sterol-acids, based on a 3β-hydroxy-5-ene framework are routinely analysed by both gas chromatography – mass spectrometry (GC-MS) (31-33) and liquid chromatography (LC) – MS (31, 32, 34-36), with the exception of 7α-hydroxycholest-4-en-3-one (7α-HCO) and 7α-hydroxy-3-oxocholest-4-en-(25R/S)26-oic acid (7αH,3O-CA), this is not normally the case for the 3-oxo-4-ene sterols (31, 37-39). The “enzyme-assisted derivatisation for sterol analysis” (EADSA) technology, as used in the current study, allows the analysis of 3β-hydroxy-5-ene and 3-oxo-4-ene oxysterols, including sterol-acids, in a single LC-MS run (31, 40-43). In this technology, after extraction each sample is split into two equal aliquots; to the B-fraction endogenous 3-oxo-4-ene sterols are reacted with the [^2^H_0_]Girard P hydrazine (GP) reagent to tag a charge to the sterol skeleton to enhance sensitivity of LC-MS analysis; while to the A-fraction bacterial cholesterol oxidase enzyme is added. This converts sterols with a 3β-hydroxy-5-ene function to their 3-oxo-4-ene equivalents which are then reacted with [^2^H_5_]GP reagent (see Supplemental Figure S1). A salient feature of the method is that *in fraction-B only sterols with a natural oxo group are derivatised* (*with [*^*2*^*H*_*0*_*]GP*), while *in fraction-A sterols with a natural oxo group and those oxidised by cholesterol oxidase to contain one are derivatised (with [*^*2*^*H*_*5*_*]GP)*. The derivatisation products are then combined and analysed by LC-MS, the originating structure of sterol, whether 3-oxo-4-ene or 3β-hydroxy-5-ene, is revealed by the isotope labelling of the GP reagents and deconvolution of the resultant data from A- and B-fractions (40).

3β-Hydroxycholest-5-en-(25R)26-oic acid (3β-HCA) does not have a 7α-hydroxy group and is a C_27_ sterol so should not be a substrate for HSD3B7 or HSD3B1/2 enzymes, however, 3-oxocholest-4-en-(25R)26-oic acid (3O-CA) is found at low levels in human plasma at about 1 – 5% of 3β-HCA (31, 41-43). Here we report evidence for the presence of (25R)26-hydroxycholest-4-en-3-one (26-HCO), a precursor of 3O-CA, in human plasma and its elevated concentration in plasma from pregnant women taken 1 – 2 days prior to elective caesarean section at 37+ weeks of gestation and in plasma generated from blood of the umbilical cord. Besides 26-HCO and 3O-CA we identify other C_27_ - C_24_ cholesterol metabolites with a 3-oxo-4-ene structure but no 7α-hydroxy group in these plasmas and in human placenta. HSD3B1 is abundant in human placenta (19), and we present proof-of-principle data that demonstrates that the HSD3B1 enzyme will convert monohydroxycholesterols, where the added hydroxy group is in the side-chain, into hydroxycholest-4-en-3-ones.

## Materials and Methods

### Materials

The source of all materials for oxysterol analysis can be found in reference (43).

### Human material

Maternal blood was taken 24 - 48 hr prior to elective caesarean section at 37+ weeks of gestation for reasons that did not include maternal or fetal anomaly. Umbilical cord blood and placenta were collected at delivery of the baby. Control plasma was from non-pregnant females. All samples were collected with approval from a Health Research Authority Research Ethics Committee (approval numbers 11/WA/0040 and 13/WA/019). All participants provided informed consent and the study adhered to the principles of the Declaration of Helsinki.

### Oxysterol Analysis

Oxysterol analysis was performed as described in detail in (41-44) with minor modifications. In brief, oxysterols were extracted in acetonitrile (absolute ethanol in the case of placental tissue) and isolated by solid phase extraction (SPE). The extract was split into two equal fractions -A and -B. Cholesterol oxidase was added to fraction-A to convert endogenous 3β-hydroxy-5-ene oxysterols to the 3-oxo-4- ene equivalents which were then derivatised with [^2^H_5_]GP (Supplemental Figure S1). Fraction-B was treated with [^2^H_0_]GP in the *absence of cholesterol oxidase*, thus giving a measure of endogenous 3- oxo-4-ene oxysterols while Fraction-A gives a measure of endogenous 3β-hydroxy-5-ene plus 3-oxo- 4-ene oxysterols. Fractions -A and -B were combined and analysed by LC-MS with multi-stage fragmentation (MS^3^) i.e. [M]^+^→[M-Py]^+^→ on a Orbitrap Elite high resolution mass spectrometer (ThermoFisher Scientific). In the MS mode resolution was 120,000 FWHM at *m/z* 400 with mass accuracy better than 5 ppm. MS^3^ spectra were recorded in the linear ion trap (LIT) in parallel to acquisition of mass spectra in the Orbitrap. Oxysterols were identified by reference to authentic standards, unless stated otherwise. Quantification was achieved with the isotope-labelled standards [25,26,26,26,27,27,27-^2^H_7_]24R/S-HC and [25,26,26,26,27,27,27-^2^H_7_]22R-hydroxycholest-4-en-3-one ([^2^H_7_]22R-HCO) which have been shown to be suitable for quantification of side-chain oxysterols (40, 43). Further details of the experimental methods can be found in Supplemental Information.

### Transfection studies

HSD3B1 transfection was performed using the pcDNA3-HSD3B1-STOP plasmid. The plasmid construct was transfected into HEK293 cells using JetOPTIMUS© transfection reagent. Plasmid DNA was combined with JetOPTIMUS buffer at 1 μL per 10 ng of DNA and vortexed briefly. The JetOPTIMUS transfection reagent was added at 1 μL per 1000 ng of plasmid DNA and mixed gently. The mixture was incubated at room temperature for 10 min before pipetting evenly into seeded 60 mm dishes. The cells were incubated for 24 hr at 37°C and 5% CO_2_ to allow for transfection and *in vitro* expression of the HSD3B1 enzyme from the HSD3B1 open reading frame (ORF) in the plasmid.

Transfected HEK293 cells were treated with oxysterols. The incubation buffer was potassium phosphate, pH 6.8 containing 1 mM EDTA, 1 mM of NAD^+^ and 1 μM of oxysterol. 1 mL of oxysterol incubation buffer was added to each dish. The cells were incubated for 1 hr at 37°C, 5% CO_2_. An aliquot of cells was taken for immunoblot while a separate aliquot was taken for LC-MS(MS^3^) analysis.

## Results

### Plasma from pregnant women, plasma from umbilical cord blood and placental tissue contains 3-oxocholest-4-en-(25R)26-oic acid and (25R)26-hydroxycholest-4-en-3-one

The sterol-acid 3O-CA has been identified previously at low levels compared to 3β-HCA in human plasma. Its origin is unknown. As part of an investigation into oxysterols, including sterol-acids, associated with human pregnancy EADSA technology was employed to identify oxysterols based on their 3β-hydroxy-5-ene scaffold or native oxo group. As anticipated, 3O-CA was found in blood from pregnant women taken 1 – 2 days before elective caesarean section, but at quite appreciable levels (2.41 ± 0.61 ng/mL ± SD), corresponding to about 6% of that of 3β-HCA (Figure 2C-D, Table 1, see also Supplemental Figure S2B). In terms of absolute concentration 3O-CA was found at similar levels in plasma from non-pregnant women (1.99 ± 0.69 ng/mL), but when compared to 3β-HCA at a level of only 2% (Figure 2A – B, Supplemental Figure S2A). Cord blood is blood left over in the placenta after birth and collected from the umbilical cord, the level of 3O-CA in cord plasma (12.74 ± 6.60 ng/mL) is higher than in circulating plasma from both pregnant and non-pregnant women, and similar to that of 3β-HCA in cord plasma (Figure 2E – F, Supplemental Figure S2C). This data suggests that 3O-CA may be produced from 3β-HCA in the placenta, where the levels of 3O-CA (3.99 ± 1.29 ng/g) are about the same as 3β-HCA (Figure 2G – H, Supplemental Figure S2D). Note concentrations of 3β-HCA are reported in (44).

**Table 1.** 3β-Hydroxy-5-ene and 3-oxo-4-ene sterols in plasma from non-pregnant and pregnant women and the umbilical cord.

| [<sup>2</sup>H<sub>0</sub>]GP | [<sup>2</sup>H<sub>5</sub>]GP | COMPOUND | CORD PLASMA (n = 14) |  |  | PREGNANT WOMAN PLASMA (n = 10) |  |  | CONTROL PLASMA (n = 5) |  |  | NOTE |
| --- | --- | --- | --- | --- | --- | --- | --- | --- | --- | --- | --- | --- |
| m/z | m/z | ABBREVIATION | MEAN (ng/mL) | SD | 3-ONE/3β-OL | MEAN (ng/mL) | SD | 3-ONE/3β-OL | MEAN (ng/mL) | SD | 3-ONE/3β-OL | |
|  | 539.4368 | 22R-HC | 6.19 | 3.01 |  | 2.55 | 1.18 |  | 0.00 | 0.00 |  |  |
| 534.4054 |  | 22R-HCO | 0.11 | 0.21 | 0.02 | 0.00 | 0.00 | 0.00 | 0.00 | 0.00 | 0.00 |  |
|  | 539.4368 | 24S-HC | 7.34 | 2.38 |  | 14.99 | 4.28 |  | 14.48 | 3.33 |  |  |
| 534.4054 |  | 24S-HCO | 0.29 | 0.39 | 0.04 | 0.04 | 0.10 | 0.00 | 0.00 | 0.00 | 0.00 |  |
|  | 539.4368 | 25-HC | 2.75 | 1.88 |  | 2.74 | 1.66 |  | 1.55 | 0.13 |  |  |
| 534.4054 |  | 25-HCO | 0.00 | 0.00 | 0.00 | 0.00 | 0.00 | 0.00 | 0.00 | 0.00 | 0.00 | 1 |
|  | 539.4368 | 26-HC | 7.23 | 2.88 |  | 21.61 | 5.63 |  | 25.67 | 2.30 |  |  |
| 534.4054 |  | 26-HCO | 1.65 | 0.68 | 0.23 | 0.69 | 0.42 | 0.03 | 0.00 | 0.00 | 0.00 |  |
| | 555.43172 | 7α,25-diHC | 0.48 | 0.48 | | 1.07 | 0.35 | | 0.79 | 0.64 | | |
| 550.40032 | | 7α,25-diHCO | 0.72 | 0.88 | 1.50 | 1.77 | 0.76 | 1.65 | 1.40 | 0.43 | 1.77 | |
| | 555.43172 | 7α,26-diHC | 0.49 | 0.23 | | 1.03 | 0.54 | | 1.08 | 0.55 | | |
| 550.40032 | | 7α,26-diHCO | 3.24 | 1.26 | 6.56 | 3.89 | 1.19 | 3.79 | 4.13 | 1.23 | 3.82 | |
|  | 555.43172 | 20R,22R-diHC | 49.74 | 23.97 |  | 13.23 | 6.79 |  | 0.00 | 0.00 |  |  |
| 550.40032 |  | 20R,22R-diHCO | 12.86 | 6.09 | 0.26 | 6.04 | 3.94 | 0.46 | 0.00 | 0.00 | 0.00 |  |
| | 553.41607 | 3β-HCA | 10.56 | 4.71 | | 43.64 | 13.71 | | 90.02 | 21.40 | | |
| 548.38467 |  | 3O-CA | 12.74 | 6.60 | 1.21 | 2.41 | 0.61 | 0.06 | 1.99 | 0.69 | 0.02 |  |
| | 511.3691 | 3βH-$\Delta^5$-BA | 3.37 | 2.64 | | 1.50 | 0.81 | | 1.89 | 0.65 | | |
| 506.3377 | | 3O-$\Delta^4$-BA | 0.76 | 0.47 | 0.23 | 0.00 | 0.00 | 0.00 | 0.00 | 0.00 | 0.00 | |
| | 553.41607 | 3β,20-diHC-22O | 103.60 | 44.51 | | 21.72 | 8.22 | | 10.59 | 5.77 | | 2,3 |
| 548.38467 |  | 20-HC-3,22-diO | 78.24 | 30.85 | 0.76 | 18.21 | 6.91 | 0.84 | 0.00 | 0.00 | 0.00 | 2,4 |
| | 569.41098 | 3β,25-diHCA | 13.02 | 6.47 | | 6.34 | 2.03 | | 6.36 | 3.05 | | 2,5 |
| 564.37958 |  | 25H,3O-CA | 12.12 | 5.03 | 0.93 | 4.29 | 2.64 | 0.68 | 0.00 | 0.00 | 0.00 | 2,6 |
| | 569.41098 | 3β,x-diHCA | 34.00 | 16.87 | | 7.42 | 2.95 | | 4.68 | 1.82 | | 2,7 |
| 564.37958 |  | xH,3O-CA | 11.80 | 4.60 | 0.35 | 4.99 | 2.25 | 0.67 | 0.00 | 0.00 |  | 2,8 |

Data for 22R-HC, 24S-HC, 26-HC, 7α ,26-diHC and 3β-HCA reported in [doi: https://doi.org/10.1101/2022.02.07.478301].

**Figure 2.**
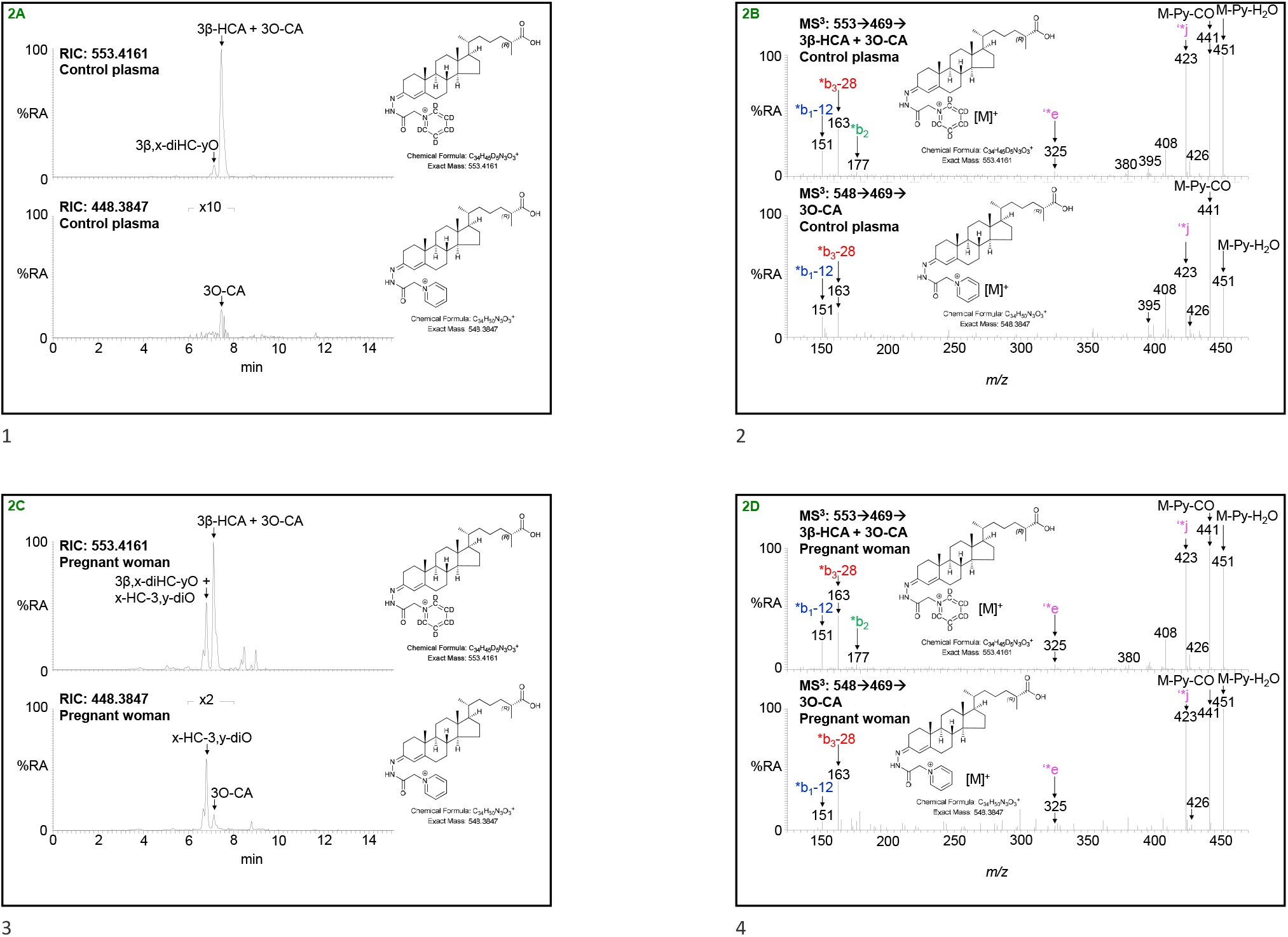
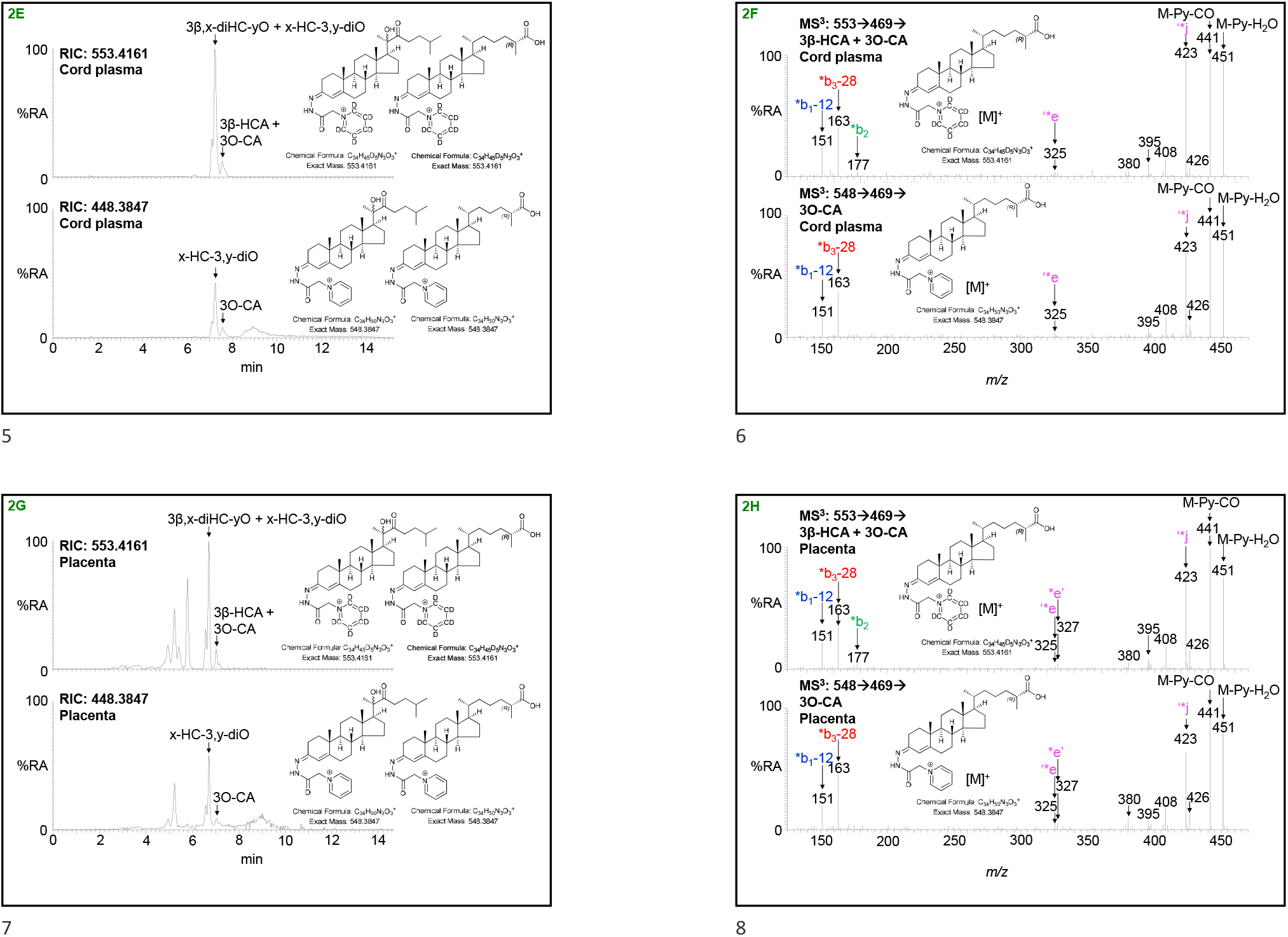
3β-HCA and 3O-CA are present in plasma from non-pregnant (control) and pregnant women, umbilical cord plasma and in placental tissue. Reconstructed ion chromatograms (RICs) of 553.4161 ± 5 ppm corresponding to 3β-HCA plus 3O-CA derivatised with [^2^H_5_]GP (upper panels) and 548.3847 ± 5 ppm corresponding to 3O-CA derivatised with [^2^H_0_]GP (lower panels). (A) Non-pregnant woman’s (control) plasma, (C) pregnant woman’s plasma, (E) cord plasma, and (G) placenta. Chromatograms in upper and lower panels are plotted on the same y-axis and magnified as indicated. MS^3^ ([M]^+^→[M-Py]^+^→) spectra of 3β-HCA plus 3O-CA derivatised with [^2^H_5_]GP (upper panels) and 3O-CA derivatised with [^2^H_0_]GP (lower panels) from (B) non-pregnant woman’s (control) plasma, (D) pregnant woman’s plasma, (F) cord plasma, and (H) placenta. There is some shift in retention time between samples which were analysed at different times on different LC columns but of the same type. MS^3^ spectra can be compared to those of authentic standards (43). Further data can be found in Supplemental Figure S2A – D.

3O-CA may be derived by oxidation of 3β-HCA at C-3 or conceivably by oxidation of 26-HCO at C-26 to yield the carboxylic acid group (Figure 3). Although to the best of our knowledge concentrations of 26-HCO have not been reported in plasma, we have seen evidence in previous studies for the presence of 26-HCO at about the level of detection but below the level of quantification (<0.2 ng/mL). By investigating the reconstructed ion chromatogram (RIC) at *m/z* 534.4054 ± 5 ppm appropriate to 26-HCO following [^2^H_0_]GP-derivatisation, a peak with the correct retention time is evident in plasma from pregnant women and co-eluting with that of 26-HC following *ex vivo* cholesterol oxidase treatment and [^2^H_5_]GP-derivatisation at *m/z* 539.4368 ± 5 ppm (Figure 4C, see also Supplemental Figure S2F). The MS^3^ ([M]^+^→[M-Py]^+^→) fragmentation spectra of GP-derivatised 26-HCO and 26-HC are identical, with the exception of the precursor-ion *m/z*, confirming the identification endogenous 26-HCO (Figure 4D, see Yutuc et al (43) for MS^3^ spectra of reference standards). The level of 26-HCO in pregnant women’s plasma is low at 0.69 ± 0.42 ng/mL and was only detected in nine of the ten samples analysed, being present at 3% that of 26-HC. In plasma from non-pregnant females, 26-HCO was only just detectable (Figure 4A - B) but was below the limit of quantification (0.2 ng/mL) in all samples analysed. The situation in cord plasma is quite different (Figure 4E - F), the level of 26-HCO was found to be 1.65 ± 0.68 ng/mL, 23% that of 26-HC. As was the case in placenta for 3O-CA and 3β-HCA, the ratio of 26-HCO to 26-HC in this tissue was also high, with 26-HCO at 16.0 ± 3.39 ng/mg being 39% that of 26-HC (Figure 4G -H).

**Figure 3.**
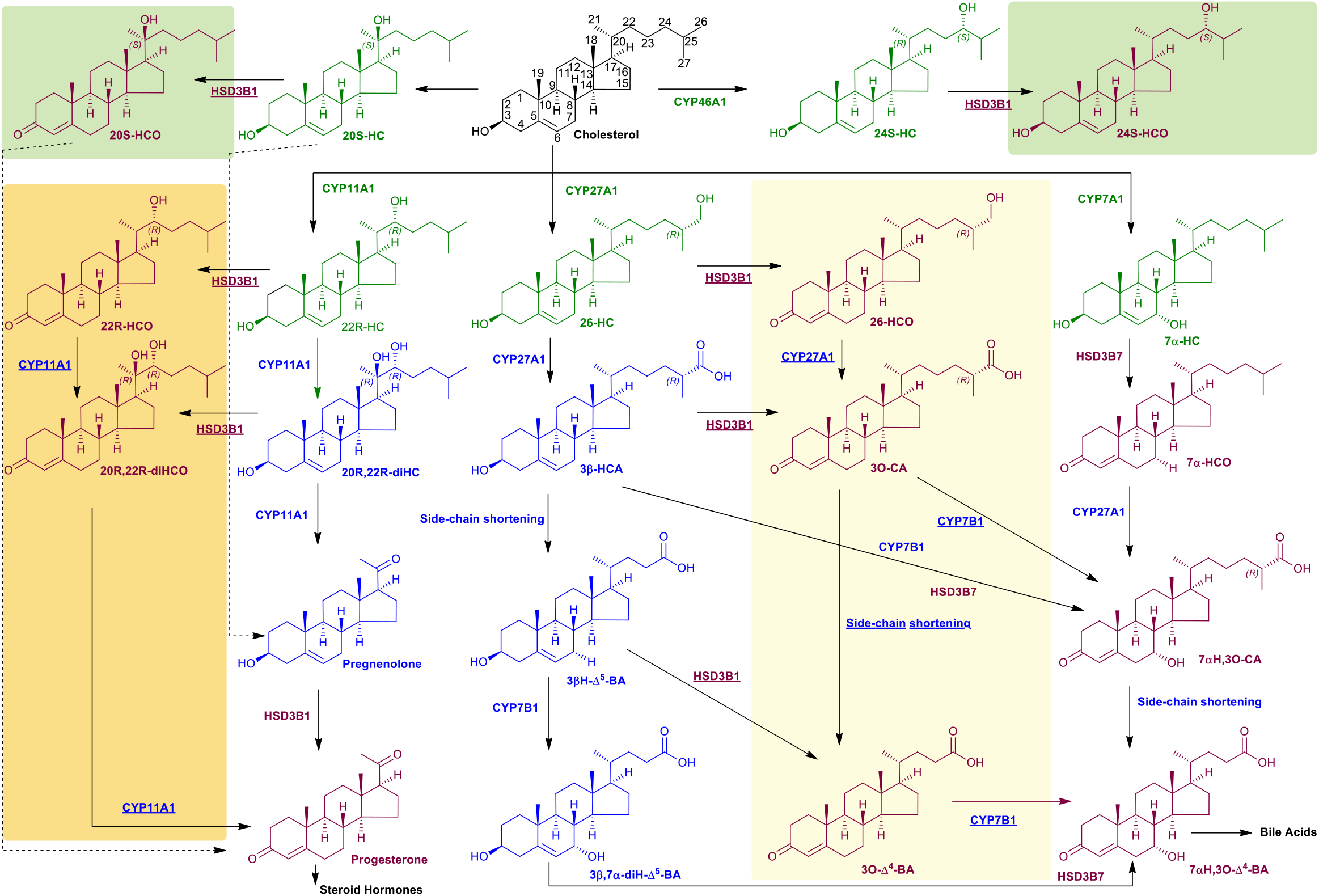
Suggested metabolic pathways generating oxysterols with a 3-oxo-4-ene function. Unexpected pathways are shown on pale yellow, orange and lime backgrounds. Oxysterols and steroids are coloured as in Figure 1.

**Figure 4.**
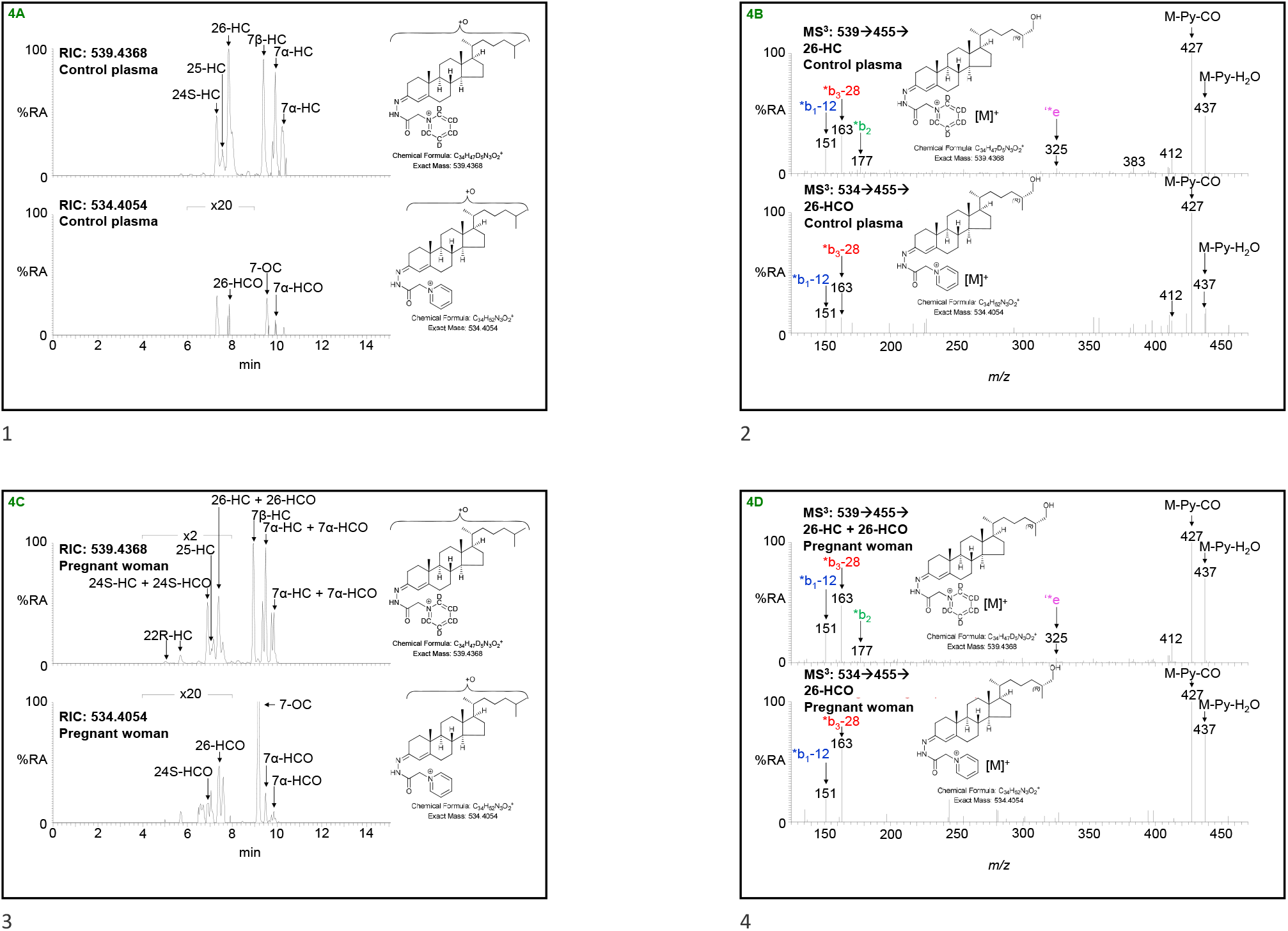
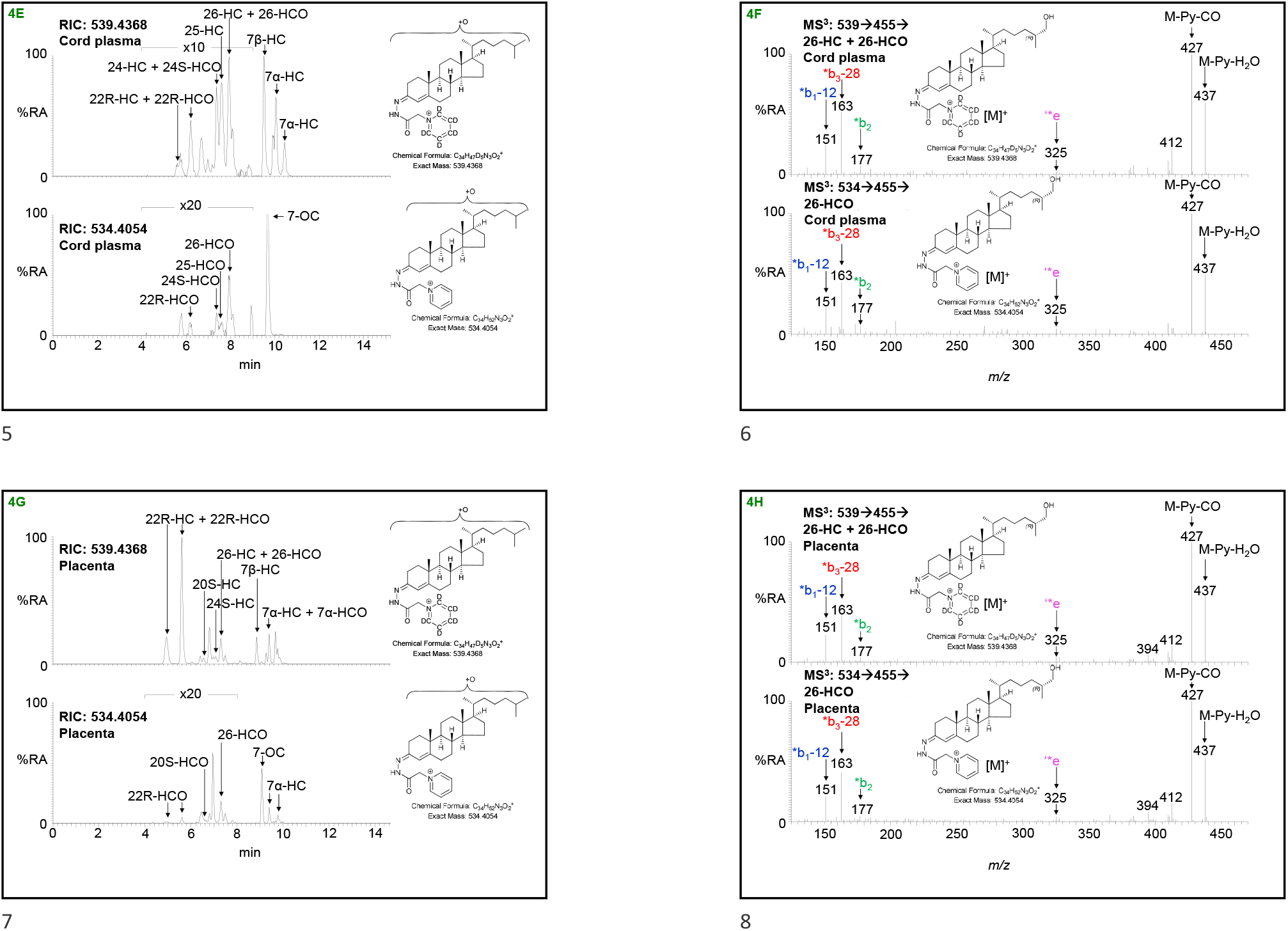
Monohydroxycholesterols (HC) and monohydroxycholestenones (HCO) are present in plasma from non-pregnant (control) and pregnant women, umbilical cord plasma and in placental tissue. RICs of 539.4368 ± 5 ppm corresponding to monohydroxycholesterols plus monohydroxycholestenones derivatised with [^2^H_5_]GP (upper panels) and 534.4054 ± 5 ppm corresponding to monohydroxycholestenones derivatised with [^2^H_0_]GP (lower panels). (A) Non-pregnant woman’s (control) plasma, (C) pregnant woman’s plasma, (E) cord plasma, and (G) placenta. Chromatograms in upper and lower panels are plotted on the same y-axis and magnified as indicated. MS^3^ ([M]^+^→[M-Py]^+^→) spectra of 26-HC plus 26-HCO derivatised with [^2^H_5_]GP (upper panels) and 26-HCO derivatised with [^2^H_0_]GP (lower panels) from (B) non-pregnant woman’s (control) plasma, (D) pregnant woman’s plasma, (F) cord plasma, and (H) placenta. There is some shift in retention time between samples which were analysed at different times on different LC columns but of the same type. MS^3^ spectra can be compared to those of authentic standards (43). Further data can be found in Supplemental Figure S2E - H.

### Potential artefactual formation of 3-oxo-4-ene oxysterols from 3β-hydroxy-5-ene oxysterols

An alternative explanation for the current data showing the existence of 3-oxo-4-ene oxysterols formed in the absence of a 7α-hydroxy group is that their defining LC-MS peaks may be artefacts generated as a consequence of the presence of contaminating [^2^H_0_]GP within the [^2^H_5_]GP reagent which is used in conjunction with cholesterol oxidase enzyme in the EADSA process. Such a situation would result in a fraction of 3β-hydroxy-5-ene-sterols becoming derivatised with [^2^H_0_]GP instead of [^2^H_5_]GP and being incorrectly interpreted as being derived from endogenous 3-oxo-4-ene oxysterols. This situation is unlikely to be significant as the [^2^H_5_]pyridine from which [^2^H_5_]GP reagent is prepared is 99.94% isotopically pure, which should lead to no more than 0.06% artefactual formation of the [^2^H_0_]GP derivatives of 26-HCO or 3O-CA derived from native 26-HC and 3β-HCA, respectively. The levels of 26-HCO in pregnant women’s and cord plasma are 3% and 23% of 26-HC, respectively, and in placenta 26-HCO is 39% of 26-HC, values very much great than 0.06%. Another possibility whereby unreliable data can be generated is through back-exchange of the derivatisation groups when fractions-A ([^2^H_5_]GP) and fraction-B ([^2^H_0_]GP) are mixed just prior to LC-MS analysis. However, in the absence of an acid catalyst the exchange reaction does not proceed. To confirm that the observation of unexpected 3-oxo-4-ene oxysterols were not *ex vivo* artifacts of sample preparation, including the EADSA process or of sample injection, the formation of [^2^H_7_]24R/S-hydroxycholest-4-en-3-one ([^2^H_7_]24R/S-HCO) from [^2^H_7_]24R/S-hydroxycholesterol ([^2^H_7_]24R/S-HC) internal standard, added during the first step of oxysterol extraction, was monitored for every sample analysed. In no case was the [^2^H_0_]GP derivative of [^2^H_7_]24R/S-HCO observed (Supplemental Figure S3), eliminating the possibility of *ex vivo* formation of 3-oxo-4-ene oxysterols and confirming the high isotopic purity of [^2^H_5_]GP-hydazine.

### Metabolic products of 3O-CA

The immediate metabolic product of 3β-HCA in bile acid biosynthesis is 3β,7α-diHCA, formed in a reaction catalysed by CYP7B1 (Figure 1). 3β,7α-diHCA is then converted by HSD3B7 to 7αH,3O-CA, which then undergoes peroxisomal side-chain shortening to give the C_24_ acid 7α-hydroxy-3-oxochol-4-en-24-oic acid (7αH,3O-Δ^4^-BA); or 7αH,3O-CA may be first reduced in the A-ring by aldoketoreductase (AKR) 1D1 then by AKR1C4 prior to peroxisomal side-chain shortening to ultimately give chenodeoxycholic acid (5, 6). Alternatively, 3β,7α-diHCA itself can undergo side-chain shortening to give 3β,7α-dihydroxychol-5-en-24-oic acid (3β,7α-diH-Δ^5^-BA). Each of these pathways proceeds following 7α-hydroxylation of the core 3β-hydroxy-5-ene sterol. It is unknown whether CYP7B1 will accept 3-oxo-4-ene oxysterols and sterol-acids as substrates (Figure 3), and whether 3O-CA will undergo side-chain shortening leading to 3-oxochol-4-en-24-oic acid (3O-Δ^4^-BA). Interrogation of the RIC at *m/z* 506.3377 ± 5 ppm corresponding to [^2^H_0_]GP derivatised 3O-Δ^4^-BA indicates its presence in plasma from pregnant women but below the limit of quantification (0.2 ng/mL, Figure 5C - D, Supplemental Figure S4B). In cord plasma the concentration of 3O-Δ^4^-BA is still low (0.76 ± 0.47 ng/mL) but about 23% that of 3β-hydroxychol-5-en-24-oic acid (3βH-Δ^5^-BA, Figure 5E - F). In placenta the concentration of 3O-Δ^4^-BA is also low at 1.26 ± 0.72 ng/mg and about 16% that of 3βH-Δ^5^-BA (Figure 5G - H). 3O-Δ^4^-BA was not observed in plasma from non-pregnant women (Figure 5A).

**Figure 5.**
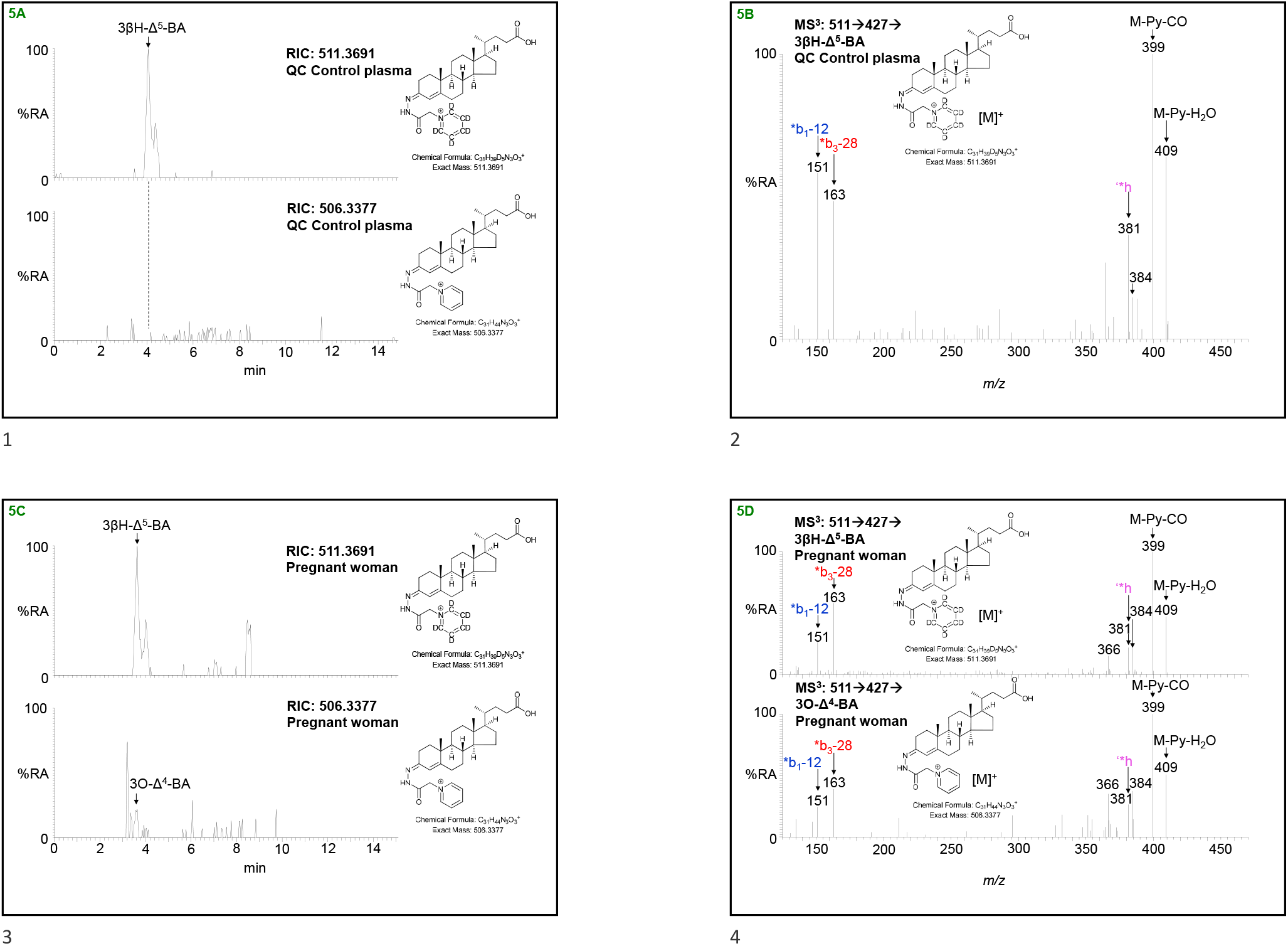
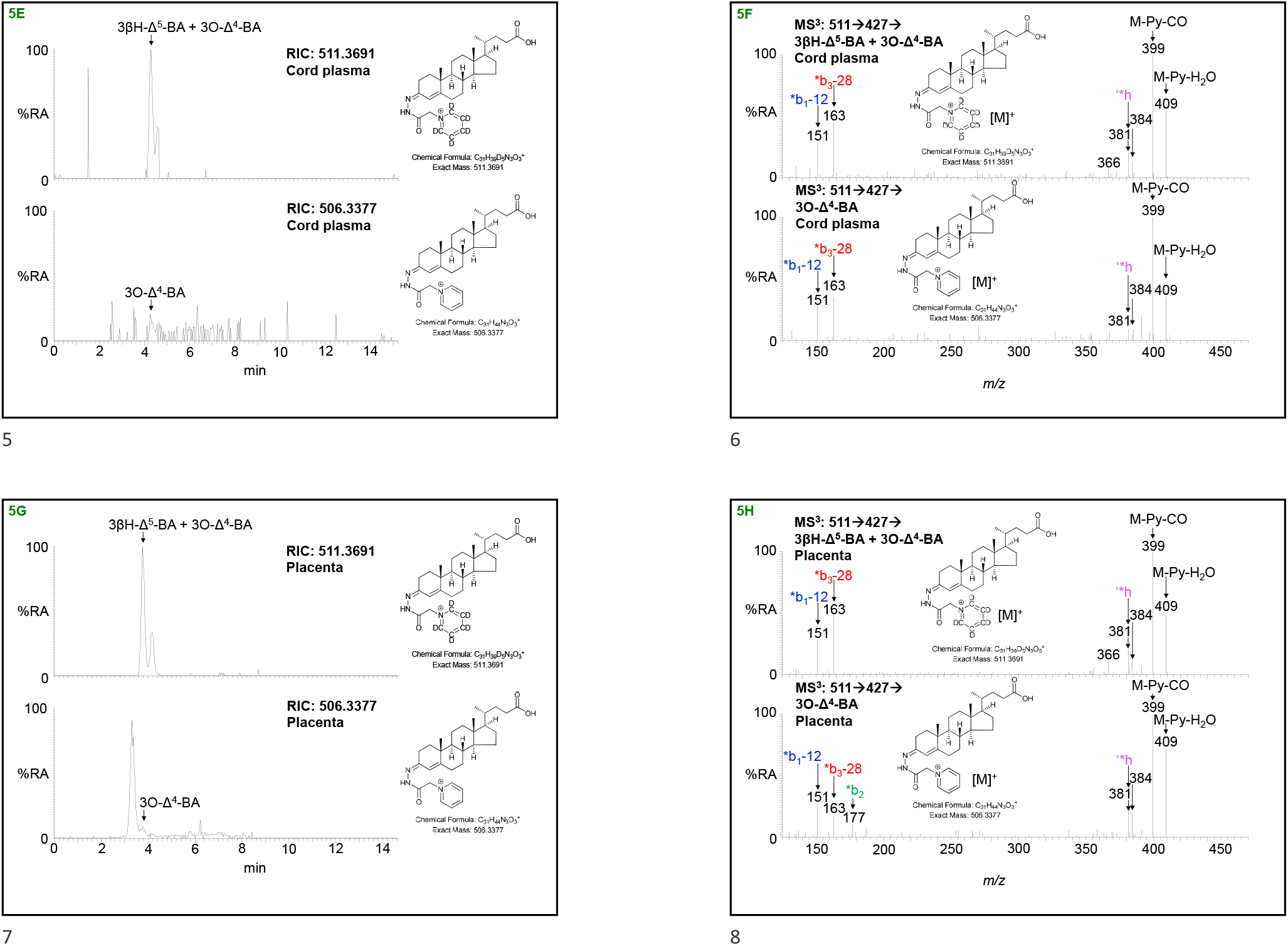
3-Oxo-4-ene bile acids in pregnant women’s and cord plasma and in placental tissue. RICs of 511.3691 ± 5 ppm corresponding to 3βH-Δ^5^-BA plus 3O-Δ^4^-BA (where present) derivatised with [^2^H_5_]GP (upper panels) and 506.3377 ± 5 ppm corresponding to 3O-Δ^4^-BA derivatised with [^2^H_0_]GP (lower panels). (A) Non-pregnant woman’s (control) plasma, (C) pregnant woman’s plasma, (E) cord plasma, and (G) placenta. Chromatograms in upper and lower panels are plotted on the same y-axis and magnified as indicated. MS^3^ ([M]^+^→[M-Py]^+^→) spectra of 3βH-Δ^5^-BA plus 3O-Δ^4^-BA (if present) derivatised with [^2^H_5_]GP (upper panels) and where present 3O-Δ^4^-BA derivatised with [^2^H_0_]GP (lower panels) from (B) non-pregnant woman’s (control) plasma, (D) pregnant woman’s plasma, (F) cord plasma, and (H) placenta. There is some shift in retention time between samples which were analysed at different times and on different LC columns but of the same type. MS^3^ spectra can be compared to those of authentic standards (43).

The current data suggests that 3-oxo-4-ene sterol acids may be formed in placenta even when they do not possess a 7α-hydroxy group. A pathway can be envisaged starting with 26-HC and involving the intermediates 26-HCO, 3O-CA and 3O-Δ^4^-BA (shown on a pale-yellow background in Figure 3). Alternatively, like 26-HCO, 3O-CA and 3O-Δ^4^-BA could be formed directly from their 3β-hydroxy-5-ene equivalents by a hydroxysteroid dehydrogenase enzyme other than HSD3B7.

### Placenta and umbilical cord blood contain novel intermediates in a pathway leading to progesterone

Progesterone is formed in steroidogenic tissue from cholesterol by mitochondrial CYP11A1 and HSD3B1/2 (Figure 1) (21), CYP11A1 and HSD3B1 are both enriched in placenta (9, 19). In the first step, cholesterol is converted to 22R-HC and then further to 20R,22R-diHC and onto pregnenolone all by CYP11A1 (22, 23). Pregnenolone is then converted to progesterone by HSD3B1 in placenta and by HSD3B2 in other steroidogenic tissues. The evidence presented above suggesting that 26-HC can be converted to 26-HCO in placenta raises the possibility that similar oxidation reactions may proceed with 22R-HC and 20R,22R-diHC as substrates leading to 22R-hydroxycholest-4-en-3-one (22R-HCO) and 20R,22R-dihydroxycholest-4-en-3-one (20R,22R-diHCO) products, providing an alternative route to progesterone involving the same enzymes as the conventional route but acting in a different order and avoiding pregnenolone (Figure 3, see pathway on an orange background).

22R-HCO was found to be present in only 3 out of the 14 cord plasma samples, where it was found, its level was about 0.5 ng/mL, for comparison 22R-HC was found in all samples at a level of 6.19 ± 3.01 ng/mL (Figures 4E & 6B, Supplemental Figure S2G) (44). 22R-HCO was at or below the detection limit in plasma from pregnant and non-pregnant women (control) plasma, although 22R-HC was found in pregnant women’s plasma (2.55 ± 1.18 ng/mL, Figures 4A, 4C & 6A (44)). In placenta the concentration of 22R-HCO was found to be 2.16 ± 0.38 ng/mL, 1% that of 22R-HC (Figure 4G & 6C).

**Figure 6.**
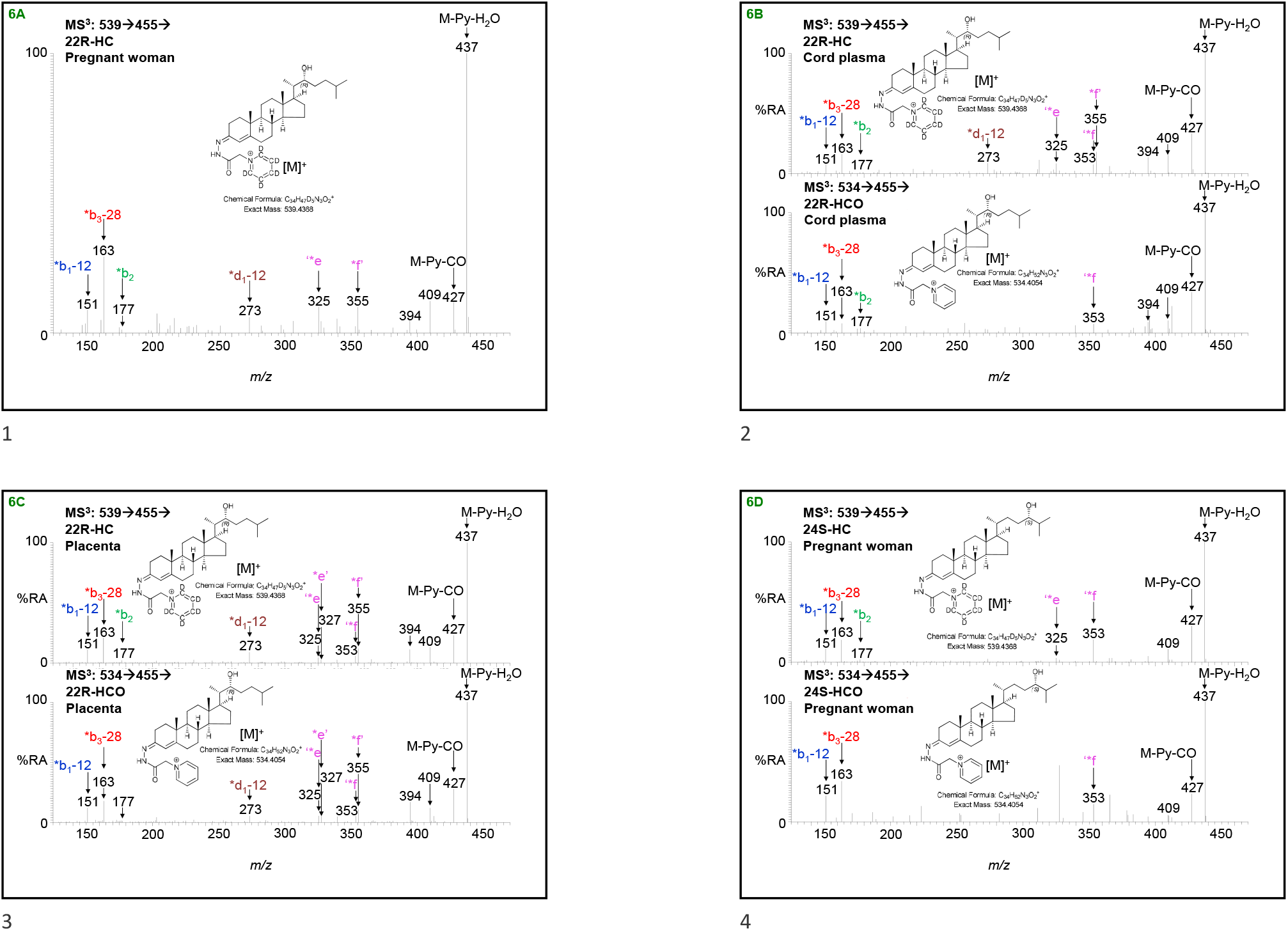
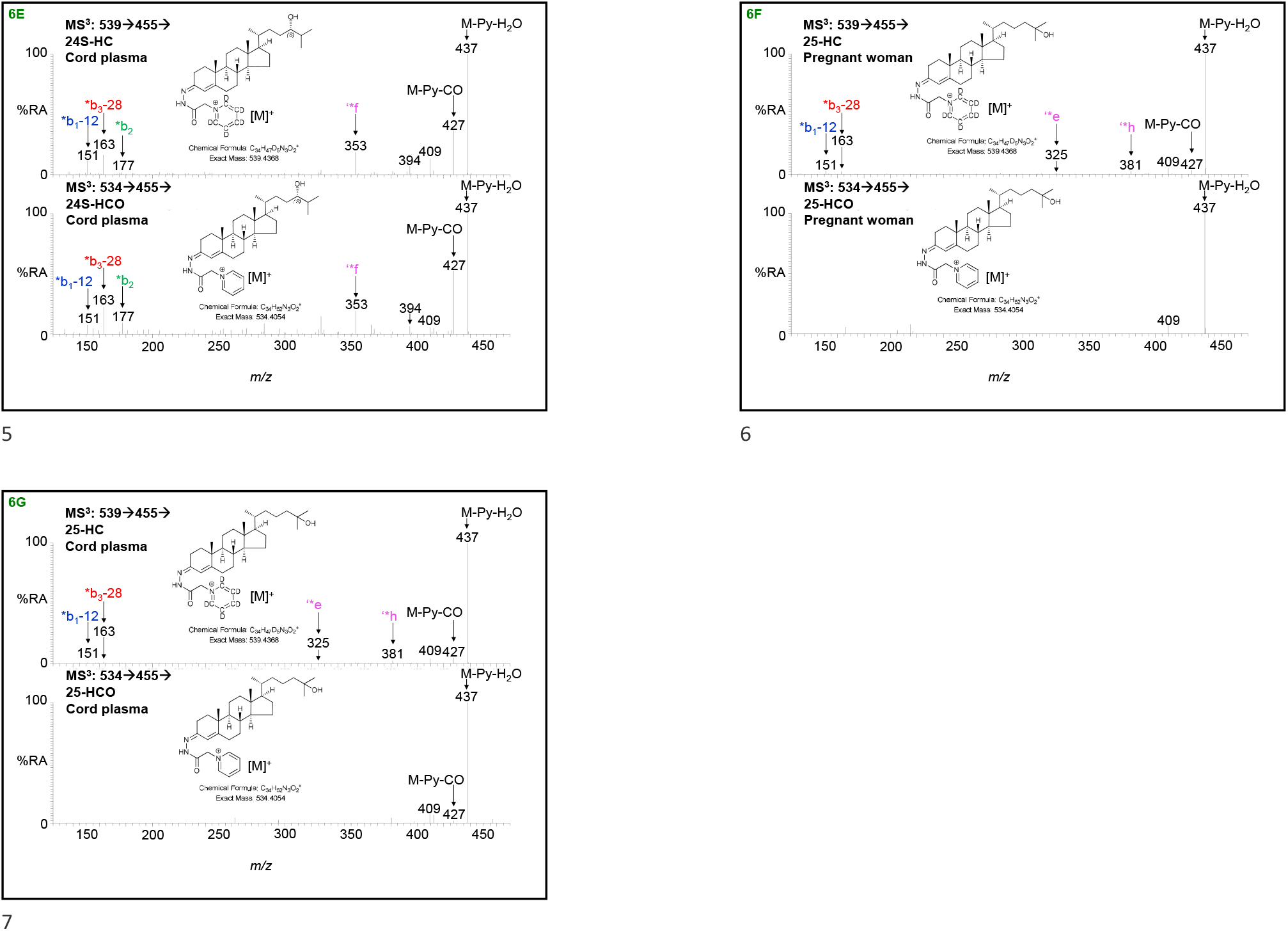
Hydroxycholestenones are present in pregnant women’s and cord plasma and in placenta tissue. MS^3^ ([M]^+^→[M-Py]^+^→) spectra of hydroxycholesterol (HC) plus hydroxycholestenone (HCO, where present) derivatised with [^2^H_5_]GP (upper panels) and hydroxycholestenones (if present) derivatised with [^2^H_0_]GP (lower panels). (A) 22R-HC is present in pregnant woman’s and (B) cord plasma but 22R-HCO is at or below the detection limit in these plasmas, while it is present in (C) placenta. 24S-HCO is present in (D) pregnant woman’s and (E) cord plasma. 25-HCO is at the detection limit in (F) pregnant woman’s plasma and (G) cord plasma. See Figure 8D & 8E for spectra of 24S-HCO and 25-HCO in placenta. Note, the presence of the fragment ion at *m/z* 327 in the MS^3^ spectra of 24S-HCO from (6D) pregnant woman’s and (6E) cord plasma indicating the additional presence of minor amounts of 20S-HCO. See Figure 4C, 4E and 4G for chromatograms from pregnant woman’s and cord plasma, and placenta, respectively. There is some shift in retention time between samples which were analysed at different times and on different LC columns but of the same type. MS^3^ spectra can be compared to those of authentic standards (43).

20R,22R-HCO was found in each of the cord plasma samples analysed, it was much more abundant than 22R-HCO, at a concentration of 12.86 ± 6.09 ng/mL corresponding to 26% that of 20R,22R-diHC (Figure 7C & 7F, Supplemental Figure S5C). In plasma from pregnant women 20R,22R-diHCO was present at a concentration of 6.04 ± 3.94 ng/mL, 46 % that of 20R,22R-diHC (Figure 7B & 7E), although it, and 20R,22R-diHC, were absent from plasma from non-pregnant females (Figure 7A). Both 20R,22R- diHCO and 20R,22R-diHC were present in placenta (Figure 7D & 7G).

**Figure 7.**
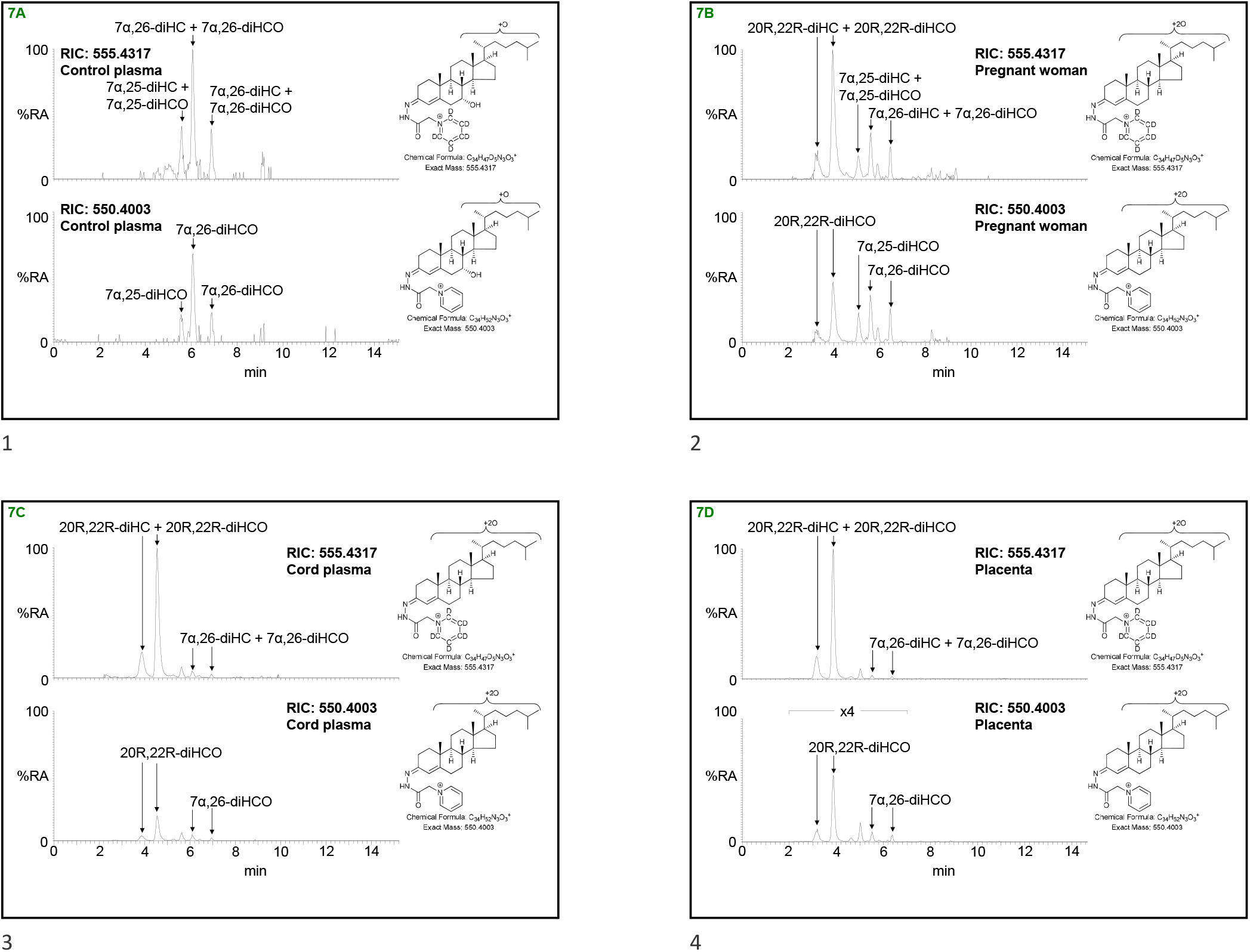
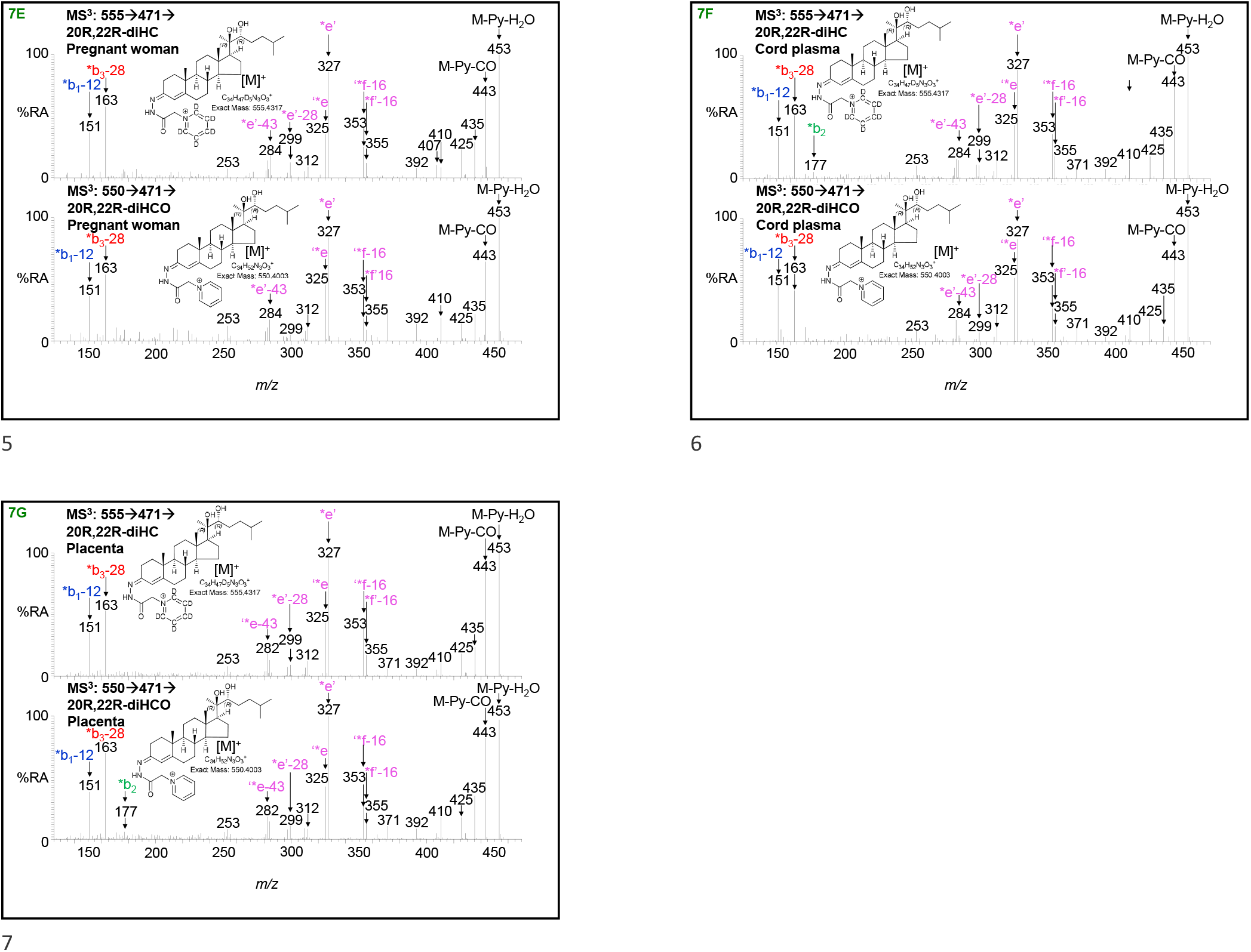
20R,22R-diHCO is present in pregnant women’s and cord plasma and in placental tissue. RICs of 555.4317 ± 5 ppm corresponding to dihydroxycholesterols (diHC) plus dihydroxycholestenones (diHCO) including 20R,22R-diHC plus 20R,22R-diHCO when present, derivatised with [^2^H_5_]GP (upper panels) and 550.4003 ± 5 ppm corresponding to dihydroxycholestenones, including 20R,22R-diHCO if present, derivatised with [^2^H_0_]GP (lower panels). (A) Non-pregnant female (control) plasma, (B) pregnant woman’s plasma, (C) cord plasma, and (D) placenta. Chromatograms in upper and lower panels are plotted on the same y-axis and magnified as indicated. MS^3^ ([M]^+^→[M-Py]^+^→) spectra of 20R,22R-diHC plus 20R,22R-diHCO derivatised with [^2^H_5_]GP (upper panels) and 20R,22R-diHCO derivatised with [^2^H_0_]GP (lower panels) from (E) pregnant woman’s plasma, (F) cord plasma, and (G) placenta. There is some shift in retention time between samples which were analysed at different times and on different LC columns but of the same type. MS^3^ spectra can be compared to those of authentic standards (43). Further data can be found in Supplemental Figure S5A - D.

It is surprising that the ratio of 20R,22R-diHCO to 20R,22R-diHC is greater in plasma from pregnant women than cord plasma, but the fact that both these sterols are more abundant than their 20R-monohydroxy analogues raise the possibility of further metabolism of 20R,22R-diHCO by CYP11A1 to progesterone in the placenta or alternatively to intermediates analogous to a recently described pathway to bile acids (44).

### 24S-Hydroxycholest-4-en-3-one is present in pregnant women’s and cord plasma, 20S-hydroxycholest-4-en-3-one is present in placenta

In plasma from pregnant women, 24S-HC is the second most abundant side-chain monohydroxycholesterol after 26-HC (Figure 4C & 6D, Table 1). While 26-HCO was present in nine of the ten samples analysed, at about 3% of 26-HC, 24S-hydroxycholest-4-en-3-one (24S-HCO) could only be detected in three of the ten samples, giving a mean concentration of 0.04 ± 0.10 ng/mL which is less than 1% of the concentration of 24S-HC. 24S-HCO was below the limit of quantification in all five plasma samples from non-pregnant females (Figure 4A). In cord plasma 24S-HCO was quantified in seven of the fourteen samples (Figure 4E & 6E), leading to a mean concentration of 0.29 ± 0.39 ng/mL, only 4% that of 24S-HC, compared to 26-HCO being 23% of 26-HC.

20S-HC is not normally found in plasma, but it is reported to be present in rodent brain and human placenta (44-46). Both 20S-HC and 24S-HC are found in placenta as are 20S-hydroxycholest-4-en-3-one (20S-HCO) and 24S-HCO (Figure 4G & 8A - 8D & Supplemental Figure S6). 20S-HC and 20S-HCO can be targeted by generating multiple reaction monitoring (MRM)-like chromatograms [M]^+^→[M-Py]^+^→327 (Figure 8B 1^st^ and 2^nd^ panels) while 24S-HC and 24S-HCO can be targeted by [M]^+^→[M-Py]→353 chromatograms (Figure 8B, 3^rd^ and 4^th^ panels). As is the case of most oxysterols, 20S-HC and 24S-HC and 20S-HCO and 24S-HCO each give twin peaks when derivatised with GP-hydrazine corresponding to *syn* and *anti* conformers about the hydrazone C=N double bond (43). While the first peaks for 20S-HC and 20S-HCO are resolved from 24S-HC and 24S-HCO peaks, and the second peaks for 24S-HC and 24S-HCO are similarly resolved from the 20S-isomers, the second 20S-HC and 20S-HCO peaks co-elute with the first peaks of 24S-HC and 24S-HCO (Figure 8A & B). This means that quantification in placenta is only approximate. As an estimation, 24S-HCO and 20S-HCO are present at about 5% of 24S-HC and 20S-HC, respectively.

**Figure 8.**
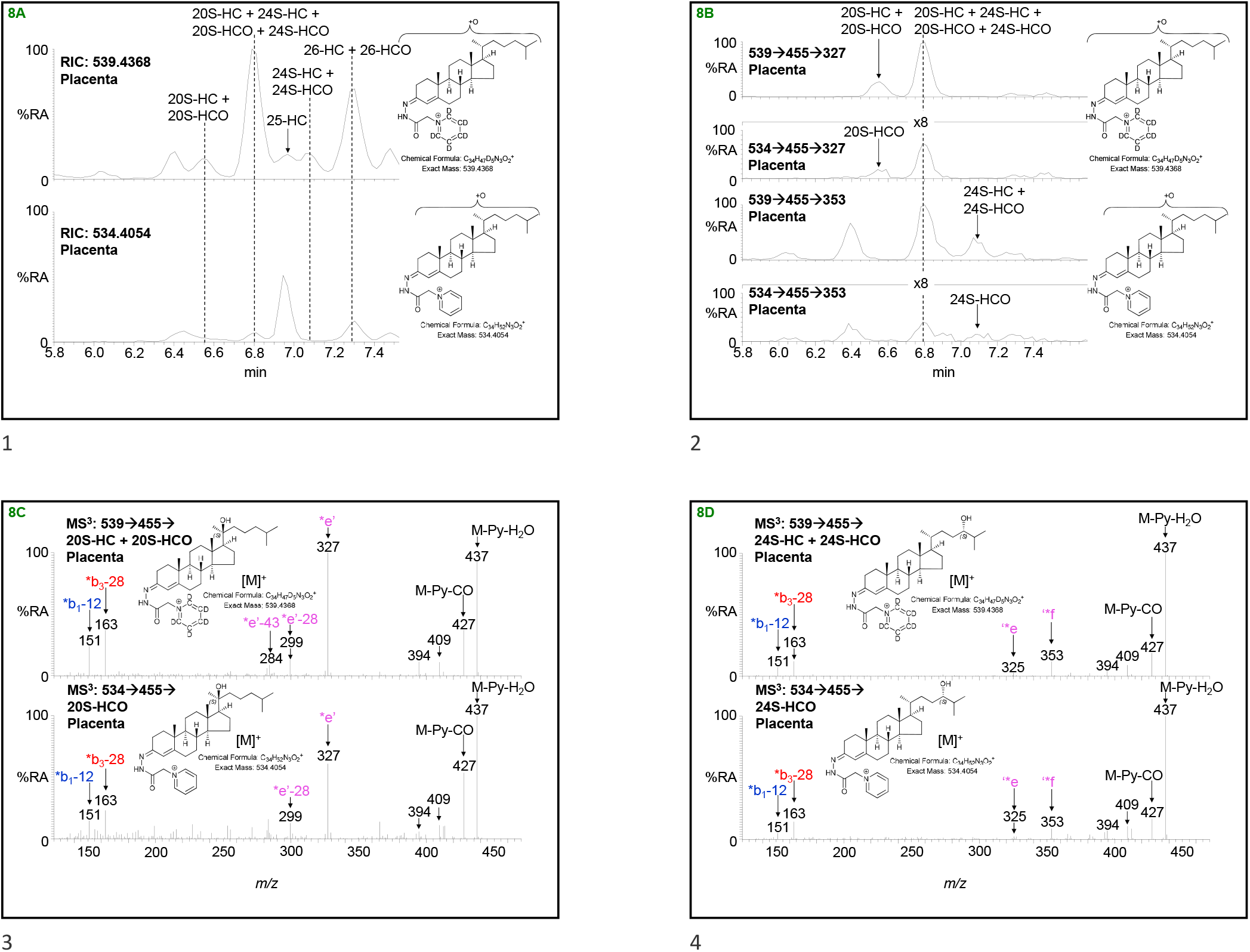
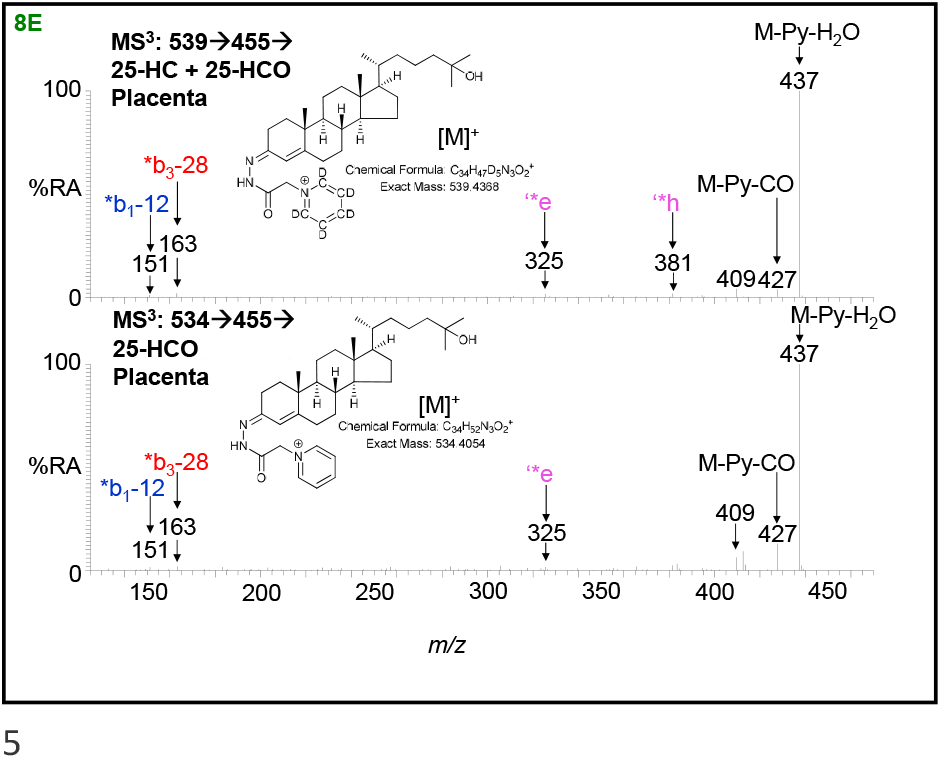
20S-HC, 20S-HCO, 24S-HC, 24S-HCO, 25-HC, 25-HCO, 26-HC and 26-HCO are present in placenta. (A) RICs of 539.4368 ± 5 ppm (upper panel) and 534.4054 ± 5 ppm (lower panel) over the elution time of 20S-HC, 20S-HCO, 24S-HC, 24S-HCO, 25-HC, 25-HCO, 26-HC and 26-HCO. The full-length chromatogram is presented in Figure 4G. (B) MRM-like chromatograms targeting 20S-HC (top panel) and 20S-HCO (2^nd^ panel) i.e. [M]^+^→[M-Py]^+^→327, plotted on the same y-axis, and 24S-HC (3^rd^ panel) and 24S-HCO (bottom pane) i.e. [M]^+^→[M-Py]→353, plotted on the same y-axis. MS^3^ ([M]^+^→[M-Py]^+^→) spectra of (C) the first peaks of 20S-HC plus 20S-HCO (upper panel) and 20S-HCO (lower panel), (D) the second peaks of 24S-HC plus 24S-HCO (upper panel) and 24S-HCO (lower panel), and (E) 25-HC plus 25-HCO (upper panel) and 25-HCO (lower panel). Note, the fragment ion at *m/z* 353 is characteristic of 24S-HC/24S-HCO while *m/z* 327 is characteristic of 20S-HC/20-HCO. Further data can be found in Supplemental Figure S6.

### Other 3-oxo-4-enes in plasma and placenta

Oxysterols with a 7α-hydroxy-3-oxo-4-ene structure are likely to be formed via the action of HSD3B7 on 3β,7α-dihydroxy-5-ene substrates (Figure 1) (8). Their identification and quantification in pregnant women’s and cord plasma is presented elsewhere (44).

In both pregnant women’s and cord plasma low levels of 25-HCO are detected but below the limit of quantification (Figures 4C & 4E, Supplemental Figure S2F & S2G). 25-HC the likely precursor is, however, present at low levels in these two types of plasma at about 2.75 ng/mL. 25-HC and 25-HCO are also found in placenta but were not quantified (Figure 8A & 8E).

In previous studies we have partially identified an oxysterol in plasma as either 3β,20- dihydroxycholest-5-en-22-one (3β,20-diHC-22O) or 3β,22-dihydroxycholest-5-en-24-one (3β,22-diHC-24O) but the identity was not confirmed in the absence of an authentic synthetic standards (43). This oxysterol, is present in the current samples generically named as 3β,x-diHC-yO (Figure 2 & Supplemental Figure S7) and its concentration increases in the order non-pregnant women’s plasma (10.59 ± 5.77 ng/mL, Figure 2A), pregnant women’s plasma (21.72 ± 8.22 ng/mL, Figure 2C) and cord plasma (103.6 ± 44.51 ng/mL, Figure 2E). It is also present in placenta (Figure 2G). Interestingly, the 3-oxo-4-ene version of this molecule (generically x-HC-3,y-diO), 20-hydroxycholest-4-ene-3,22-dione (20-HC-3,22-diO) or 22-hydroxycholest-4-ene-3,24-dione (22-HC-3,24-diO) shows the same trend in concentration, being below the limit of detection in non-pregnant female plasma, at 18.21 ± 6.91 ng/mL in plasma from pregnant women and 78.24 ± 30.85 ng/mL in cord plasma. These values correspond to <1%, 84% and 76% of the 3β-hydroxy-5-ene versions. 20-HC-3,22-diO or 22-HC-3,24- diO is also abundant in placenta. In the derivatisation method employed in this study 24-oxo groups may be derived from natural 24,25-epoxy groups (43), which raises the possibility that this may also be the case here.

Another oxysterol previously found in plasma and partially identified, based on exact mass, retention time and MS^3^ data, is either 3β,25-dihydroxycholest-5-en-26-oic acid (3β,25-diHCA) or 3β,25,x-trihydroxycholest-5-en-y-one (3β,25,x-triHC-y-O) (43). In the present study this compound is found at similar levels in plasma from non-pregnant (6.36 ± 3.05 ng/mL, Table 1, see Supplemental Figure S8A - C) and pregnant women (6.34 ± 2.03 ng/mL, Supplemental Figures S8E - G) but is more abundant in cord plasma (13.02 ± 6.47 ng/mL, see Supplemental Figure S9A - C). As might be expected in light of the data presented above, the 3-oxo-4-ene version i.e. either 25-hydroxy-3-oxocholest-4-en-26-oic acid (25H,3O-CA) or 25,x-dihydroxycholest-4-en-3,y-dione (25,x-diHC-3,y-diO) is absent from non-pregnant women’s plasma (Figure S8A & B), but present at increasing concentration in pregnant women’s (4.29 ± 2.64 ng/mL, Figure S8E - G) and cord plasma (12.12 ± 5.03 ng/mL Figure S9A - C), the latter two concentrations corresponding to 68% and 93% of their 3β-hydroxy-5-ene analogues. Both 3β,25-diHCA or 3β,25,x-triHC-y-O and 25H,3O-CA or 25,x-diHC-3,y-diO are present in placenta (Figure S9E - G).

A further partially identified pair of 3β-hydroxy-5-ene and 3-oxo-4-ene sterol-acids found in pregnant women’s and cord plasma is 3β,x-dihydroxycholest-5-en-26-oic (3β,x-diHCA) and x-hydroxy-3-oxocholest-4-en-26-oic acid (xH,3O-CA, see Supplemental Figure S8E, F & H and Figure S9A, B & D). The extra hydroxy group designated by x is probably on the side-chain. Only the 3β-hydroxy-5-ene sterol-acid is found in non-pregnant females’ plasma (see Supplemental Figure S8A, B & D). The concentration of 3β,x-diHCA increases from non-pregnant women’s plasma (4.68 ± 1.82 ng/mL) to pregnant women’s plasma (7.42 ± 2.95 ng/mL) to cord plasma (34.00 ± 16.87 ng/mL). The concentrations of xH,3O-CA in pregnant women’s and cord plasma are 4.99 ± 2.25 ng/mL and 11.8 ± 4.60 ng/mL, these values correspond to 67% and 35%, respectively, of the equivalent 3β-hydroxy-5-ene sterols. Both 3β,x-diHCA and xH,3O-CA are also present in placenta (see Supplemental Figure S9E, F & H). An alternative identification for this pair of 3β-hydroxy-5-ene and 3-oxo-4-ene sterols is 3β,x,y-trihydroxycholest-5-en-z-one (3β,x,y-triHC-zO) and x,y-dihydroxycholest-4-en-3,z-dione (x,y-diHC-3,z-diO). If this were the correct structure is likely that the extra hydroxy groups designated x and y and the oxo group z are in the side-chain (43).

### HSD3B1 converts 24S-HC and 20S-HC to 24S-HCO and 20S-HCO, respectively

HSD3B1 converts pregnenolone to progesterone (Figure 1) and is highly expressed in placenta (9, 19). HSD3B7, the enzyme that converts 7α-hydroxycholesterols to their equivalent 3-oxo-4-ene equivalents does not use C_27_ sterols lacking the 7α-hydroxy group as a substrate (8), and is not expressed in placenta (9). Thus, HSD3B1 represents a plausible enzyme which is present in placenta that could have 3β-hydroxy-Δ^5^-C_27_-steroid oxidoreductase Δ^5^-isomerase activity and convert 3β-hydroxy-5-enes to their 3-oxo-4-ene equivalents. To test this hypothesis the plasmid encoding human *HSD3B1* (pHSD3B1) was transfected into HEK 293 cells using JetOPTIMUS transfection reagent and the expression of protein confirmed by immunoblot analysis (Supplemental Figure S10). The activity of HSD3B1 expressed in transfected HEK 293 cells towards 1 μM [^2^H_7_]24R/S-HC and [^2^H_7_]20S-HC was investigated in incubation buffer (47). After 1 hr of incubation oxysterols were extracted from cell pellets (approx. 3 × 10^6^ cells) and subjected to EADSA and LC-MS(MS^3^). [^2^H_7_]22S-HCO and [^2^H_7_]7α-HC were included in the extraction solvent to act as internal standards to quantitatively monitor newly formed hydroxycholestenones and any spurious formation of 3-oxo-4-enes during sample preparation, respectively.

As is evident from Figures 9A & E un-transfected cells do not convert [^2^H_7_]24R/S-HC or [^2^H_7_]20S-HC to their 3-ones. However, HEK 293 cells when transfected with HSD3B1 and incubated for 1 hr with 1 μM [^2^H_7_]20S-HC, about 75% of substrate was converted to [^2^H_7_]20S-HCO (Figure 9C & D). Similar results were obtained in incubations with [^2^H_7_]24R/S-HC but conversion to 24S-HCO was only about 35% (Figure 9G & H).

**Figure 9.**
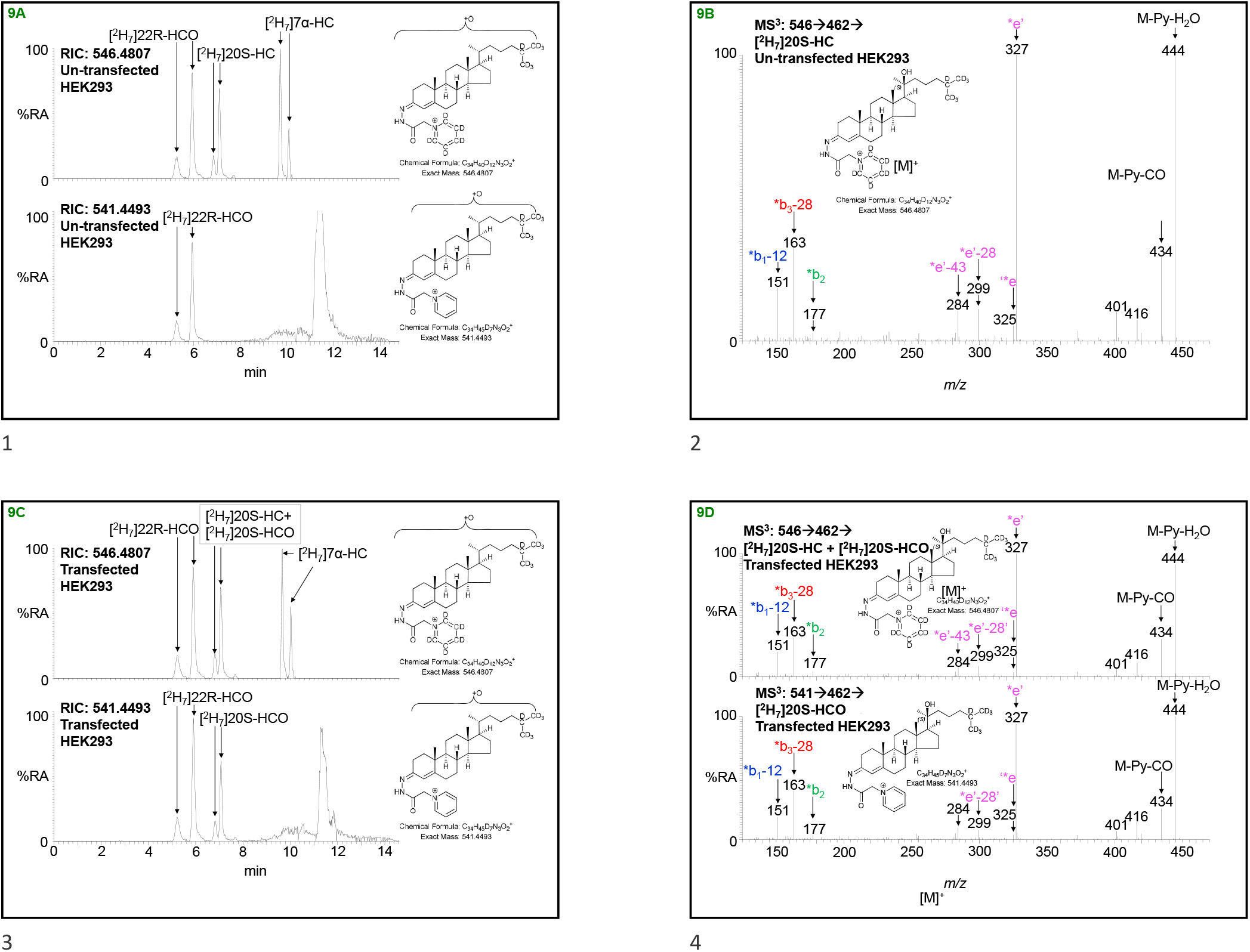
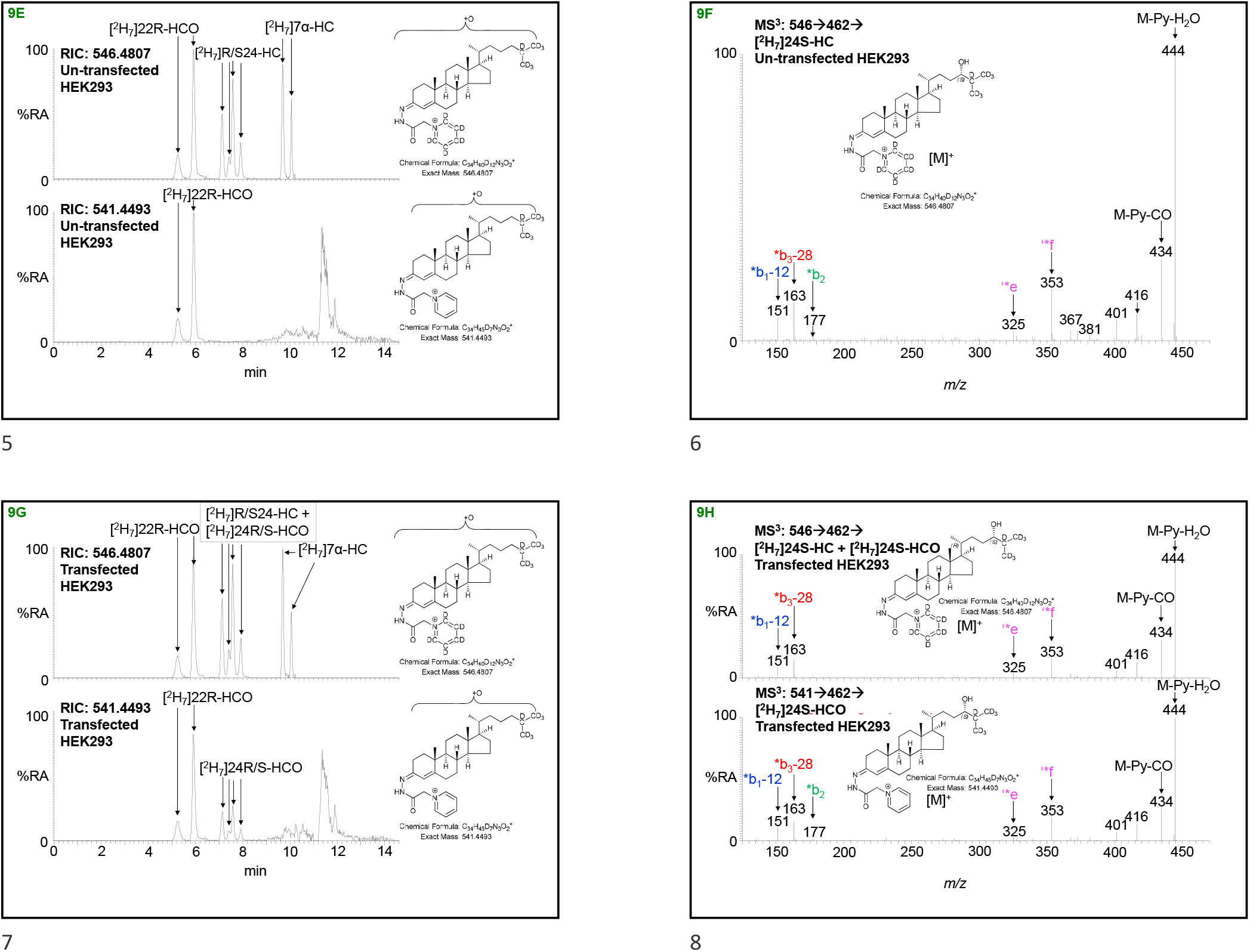
[^2^H_7_]20S-HC and [^2^H_7_]24R/S-HC are converted to [^2^H_7_]20S-HCO and [^2^H_7_]24R/S-HCO by HEK 293 cells transfected with HSD3B1. [^2^H_7_]20S-HC was incubated with un-transfected HEK 293 cells (A - B) or cells transfected with HSD3B1 (C - D). (A & C) RICs of *m/z* 546.4807 corresponding to GP-derivatised [^2^H_7_]20S-HC (upper panel) and *m/z* 541.4493 corresponding [^2^H_7_]20S-HCO (lower panel). [^2^H_7_]22R-HCO and [^2^H_7_]7α-HC were added in the extraction solvent to quantitatively monitor [^2^H_7_]20S-HC formation and any spurious oxidation at C-3. (B & D) MS^3^ ([M]^+^→[M-Py]^+^→) spectra of [^2^H_7_]20S-HC plus [^2^H_7_]20S-HCO, if present (upper panel) and [^2^H_7_]20S-HCO, if present (lower panel). (E & G) RICs of *m/z* 546.4807 corresponding to GP-derivatised [^2^H_7_]24R/S-HC plus [^2^H_7_]24R/S-HCO if present (upper panel) and *m/z* 541.4493 corresponding [^2^H_7_]24R/S-HCO (lower panel). (F & H) MS^3^ ([M]^+^→[M-Py]^+^→) spectra of [^2^H_7_]24S-HC plus [^2^H_7_]24S-HCO if present (upper panel) and [^2^H_7_]24S-HCO, if present (lower panel).

## Discussion

In the current study we have made the surprising observation that oxysterols, including sterol-acids, with a 3-oxo-4-ene function but lacking a 7α-hydroxy group are present in plasma from pregnant women and the umbilical cord. The origin of these unexpected metabolites is likely to be the placenta based on their relative abundance in this tissue. In the analysis of adult plasma, oxysterols with a 3-oxo-4-ene function are nearly always found to be 7α-hydroxylated (38, 41-43, 48). HSD3B7 is the C_27_ oxidoreductase Δ^5^-isomerase which converts the 3β-hydroxy-5-ene group in oxysterols possessing a 7α-hydroxy group to the 3-oxo-4-ene group (Figure 1) (8). HSD3B7 only uses C_27_ oxysterols with a 7α- hydroxy group as its substrates and does not convert pregnenolone to progesterone or oxidise other C_21_ steroids at C-3 (8). Instead, this activity is the preserve of HSD3B1 and 3B2 (19-21, 47). HSD3B2 is primarily expressed in the adrenal gland ovary and testis, while HSD3B1 is primarily localised to placenta and non-steroidogenic tissue (9, 19). In the placenta, HSD3B1 is reported to have both microsomal and mitochondrial locations, the latter being optimal for progesterone synthesis from pregnenolone, as CYP11A1 producing pregnenolone is also mitochondrial (49, 50). As HSD3B1 is a 3β- hydroxysteroid dehydrogenase Δ^5^ isomerase and is abundant in placenta, it represents a good candidate enzyme to catalyse the unexpected conversion of C_27_ 3β-hydroxy-5-ene oxysterols lacking a 7α-hydroxy group to their 3-oxo-4-ene analogues. We performed a proof of principle study where we incubated isotope labelled versions of oxysterols (i.e. [^2^H_7_]24R/S-HC and [^2^H_7_]20S-HC), whose natural S-isomers are present in placenta, with HEK 293 cells transfected with *HSD3B1* and expressing HSD3B1. We selected these oxysterols as the natural-isotopic version of the former is not known to be synthesised in placenta, and is likely derived from maternal blood, while the latter has previously been identified in placenta (44, 45). We found that both oxysterols were converted to their 3-oxo-4- ene forms. This preliminary study confirms that HSD3B1 has activity towards C_27_ oxysterols and is likely to be the enzyme converting oxysterols without a 7α-hydroxy group from their 3β-hydroxy-5-ene to 3-oxo-4-ene form.

When unexpected oxysterols are identified, it is wise to be wary of *ex vivo* autoxidation leading to artefact formation. We were able to confirm that this was not the case here by the inclusion of isotope-labelled standards in the extraction solvent and by noting an absence of isotope-labelled oxysterols with a changed structure. Another potential stage for autoxidation is during sample storage. However, the most labile positions in cholesterol are at C-4 and C-7 allyl to the Δ^5^ double bond (51-54), and it is difficult to conceive a free radical mechanism for the transition of a 3β-hydrox-5-ene group to a 3-oxo-4-ene and a more plausible route is therefore through enzymatic oxidation. Bacterial cholesterol oxidase can convert the 3β-hydroxy-5-ene group to the 3-oxo-4-ene (55), and in our analytical method we perform such a reaction (Supplemental Figure S1), however, we detect the unexpected 3-oxo-4-ene oxysterols in the absence of added cholesterol oxidase. Bacteria in the uterus could provide an alternative source of cholesterol oxidase activity and interestingly mycobacteria can express a 3β-HSD which will convert 25-HC to 25-HCO (56). However, our finding that cells expressing human HSD3B1 will convert the 3β-hydroxy-5-ene function to the 3-oxo-4-ene, at least in [^2^H_7_]24R/S-HC and [^2^H_7_]20S-HC strongly favours the conclusion that the unexpected oxysterols with a 3-oxo-4-ene group are formed endogenously in placenta.

What might be the biological benefit of placental HSD3B1 having activity towards C_27_ sterols? As cord blood is the blood remaining in the umbilical cord and placenta following birth its composition can give clues to the biology of oxysterols with the 3-oxo-4-ene function (Figure 10). The oxysterol composition of cord plasma indicates two unusual pathways of metabolism. In the first we see a pathway of 26-HCO, 3O-CA and 3O-Δ^4^-BA providing a route towards bile acids as shown on the pale-yellow background in Figure 3. 26-HCO may be formed via HSD3B1 oxidation of 26-HC and then metabolised to 3O-CA by CYP27A1. CYP27A1 has been shown to be active towards sterols with a 3-oxo-4-ene structure (57). Peroxisomal side-chain shortening will then lead to 3O-Δ^4^-BA. 26-HC is an LXR ligand (58) and also a selective estrogen receptor modulator (59), so oxidation at C-3 could provide a route to deactivate the ligand. Two alternative routes of deactivation of 26-HC are (i) CYP27A1 mediated oxidation to 3β-HCA, however, 3β-HCA is itself an LXR ligand (60, 61), and (ii) 7α-hydroxylation by CYP7B1 to 7α,26-diHC, but 7α,26-diHC is also biologically active as a chemoattractant of GPR183 expressing immune cells (13). Interestingly, CYP27A1 is a mitochondrial enzyme (4), and HSD3B1 can also be located in this organelle (49), thus HSD3B1 oxidation of 3β-HCA to 3O-CA could act as a route for the former acid’s deactivation. Conversion of the 3β-hydroxy-5-ene-function in 7α,26-diHC, by HSD3B7 to the 3-oxo-4-ene in 7α,26-dihydroxycholest-4-en-3-one (7α,26-diHCO) is an established route of its deactivation (13). In each of the plasma samples 7α,26-diHC is only a minor oxysterol, 7α,26-diHCO being 3 – 7 times more abundant (Table 1, Figure 10).

**Figure 10.**
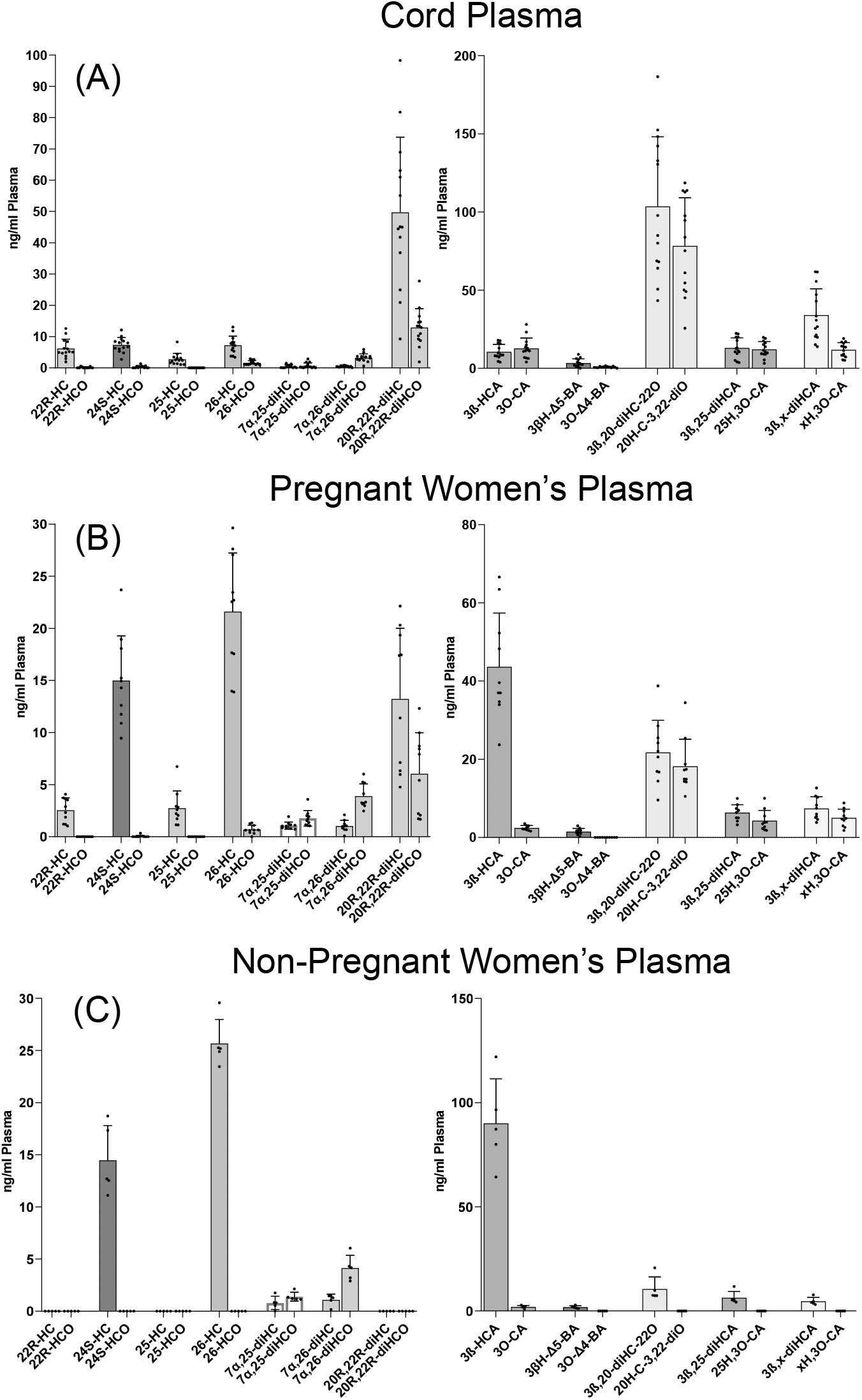
Concentrations of oxysterols in plasma from (A) the umbilical cord, (B) pregnant women and (C) non-pregnant women (controls).

The second unusual metabolic pathway revealed by the analysis of cord plasma also involves oxysterols with a 3-oxo-4-ene structure and encompasses 20R-HC, 20R,22R-diHC, 20R-HCO and 20R,22R-diHCO, presumably leading to progesterone by side-chain cleavage of 20R,22R-diHCO (Figure 3, orange background). This pathway may again be a route to deactivate LXR ligands, in this case 22R-HC and 20R,22R-diHC (24), or a route to progesterone avoiding pregnenolone, although using the same enzymes CYP11A1 and HSD3B1 as in the conventional pathway but in a different order. The introduction to this pathway may be through HSD3B1 oxidation of either 22R-HC or 20R,22R-diHC; CYP11A1 being a 22- and 20-hydroxylase and also the side-chain shortening enzyme (22). It is perhaps significant that like CYP27A1, CYP11A1 is a mitochondrial enzyme, while HSD3B1 also has a mitochondrial location. Certainly, by studying Figure 10 it is evident that 3-oxo-4-ene metabolites are most prevalent in compounds with 20- or 22-hydroxylation (introduced by CYP11A1) or 26-hydroxylation or carboxylation (introduced by CYP27A1). Extrapolating this evidence to 20S-HC and 20S-HCO suggests that the enzyme generating 20S-HC from cholesterol is mitochondrial, perhaps CYP11A1.

20S-HC like many other side-chain oxysterols inhibits the processing of SREBP-2 (27, 28) and is an LXR ligand (24, 26). It is also a reported agonist towards another nuclear receptor, retinoic acid receptor-related orphan receptor γ (RORγ) (62), an activator of the Hh signalling pathway by binding to SMO (29), and a ligand to the σ2 receptor, encoded by *TMEM97* (30). Interestingly, the expression pattern of *TMEM97* is restricted to the central nervous system and proliferating tumours. 20S-HC has previously been reported in human placenta and rodent brain (44-46). The metabolism of 20S-HC has not been studied extensively (1), however, it can bind to the active site of CYP11A1 (63), and has been shown to be converted to pregnenolone (64), although this reaction is not considered as a major route to pregnenolone (1). Besides inhibiting the processing of SREBP-2 (28), 20S-HC will also repress the activity of hydroxymethylglutaryl (HMG) - CoA reductase (65), as will other side-chain oxysterols (66). Although produced primarily in brain (67, 68), 24S-HC is abundant in the circulation (43) (see Table 1 and Figure 10) and has a route from mother to fetus via the placenta. Like 20S-HC, 24S-HC is a ligand to LXRs and inhibits the processing SREBP-2 (25-27), but unlike 20S-HC is an inverse agonist towards RORγ (69). Of these myriads of activities, few have been ascribed to the 3-oxo-4-ene sterols, suggesting once more that HSD3B1 can provide a role in deactivating oxysterols.

It should be pointed out, that very few studies have been performed on oxysterols with 3-oxo-4-ene function but devoid of a 7α-hydroxy group (1, 14, 61, 65, 70). This is probably because they are seldom identified *in vivo*. Interestingly the trio of metabolites 26-HCO, 3O-CA and 3O-Δ^4^-BA provide an exception in that like 25-HC and 26-HC they will each suppress the activity of HMG-CoA reductase in human fibroblasts (70). These cells will convert 26-HCO to 3O-CA (70), in agreement with other studies showing 3-oxo-4-ene sterols are substrates for CYP27A1 (57).

In summary, the placenta facilitates exchange of metabolites between the fetus and mother. It is also an endocrine organ producing hormones that regulate maternal and fetal physiology. The umbilical cord connects the placenta to the fetus and its blood content can be sampled after birth as cord blood representing the fetal blood content of the placenta. By analysing plasma from pregnant women, cord blood and placental tissue along with plasma from non-pregnant women we provide evidence for the conversion of oxysterols with a 3β-hydroxy-5-ene structure to 3-oxo-4-ene analogues in placenta. Based on a proof-of-principle study this activity is likely to be catalysed by HSD3B1. We speculate that these unexpected reactions provide a mechanism to regulate biologically active oxysterols and deserve further study.

## Supplementary material

See Supplemental file.

## Acknowledgements

This work was supported by funding from the Biological Sciences Research Council (BBSRC, grant numbers BB/I001735/1, BB/N015932/1 and BB/S019588/1 to WJG, BB/L001942/1 to YW), and the European Union, through European Structural Funds (ESF), as part of the Welsh Government funded Academic Expertise for Business project (to WJG and YW). ALD was supported via a KESS2 award with Markes International from the Welsh Government and European Social Fund. Work at the Oxford laboratory was supported by the LEAN network grant by the Leducq Foundation (UO). JED is supported by NIHR Oxford Biomedical Research Centre, Oxford UK. We are grateful to Dr Peter Douglas of Swansea University for provision of facilities and guidance for synthetic chemistry work. Dr Peter Grosshans and Steve Smith of Markes International are thanked for helpful discussions. Members of the European Network for Oxysterol Research (ENOR, https://www.oxysterols.net/) are thanked for informative discussions.

