## Supplemental for "HSD3B1 is an Oxysterol 3β-Hydroxysteroid Dehydrogenase in Human Placenta"

**Supplemental Methods**

*Materials*

Isotope-labelled standards [25,26,26,26,27,27,27-$^2H_7$]24R/S-hydroxycholesterol ([$^2H_7$]24R/S-HC), [25,26,26,26,27,27,27-$^2H_7$]22R-hydroxycholesterol ([$^2H_7$]22R-HC), [25,26,26,26,27,27,27-$^2H_7$]7α-hydroxycholesterol ([$^2H_7$]7α-HC), were from Avanti Polar Lipids, Alabaster, AL. [25,26,26,26,27,27,27-$^2H_7$]20S-Hydroxycholesterol ([$^2H_7$]20S-HC) was purchased from Toronto Research Chemicals (TCI, Toronto, Canada). [$^2H_7$]22R-Hydroxycholest-4-en-3-one ([$^2H_7$]22R-HCO) was prepared from [$^2H_7$]22R-HC by treatment with cholesterol oxidase enzyme (*Streptomyces sp*., Merck, Dorset, UK) (1).

*Extraction of Oxysterols from Placenta*

Exact details of the extraction of oxysterols from placenta can be found in (2). In brief, oxysterols were extracted from placenta using a modified protocol previously used to extract oxysterols from liver (3). About 400 mg of tissue from the maternal side of placenta was washed with PBS and transferred to a gentleMACS™ C tube (Miltenyi Biotec, Woking, UK). 4.2 mL of absolute ethanol containing 50 ng of [$^2H_7$]24R/S-HC and 50 ng of [$^2H_7$]22R-HCO was added, and the tissue was homogenised for 2 min. The homogenate was transferred to a 15 mL Corning tube and sonicated for 15 min during which time 1.8 mL of $H_2O$ was added dropwise to give 6 mL of homogenate in 70% ethanol. The homogenate was then centrifuged for 1 hr at 4,000 x *g.* The supernatant was transferred to a fresh Corning tube and the remaining pellet re-suspended in a further 4.2 mL of ethanol containing 50 ng of [$^2H_7$]24R/S-HC and 50 ng of [$^2H_7$]22R-HCO. The suspension was vortex mixed and transferred back into the original gentleMACS™ C tube where it was then homogenised for a further 2 min. The homogenate was removed and sonicated for 15 min, then 1.8 mL of $H_2O$ added. After centrifugation, the supernatants from the two extractions were combined. This solution was vortexed and sonicated for a further 10 min then centrifuged for 1 hr at 4,000 x *g*. 10 % (1.2 mL) of the total supernatant was added to 300 µL of 70% ethanol under sonication. The 1.5 mL of sample was subjected to solid phase extraction (SPE) by a procedure modified from references (3, 4).

Sample was loaded onto a Certified Sep-Pak $C_{18}$, 200 mg (Waters Inc. Elstree, UK) SPE column previously conditioned with ethanol (4 mL) and 70% ethanol (6 mL). The sample flow-through (1.5 mL) was combined with a column wash of 70% ethanol (5.5 mL) resulting in SPE1-Fr1 (7 mL) which contained oxysterols and sterol acids. A second fraction was obtained by further washing with 70% ethanol (4 mL) and collected as SPE1-Fr2. Cholesterol and other sterols of similar lipophilicity were eluted from the SPE column with absolute ethanol (2 mL) to give SPE1-Fr3. A fourth fraction was eluted with a further 2 mL of absolute ethanol (SPE1-Fr4). Each of the four fractions was divided equally into A and B sub-fractions and dried overnight under vacuum (ScanLaf ScanSpeed vacuum concentrator, Lynge, Denmark). Only fraction SPE1 was processed further.

The lyophilised samples were dissolved in propan-2-ol (100 µL) and mixed by vortex. To fraction-A, 50 mM $K_2HPO_4$ buffer, pH 7 (1 mL) containing cholesterol oxidase solution (3.0 µL, 2 mg/mL in $H_2O$, 44 units/mg of protein) was added. The sample was vortex mixed and incubated at 37°C for 1 hr. The reaction was then quenched by the addition of methanol (2 mL). Fraction-B was treated in parallel in an identical fashion to fraction-A but in the absence of cholesterol oxidase. Glacial acetic acid (150 µL) was added to fractions-A and -B and mixed by vortex. [$^2H_5$]Girard P (GP) reagent (1) (190 mg, bromide salt) was added to fraction-A and [$^2H_0$]GP reagent (150 mg, chloride salt, TCI Europe, Oxford UK) was added to fraction-B. The samples were mixed by vortex. The reaction was left to proceed overnight at room temperature protected from light.

An OASIS HLB 60 mg SPE cartridge (Waters Inc) was washed with methanol (6 mL), 5% methanol (6 mL) and conditioned with 70% methanol (4 mL). Sample from above (3.25 mL, 69% organic) was loaded onto the column and the flow-through collected. The sample tube was washed with 70% methanol (1 mL) which was loaded onto the SPE column and the eluent combined with the earlier flow-through. The column was re-conditioned with 35% methanol (1 mL) and the eluent combined with the earlier collection. The total eluent (~5 mL) was diluted with 4 mL of $H_2O$ to give ~9 mL of 35% methanol. The 9 mL solution was loaded onto the column and the flow-through collected. The column was re-conditioned with 17.5% methanol (1 mL) and the eluent combined with earlier flow-through. To the combined collection, $H_2O$ (9 mL) was added to give 19 mL of 17.5% methanol. This solution was loaded onto the column and the flow-through collected. The sorbent was re-conditioned with 8.75% methanol (1 mL) and the flow-through combined with the earlier collection. The total combined eluent of 20 mL was diluted with 19 mL of $H_2O$ to give 39 mL 8.75% methanol. The solution was loaded onto the column and the flow-through discarded. A 5% methanol solution (6 mL) was used to wash the column before the analytes were eluted. The samples were eluted into four separate 1.5 mL microcentrifuge tubes using 3 x 1 mL methanol followed by 1 mL ethanol to give SPE2-FR1, -Fr2, -Fr3, -Fr4. Oxysterols originating from SPE1-Fr1 elute across SPE2-Fr1 and SPE2-Fr2.

Immediately prior to LC-MS analysis of oxysterols, equal volumes of SPE2-Fr1A and SPE2-Fr2A were combined with equal aliquots of SPE2-Fr1B and SPE2-Fr2B and diluted to 60% methanol.

*Extraction of Oxysterols from Plasma*

The exact extraction protocol for oxysterols from plasma can be found in (2) and was essentially that described previously (1), except single phase extraction was into acetonitrile rather than ethanol. 100 µL of plasma was added dropwise to a solution of acetonitrile (1.05 mL) containing 20 ng of [$^2H_7$]24R/S-HC and 20 ng of [$^2H_7$]22R-HCO in an ultrasonic bath. After 5 min of sonication, 350 µL of $H_2O$ was added. The sample (1.5 mL), now in 70% acetonitrile, was sonicated for a further 5 min and centrifuged at 17,000 x *g* at 4°C for 30 min. The sample was subjected to SPE as described above and prepared for LC-MS analysis, except SPE1 was conditioned with 70% acetonitrile rather than 70% ethanol.

*Extraction of Oxysterols from Cells following Transfection studies*

After incubation, cells were gently removed using a cell scraper and cells and incubation buffer transferred to a 2 mL microcentrifuge tube. The tube was centrifuged at 11,000 x g for 5 min to pellet the cells. The supernatant was transferred into a fresh 2 mL tube. As the cells had been incubated with [$^2H_7$]20S-HC or [$^2H_7$]24R/S-HC, these compounds could not be used as internal standards in the LC-MS analysis. Instead, 2 ng of [$^2H_7$]22R-HCO was used for quantification. A mix of 1.05 mL of ethanol containing the [$^2H_7$]22R-HCO was added to the cell pellet (approximately 3.2 x $10^6$ cells in ~50 µL of buffer) whilst sonicating. After 5 min 350 µL of $H_2O$ was added. The sample was sonicated for a further 10 min. The sample was now in ~70% ethanol and suitable for application to SPE-1 and processing as for the placental extract.

*LC-MS(MS$^n$)*

Exact details of LC-MS analysis can be found in (2). In brief, LC-MS was performed on a Dionex Ultimate 3000 UHPLC system (Thermo Fisher Scientific, Hemel Hempstead, UK) interfaced via an electrospray ionisation (ESI) probe to an Orbitrap Elite MS (Thermo Fisher Scientific). Chromatographic separation was carried out on a Hypersil Gold reversed phase $C_{18}$ column (1.9 µm particle size, 50 x 2.1 mm, Thermo Fisher Scientific, UK), with mobile phase A: 33.3% methanol, 17.7% acetonitrile containing 0.1% formic acid; and mobile phase B: 63.3% methanol, 31.6% acetonitrile containing 0.1% formic acid. The gradient started at 20% B for 1 min, rising to 80% B in 7 min and staying at 80% B for 5 min, before returning to 20% B in 6 s and continuing at 20% B for 3 min 54 s. MS analysis on the Orbitrap Elite was performed in the positive-ion mode with five scan events, one high resolution (120,000 full width at half maximum height at *m/z* 400) scan over the *m/z* range 400 – 610 in the Orbitrap and four $MS^3$ scans performed in parallel in the linear ion trap (LIT). Mass accuracy in the Orbitrap was typically < 5 ppm. Injection volumes were 35 µL for plasma extracts and 90 µL for placental homogenate extracts.

**Supplemental Figures**

**Figure S1.** Schematic of the EADSA method. In fraction-A sterols with a natural 3-oxo-4-ene group (drawn in claret) and ones generated by bacterial cholesterol oxidase treatment (in dark blue) are derivatised with $[^2H_5]$GP (in orange). In fraction-B, sterols with a natural 3-oxo-4-ene function (drawn in claret) are derivatised with $[^2H_0]$GP (in purple). Fractions-A and -B are combined and analysed by LC-MS.

**Figure S2.** $MS^3$ total ion chromatograms (TICs) demonstrating the presence of 3O-CA and 26-HCO in plasmas and placenta. TIC for the transitions $[M]^+ \rightarrow [M-Py]^+ \rightarrow$ applicable to of 3β-HCA plus 3O-CA (upper panels) and 3O-CA (lower panels) in (A) non-pregnant woman's (control) plasma, (B) pregnant woman's plasma, (C) cord plasma and (D) placenta; and 26-HC plus 26-HCO, when present (upper panels) and 26-HCO if present (lower panels) in (E) non-pregnant woman's (control) plasma, (F) pregnant woman's plasma, (G) cord plasma, and (H) placenta. Chromatograms in upper and lower panels are plotted on the same y-axis and magnified as indicated. There is some shift in retention time between samples which were analysed at different times on different LC columns but of the same type.

**Figure S3.** 3-Oxo-4-ene oxysterols are not artefacts of sample preparation or analysis. $[^2H_7]$24R/S-HC, $[^2H_7]$7α-HC and $[^2H_7]$22R-HCO were added as internal standards during sample extraction. RIC of 546.4807 ± 5 ppm and TICs for the $MS^3$ transitions 546.5→462.4→ show $[^2H_7]$24R/S-HC, $[^2H_7]$7α-HC and $[^2H_7]$22R-HCO labelled with $[^2H_5]$GP (upper panels). Only $[^2H_7]$22R-HCO becomes labelled with $[^2H_0]$GP and is observed in the RIC of 541.4493 ± 5 ppm and TIC 541.5→462.4→ (lower panels). (A - B) Non-pregnant woman's (control) plasma, (C - D) pregnant woman's plasma, (E - F) cord plasma, and (G- H) placenta. Chromatograms in upper and lower panels are plotted on the same y-axis and magnified as indicated. There is some shift in retention time between samples which were analysed at different times on different LC columns but of the same type.

**Figure S4.** 3O-$\Delta^4$-BA is present in pregnant women's plasma, cord plasma and placenta. TICs for the $MS^3$ transitions 511.4→427.3→ applicable to 3βH-$\Delta^5$-BA plus 3O-$\Delta^4$-BA, when present (upper panel) and 506.3→427.3→ applicable to 3O-$\Delta^4$-BA, if present (lower panels). (A) Non-pregnant female (control) plasma, (B) pregnant woman's plasma, (C) cord plasma and (D) placenta. Chromatograms in upper and lower panels are plotted on the same y-axis and magnified as indicated. There is some shift in retention time between samples which were analysed at different times on different LC columns but of the same type.

**Figure S5.** 20R,22R-diHCO is present in pregnant woman's and cord plasma and in placenta. $MS^3$ TICs for the transitions 555.4→471.4→ applicable to dihydroxycholesterols plus dihydroxycholestenones (upper panels) and 550.4→471.4→ applicable to dihydroxycholestenones (lower panels). (A) Non-pregnant woman's (control) plasma, (B) pregnant woman's plasma, (C) cord plasma and (D) placenta. Chromatograms in upper and lower panels are plotted on the same y-axis and magnified as indicated. There is some shift in retention time between samples which were analysed at different times on different LC columns but of the same type.

**Figure S6.** 20S-HC plus 20S-HCO and 24S-HC plus 24S-HCO are present in placenta. (A) $MS^3$ TIC 539.4→455.4→ applicable 20S-HC plus 20S-HCO and 24S-HC plus 24S-HCO (upper panel) and 534.4→455.4→ applicable to 20S-HCO and 24S-HCO (lower panel).

**Figure S7.** Dihydroxycholest-5-en-x-ones and hydroxycholest-4-ene-3,x-diones are present in plasma and placenta. $MS^3$ ($[M]^+$→$[M\text{-}Py]^+$→) spectra of 3β,20-diHC-22O plus 20-HC-3,22-diO, if present (upper panels) and 20-HC-3,22-diO, when present (lower panels) from (A) non-pregnant female (control) plasma, (B) pregnant woman's plasma, (C) cord plasma, and (D) placenta. Alternative structures to 3β,20-diHC-22O and 20-HC-3,22-diO are 3β,22-diHC-24O and 22-HC-3,24-diO, respectively. See Supplemental Figure S2 for chromatograms.

**Figure S8.** RICs of *m/z* 569.4110 ± 5 ppm (upper panel) and *m/z* 564.3796 (lower panel) from (A) non-pregnant woman's (control) and (E) pregnant woman's plasma. $MS^3$ TICs for the $MS^3$ ($[M]^+$→$[M\text{-}Py]^+$→) transitions 569.4→485.3→ (upper panels) and 564.4→485.3→ (lower panels) from (B) non-pregnant woman's and (F) pregnant woman's plasma. $MS^3$ ($[M]^+$→$[M\text{-}Py]^+$→) spectra of 3β,25-diHCA plus 25H,3O-CA, when present (upper panels) and of 25H,3O-CA when present (lower panels) from (C) non-pregnant woman's (control) plasma, (G) pregnant woman's plasma. $MS^3$ ($[M]^+$→$[M\text{-}Py]^+$→) spectra of 3β,x-diHCA plus xH,3O-CA, when present (upper panels) and xH,3O-CA, if present (lower panels) from (D) non-pregnant woman's (control) plasma, (H) regnant woman's plasma. Identification are made in the absence of authentic standards and are based on exact mass, retention time and $MS^3$ spectra. Chromatograms in upper and lower panels are plotted on the same y-axis and magnified as indicated. There is some shift in retention time between samples which were analysed at different times on different LC columns but of the same type.

**Figure S9.** RICs of *m/z* 569.4110 ± 5 ppm (upper panel) and *m/z* 564.3796 (lower panel) from (A) cord plasma and (E) placenta. TIC for the $MS^3$ ($[M]^+$→$[M\text{-}Py]^+$→) transitions 569.4→485.3→ (upper panels) and 564.4→485.3→ (lower panels) from (B) cord plasma and (F) placenta. $MS^3$ ($[M]^+$→$[M\text{-}Py]^+$→) spectra of 3β,25-diHCA plus 25H,3O-CA (upper panels) and of 25H,3O-CA (lower panels) from (C) cord plasma, and (G) placenta. $MS^3$ ($[M]^+$→$[M\text{-}Py]^+$→) spectra of 3β,x-diHCA plus xH,3O-CA (upper panels) and xH,3O-CA (lower panels) from (D) cord plasma, and (H) placenta. Identifications are made in the absence of authentic standards and are based on exact mass, retention time and $MS^3$ spectra. Chromatograms in upper and lower panels are plotted on the same y-axis and magnified as indicated. There is some shift in retention time between samples which were analysed at different times on different LC columns but of the same type.

**Figure S10**. Western Blot analysis of HSD3B1. Lane 1 protein ladder; lanes 2 – 4 replicates of HEK293 cells transfected with plasmid encoding human *HSD3B1* (pHSD3B1); lane 5 human placenta tissue; lane 6 transfection with empty vector; lane 7 non-transfected cells. Lanes 9 - 11 replicates from incubation experiments with cells transfected with *HSD3B1* plasmids; lane 12 human placenta tissue; lane 13 transfection with empty vector; lanes 14 & 15 non-transfected cells. The top membrane image is the REVERT total protein stain showing lanes loaded with cell extracts or protein ladder lanes. The bottom membrane image is after the membrane was stained with αHSD3B1 (r) followed by the fluorophore containing secondary antibody stain α rabbit 800 and visualised on a LI-COR® Odyssey® System. The HSD3B1 enzyme appears as a distinct band at 42 kDa in transfected cells and human placenta tissue.

**Figure S11.** [$^2H_7$]20S-HC and [$^2H_7$]24R/S-HC are converted to [$^2H_7$]20S-HCO and [$^2H_7$]24R/S-HCO, respectively, by HEK 293 cells transfected with plasmid encoding human *HSD3B1* (pHSD3B1) but not by un-transfected cells. $MS^3$ ($[M]^+ \rightarrow [M\text{-}Py]^+ \rightarrow$) TICs for the transitions 546.5→462.4→ appropriate for [$^2H_7$]monohydroxycholesterols and [$^2H_7$]monohydroxycholestenones (upper panels), and 541.5→462.4→ appropriate for [$^2H_7$]monohydroxycholestenones. (A) Un-transfected cells incubated with [$^2H_7$]20S-HC. (B) Transfected cells incubated with [$^2H_7$]20S-HC. (C) Un-transfected cells incubated with [$^2H_7$]24R/S-HC. (D) Transfected cells incubated [$^2H_7$]24R/S-HC. [$^2H_7$]22R-HCO and [$^2H_7$]7α-HC were added in extraction solvent to allow approximate quantification and monitor spurious oxidation.


1. Crick, P. J., T. William Bentley, J. Abdel-Khalik, I. Matthews, P. T. Clayton, A. A. Morris, B. W. Bigger, C. Zerbinati, L. Tritapepe, L. Iuliano, Y. Wang, and W. J. Griffiths. 2015. Quantitative charge-tags for sterol and oxysterol analysis. *Clin Chem* **61**: 400-411.
2. Dickson, A. L., E. Yutuc, C. A. Thornton, Y. Wang, and W. J. Griffiths. 2022. Identification of Unusual Oxysterols Biosynthesised in Human Pregnancy by Charge-Tagging and Liquid Chromatography - Mass Spectrometry. *bioRxiv*: 2022.2002.2007.478301.
3. Raselli, T., T. Hearn, A. Wyss, K. Atrott, A. Peter, I. Frey-Wagner, M. R. Spalinger, E. M. Maggio, A. W. Sailer, J. Schmitt, P. Schreiner, A. Moncsek, J. Mertens, M. Scharl, W. J. Griffiths, M. Bueter, A. Geier, G. Rogler, Y. Wang, and B. Misselwitz. 2019. Elevated oxysterol levels in human and mouse livers reflect nonalcoholic steatohepatitis. *J Lipid Res* **60**: 1270-1283.
4. Yutuc, E., A. L. Dickson, M. Pacciarini, L. Griffiths, P. R. S. Baker, L. Connell, A. Öhman, L. Forsgren, M. Trupp, S. Vilarinho, Y. Khalil, P. T. Clayton, S. Sari, B. Dalgic, P. Höflinger, L. Schöls, W. J. Griffiths, and Y. Wang. 2021. Deep mining of oxysterols and cholestenoic acids in human plasma and cerebrospinal fluid: Quantification using isotope dilution mass spectrometry. *Anal Chim Acta* **1154**: 338259.

# S1

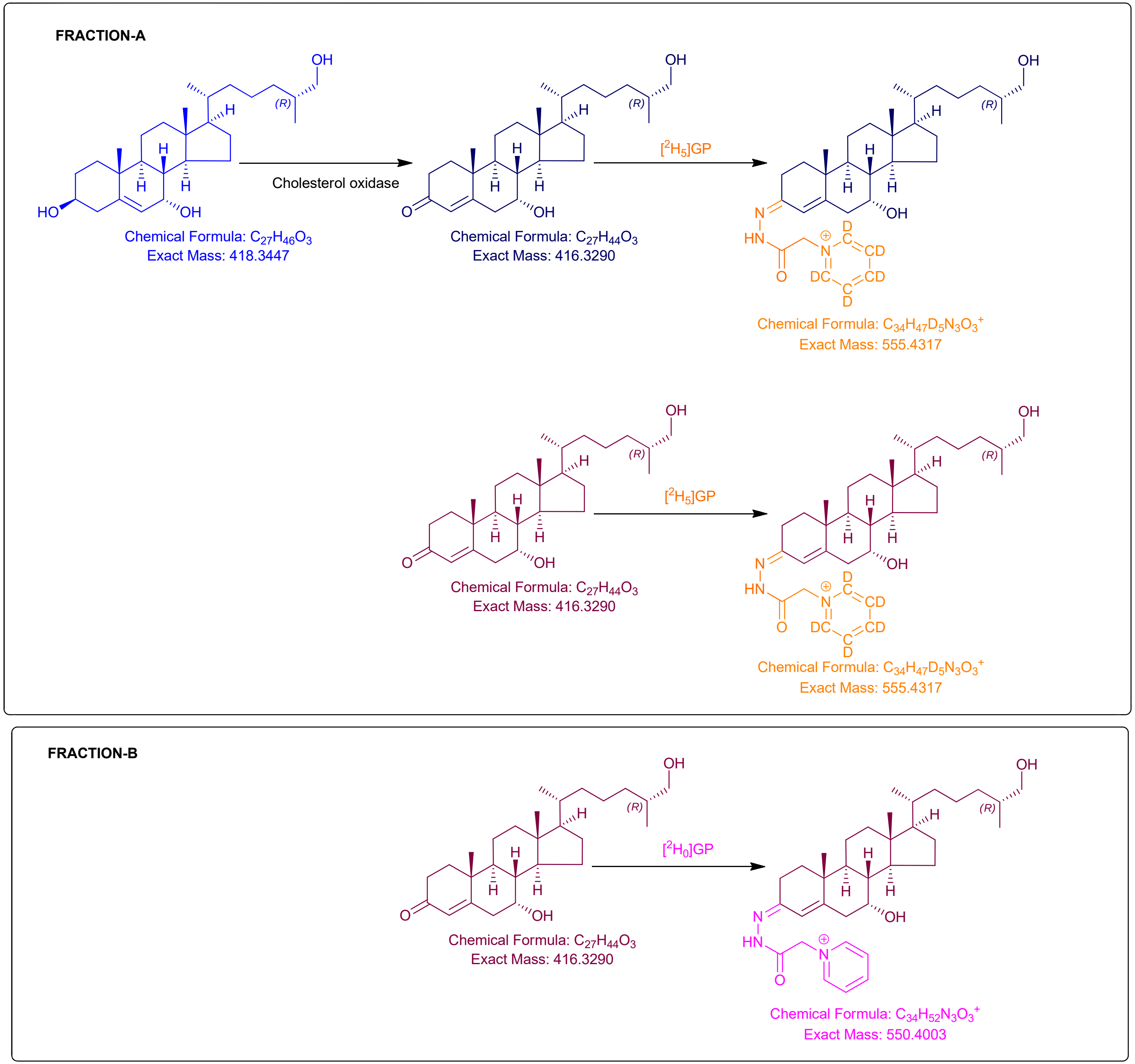
FRACTION-A
Cholesterol oxidase
Chemical Formula: $C_{27}H_{46}O_3$
Exact Mass: 418.3447
Chemical Formula: $C_{27}H_{44}O_3$
Exact Mass: 416.3290
[$^2H_5$]GP
Chemical Formula: $C_{34}H_{47}D_5N_3O_3^+$
Exact Mass: 555.4317
Chemical Formula: $C_{27}H_{44}O_3$
Exact Mass: 416.3290
[$^2H_5$]GP
Chemical Formula: $C_{34}H_{47}D_5N_3O_3^+$
Exact Mass: 555.4317
FRACTION-B
Chemical Formula: $C_{27}H_{44}O_3$
Exact Mass: 416.3290
[$^2H_0$]GP
Chemical Formula: $C_{34}H_{52}N_3O_3^+$
Exact Mass: 550.4003

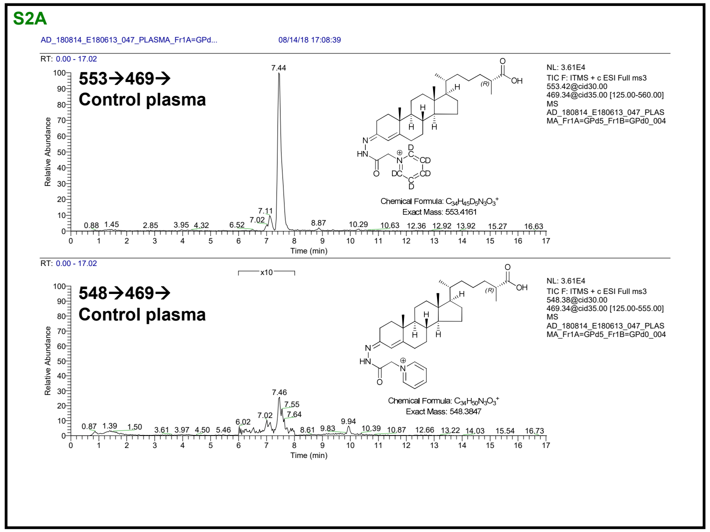
S2A
553→469→
Control plasma
548→469→
Control plasma

1

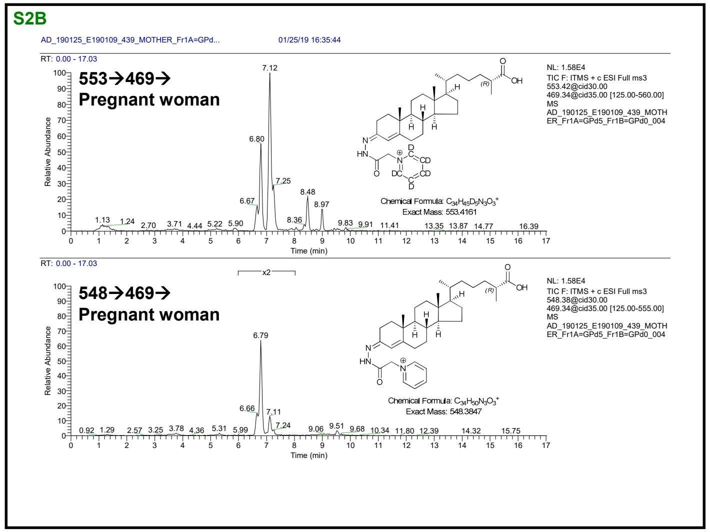
S2B
553→469→
Pregnant woman
548→469→
Pregnant woman

2

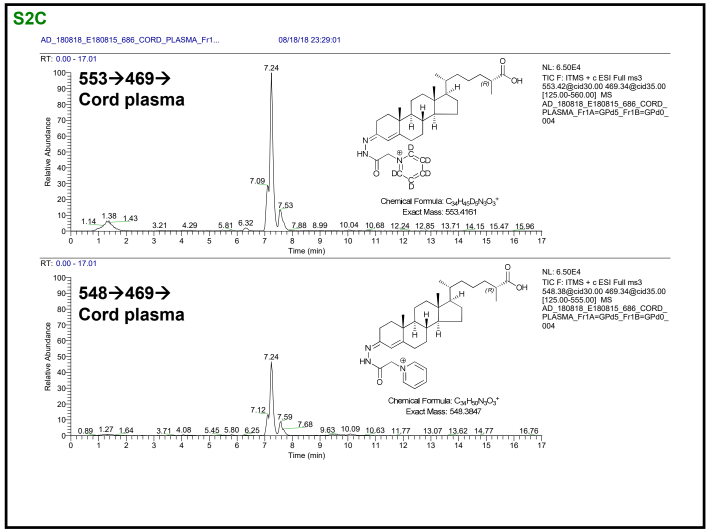
S2C
553→469→
Cord plasma
548→469→
Cord plasma

3

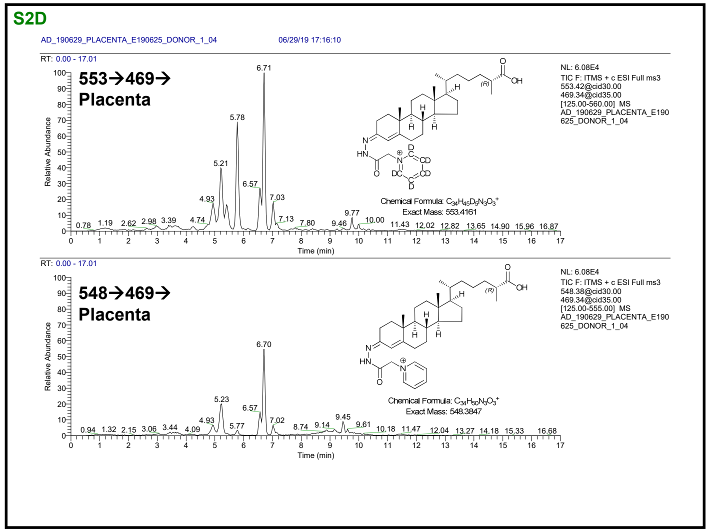
S2D
553→469→
Placenta
548→469→
Placenta

4

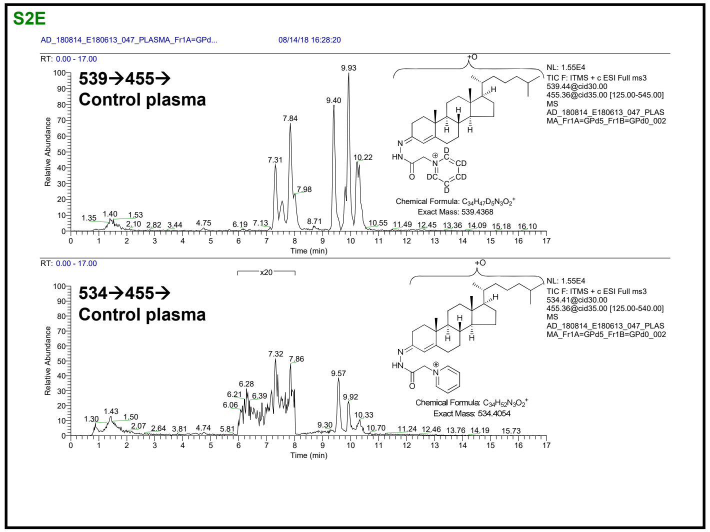
S2E
AD_180814_E180613_047_PLASMA_Fr1A=GPd...
08/14/18 16:28:20
539→455→
Control plasma
534→455→
Control plasma
Chemical Formula: $C_{34}H_{47}D_5N_3O_2^+$
Exact Mass: 539.4368
Chemical Formula: $C_{34}H_{52}N_3O_2^+$
Exact Mass: 534.4054
Relative Abundance
Time (min)

5

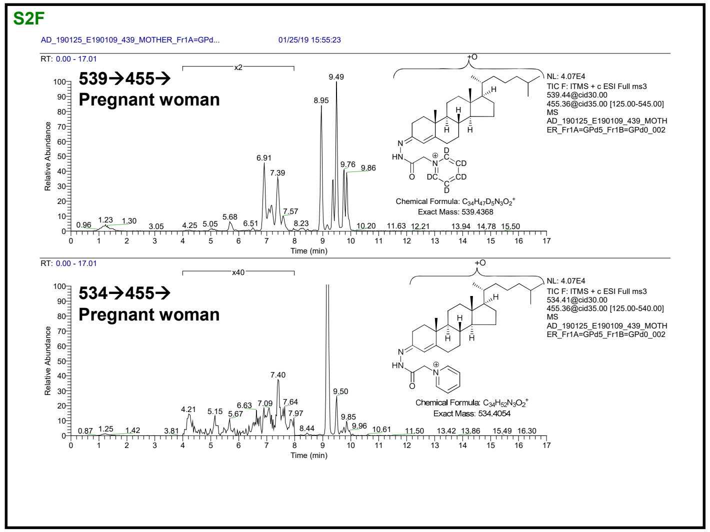
S2F
AD_190125_E190109_439_MOTHER_Fr1A=GPd...
01/25/19 15:55:23
539→455→
Pregnant woman
534→455→
Pregnant woman
Chemical Formula: $C_{34}H_{47}D_5N_3O_2^+$
Exact Mass: 539.4368
Chemical Formula: $C_{34}H_{52}N_3O_2^+$
Exact Mass: 534.4054
Relative Abundance
Time (min)

6

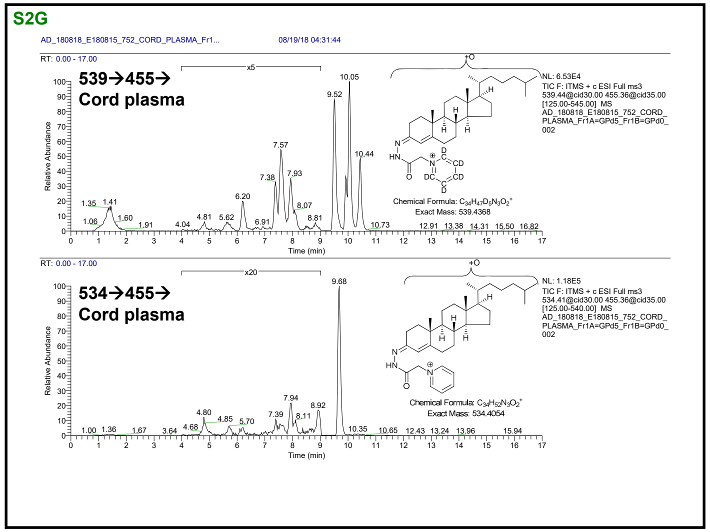
S2G
AD_180818_E180815_752_CORD_PLASMA_Fr1...
08/19/18 04:31:44
539→455→
Cord plasma
534→455→
Cord plasma
Chemical Formula: $C_{34}H_{47}D_5N_3O_2^+$
Exact Mass: 539.4368
Chemical Formula: $C_{34}H_{52}N_3O_2^+$
Exact Mass: 534.4054
Relative Abundance
Time (min)

7

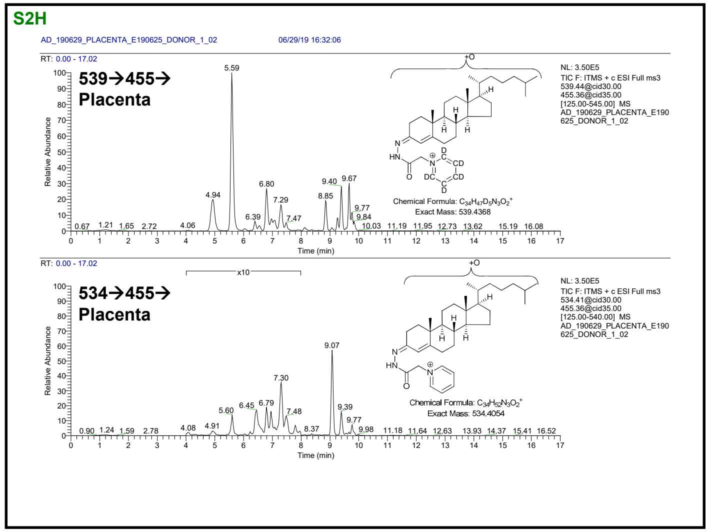
S2H
AD_190629_PLACENTA_E190625_DONOR_1_02
06/29/19 16:32:06
539→455→
Placenta
534→455→
Placenta
Chemical Formula: $C_{34}H_{47}D_5N_3O_2^+$
Exact Mass: 539.4368
Chemical Formula: $C_{34}H_{52}N_3O_2^+$
Exact Mass: 534.4054
Relative Abundance
Time (min)

8

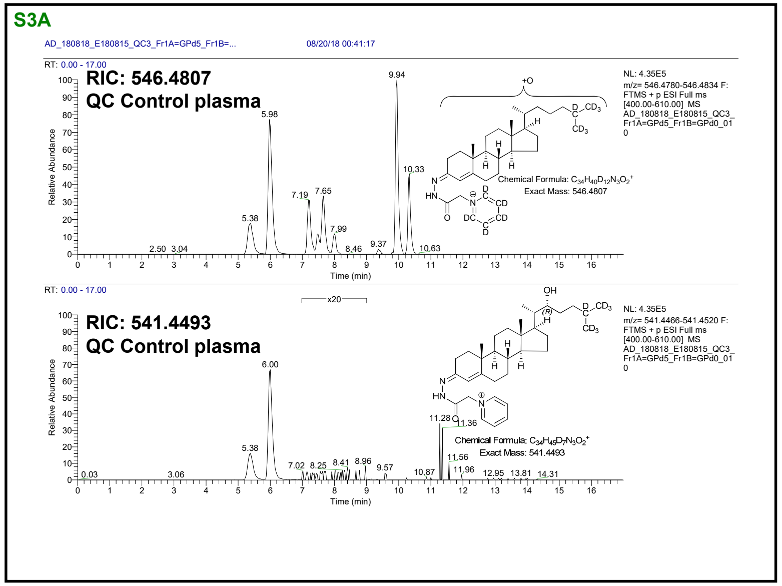
S3A
AD_180818_E180815_QC3_Fr1A=GPd5_Fr1B=...
08/20/18 00:41:17
RIC: 546.4807
QC Control plasma
Chemical Formula: $C_{34}H_{40}D_{12}N_3O_2^+$
Exact Mass: 546.4807
RIC: 541.4493
QC Control plasma
Chemical Formula: $C_{34}H_{45}D_7N_3O_2^+$
Exact Mass: 541.4493
Relative Abundance
Time (min)

1

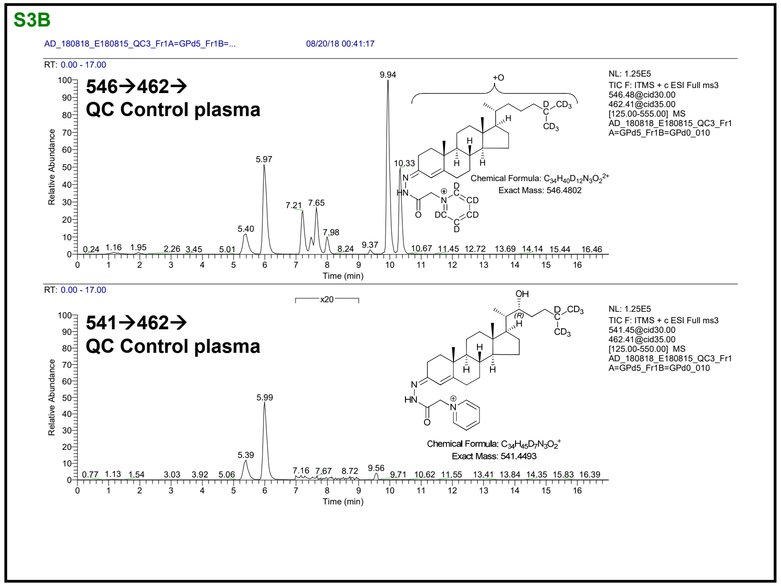
S3B
AD_180818_E180815_QC3_Fr1A=GPd5_Fr1B=...
08/20/18 00:41:17
546→462→
QC Control plasma
Chemical Formula: $C_{34}H_{40}D_{12}N_3O_2^{2+}$
Exact Mass: 546.4802
541→462→
QC Control plasma
Chemical Formula: $C_{34}H_{45}D_7N_3O_2^+$
Exact Mass: 541.4493
Relative Abundance
Time (min)

2

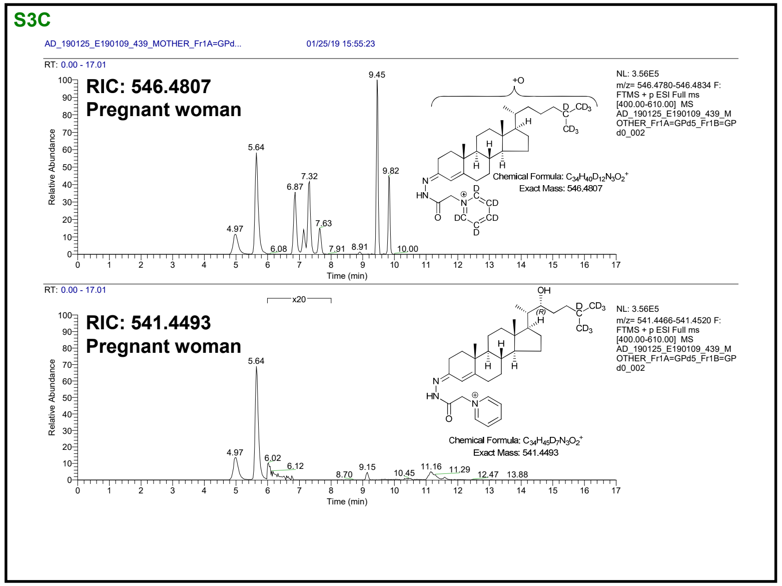
S3C
AD_190125_E190109_439_MOTHER_Fr1A=GPd...
01/25/19 15:55:23
RIC: 546.4807
Pregnant woman
Chemical Formula: $C_{34}H_{40}D_{12}N_3O_2^+$
Exact Mass: 546.4807
RIC: 541.4493
Pregnant woman
Chemical Formula: $C_{34}H_{45}D_7N_3O_2^+$
Exact Mass: 541.4493
Relative Abundance
Time (min)

3

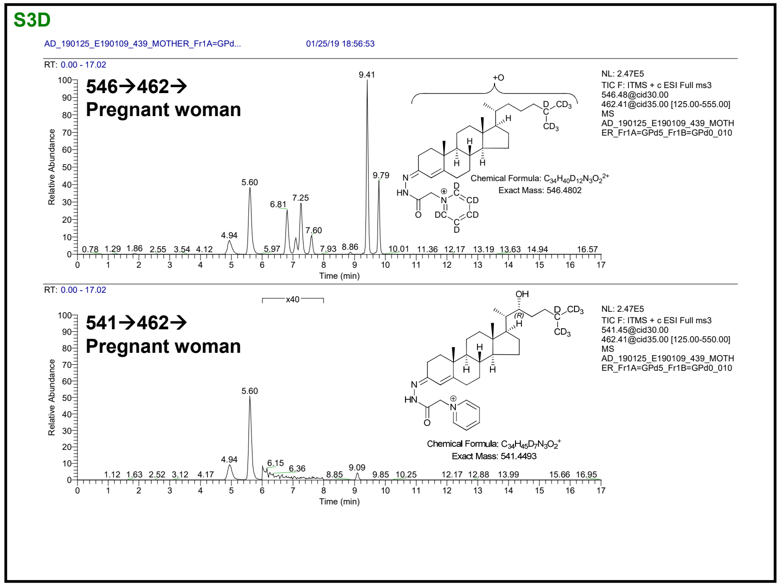
S3D
AD_190125_E190109_439_MOTHER_Fr1A=GPd...
01/25/19 18:56:53
546→462→
Pregnant woman
Chemical Formula: $C_{34}H_{40}D_{12}N_3O_2^{2+}$
Exact Mass: 546.4802
541→462→
Pregnant woman
Chemical Formula: $C_{34}H_{45}D_7N_3O_2^+$
Exact Mass: 541.4493
Relative Abundance
Time (min)

4

S3E

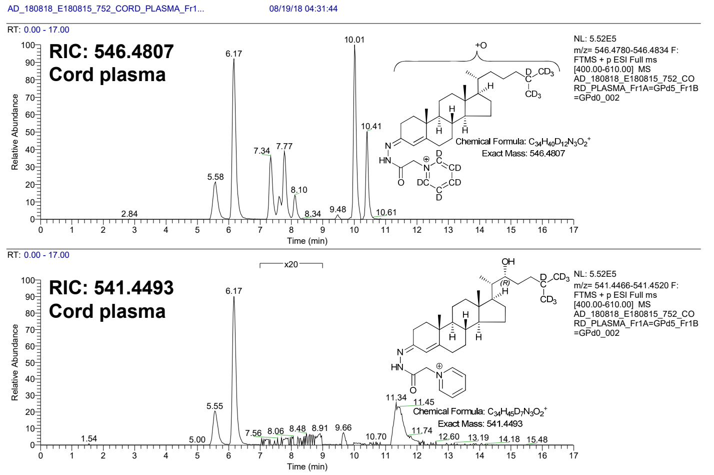


5

S3F

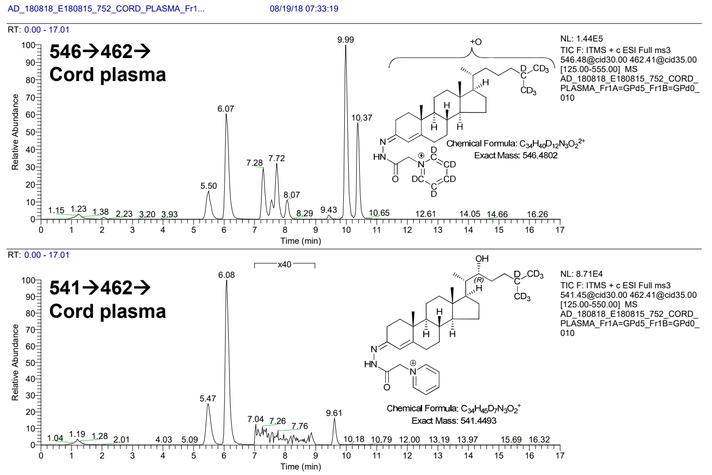


6

S3G

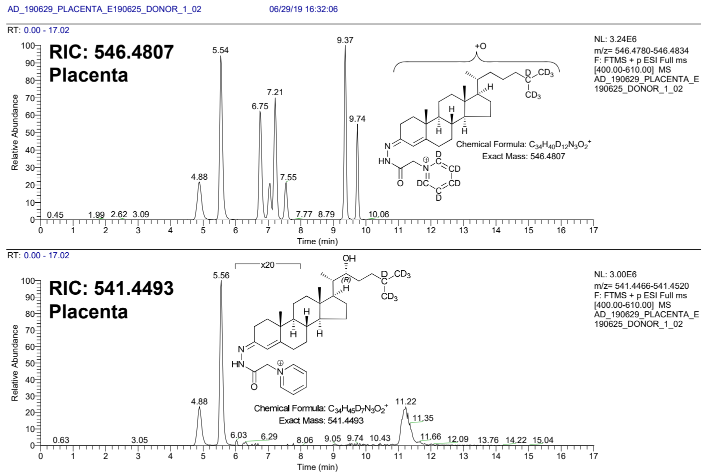


7

S3H

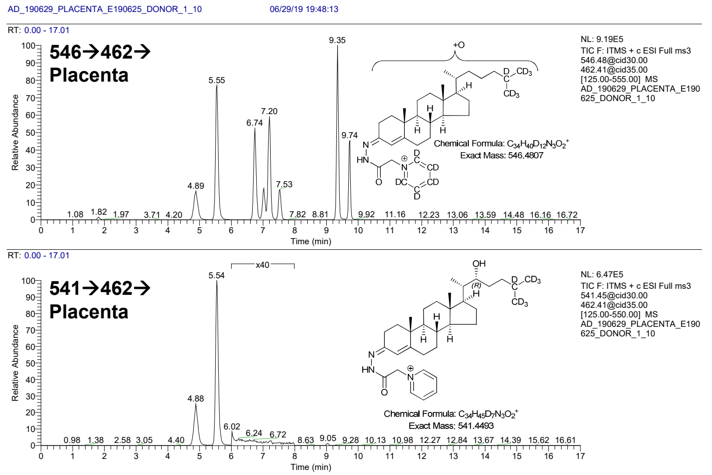


8

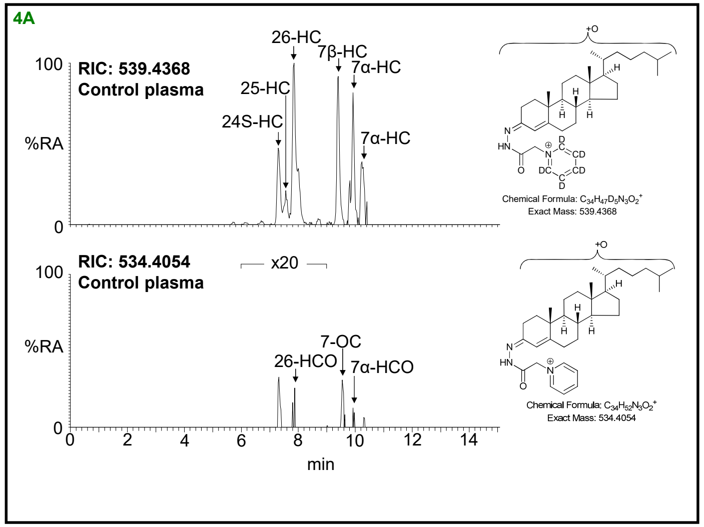
4A
RIC: 539.4368
Control plasma
26-HC
7β-HC
7α-HC
25-HC
24S-HC
7α-HC
%RA
100
0
RIC: 534.4054
Control plasma
x20
7-OC
26-HCO
7α-HCO
0 2 4 6 8 10 12 14
min
Chemical Formula: C34H47D5N3O2+
Exact Mass: 539.4368
Chemical Formula: C34H52N3O2+
Exact Mass: 534.4054

1

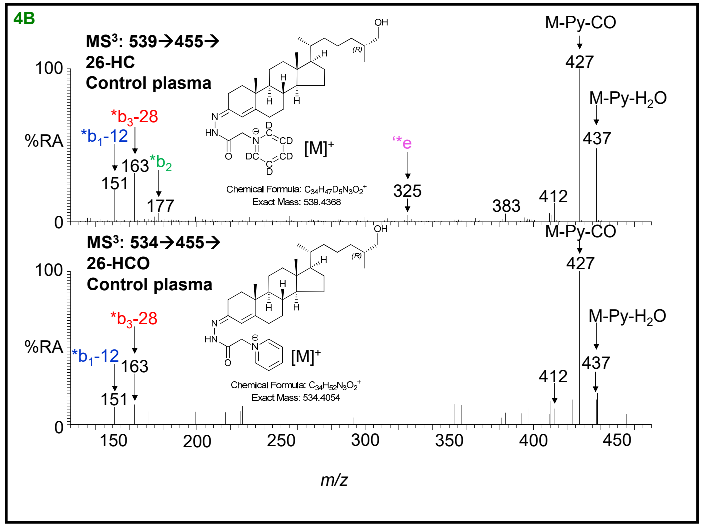
4B
MS3: 539→455→
26-HC
Control plasma
M-Py-CO
427
M-Py-H2O
437
*b3-28
*b1-12
163
*b2
151
177
'*e
325
383
412
[M]+
%RA
100
0
MS3: 534→455→
26-HCO
Control plasma
150 200 250 300 350 400 450
m/z
Chemical Formula: C34H47D5N3O2+
Exact Mass: 539.4368
Chemical Formula: C34H52N3O2+
Exact Mass: 534.4054

2

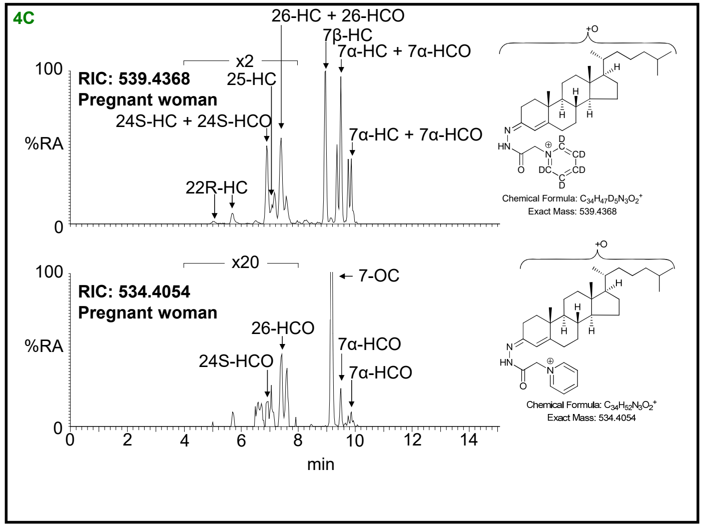
4C
RIC: 539.4368
Pregnant woman
26-HC + 26-HCO
7β-HC
7α-HC + 7α-HCO
x2
25-HC
24S-HC + 24S-HCO
7α-HC + 7α-HCO
22R-HC
%RA
100
0
RIC: 534.4054
Pregnant woman
x20
7-OC
26-HCO
24S-HCO
7α-HCO
7α-HCO
0 2 4 6 8 10 12 14
min
Chemical Formula: C34H47D5N3O2+
Exact Mass: 539.4368
Chemical Formula: C34H52N3O2+
Exact Mass: 534.4054

3

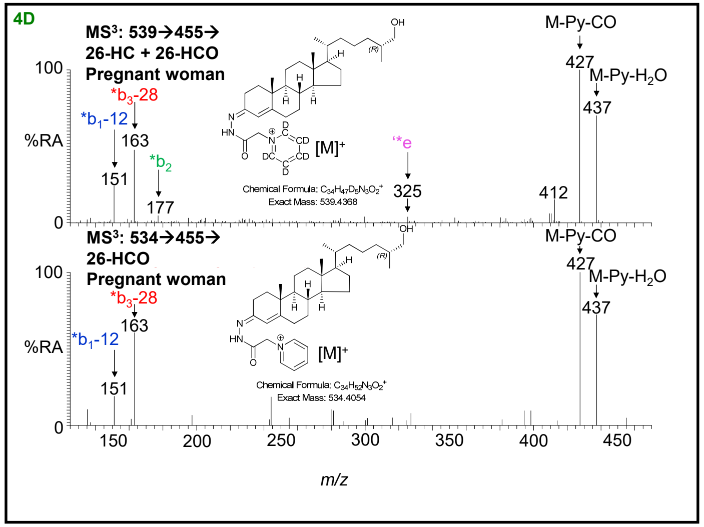
4D
MS3: 539→455→
26-HC + 26-HCO
Pregnant woman
M-Py-CO
427
M-Py-H2O
437
*b3-28
*b1-12
163
151
*b2
177
'*e
325
412
[M]+
%RA
100
0
MS3: 534→455→
26-HCO
Pregnant woman
150 200 250 300 350 400 450
m/z
Chemical Formula: C34H47D5N3O2+
Exact Mass: 539.4368
Chemical Formula: C34H52N3O2+
Exact Mass: 534.4054

4

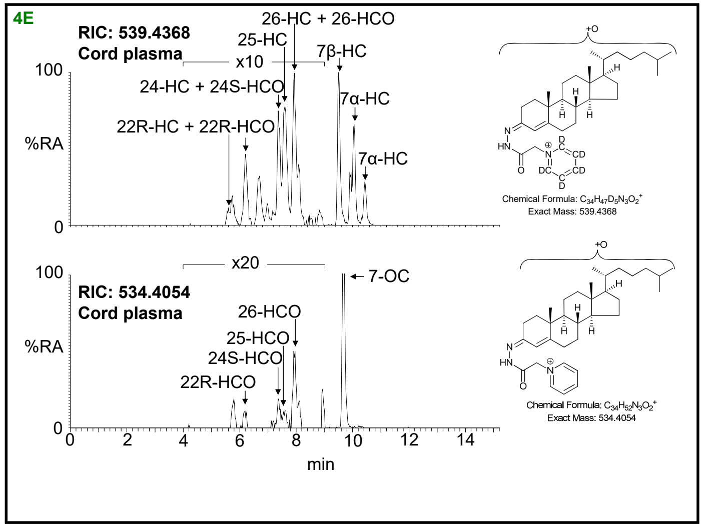
4E
RIC: 539.4368
Cord plasma
26-HC + 26-HCO
25-HC
x10
7β-HC
24-HC + 24S-HCO
7α-HC
22R-HC + 22R-HCO
7α-HC
%RA
RIC: 534.4054
Cord plasma
x20
7-OC
26-HCO
25-HCO
24S-HCO
22R-HCO
min
Chemical Formula: C34H47D5N3O2+
Exact Mass: 539.4368
Chemical Formula: C34H52N3O2+
Exact Mass: 534.4054

5

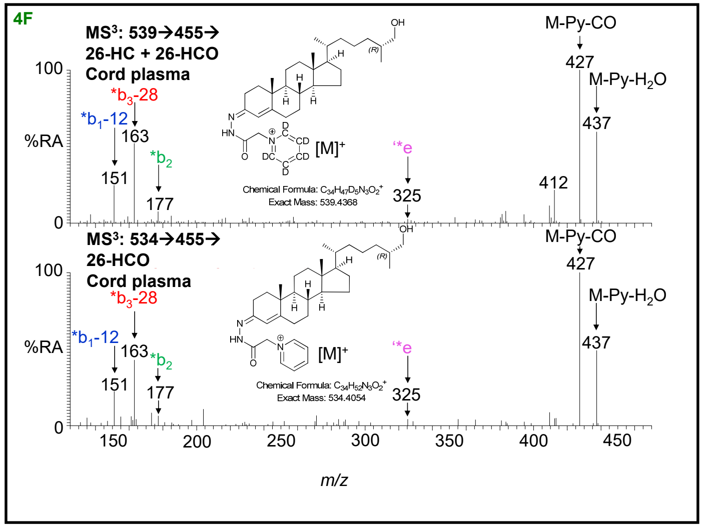
4F
MS3: 539→455→
26-HC + 26-HCO
Cord plasma
M-Py-CO
427
M-Py-H2O
437
*b3-28
*b1-12
163
151
*b2
177
[M]+
*e
325
412
Chemical Formula: C34H47D5N3O2+
Exact Mass: 539.4368
MS3: 534→455→
26-HCO
Cord plasma
Chemical Formula: C34H52N3O2+
Exact Mass: 534.4054
%RA
m/z

6

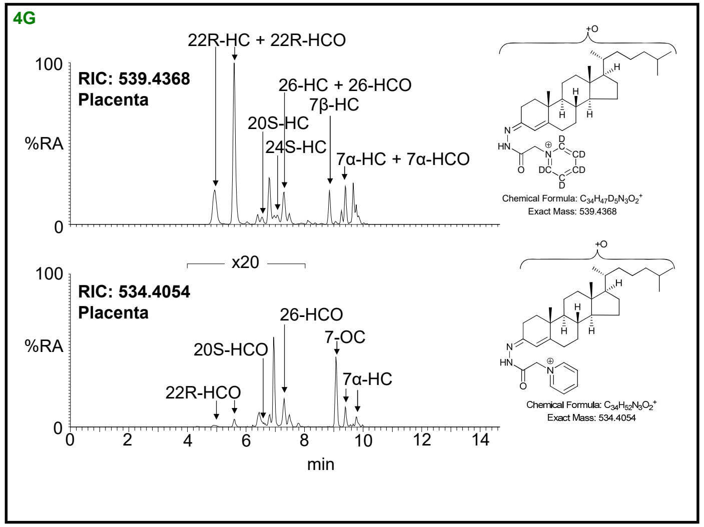
4G
RIC: 539.4368
Placenta
22R-HC + 22R-HCO
26-HC + 26-HCO
7β-HC
20S-HC
24S-HC
7α-HC + 7α-HCO
%RA
RIC: 534.4054
Placenta
x20
26-HCO
7-OC
20S-HCO
7α-HC
22R-HCO
min
Chemical Formula: C34H47D5N3O2+
Exact Mass: 539.4368
Chemical Formula: C34H52N3O2+
Exact Mass: 534.4054

7

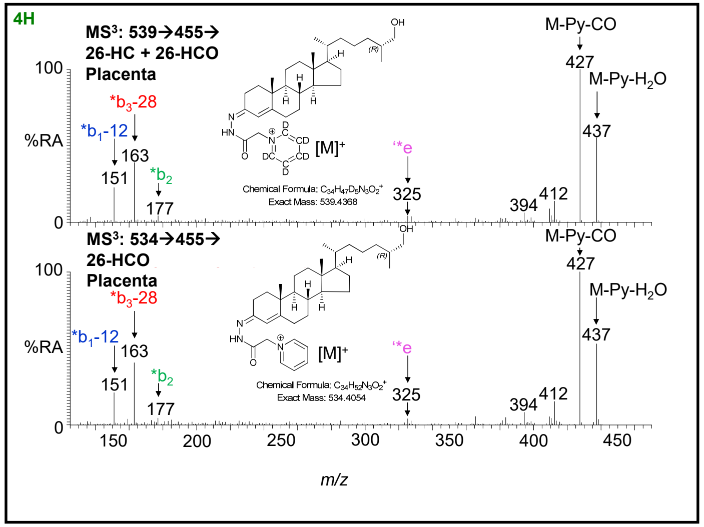
4H
MS3: 539→455→
26-HC + 26-HCO
Placenta
M-Py-CO
427
M-Py-H2O
437
*b3-28
*b1-12
163
151
*b2
177
[M]+
*e
325
394
412
Chemical Formula: C34H47D5N3O2+
Exact Mass: 539.4368
MS3: 534→455→
26-HCO
Placenta
Chemical Formula: C34H52N3O2+
Exact Mass: 534.4054
%RA
m/z

8

01/04/2022

**S5A**

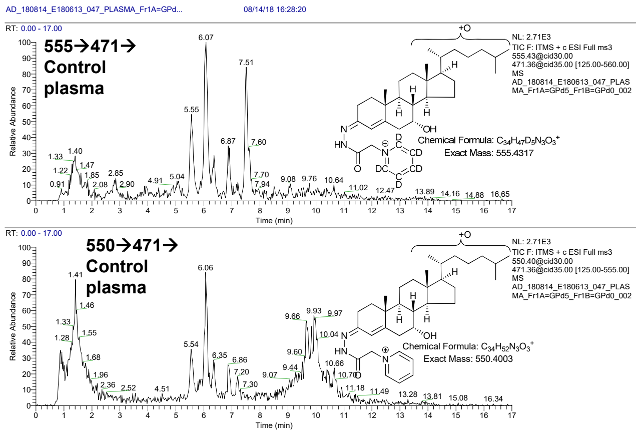


1

**S5B**

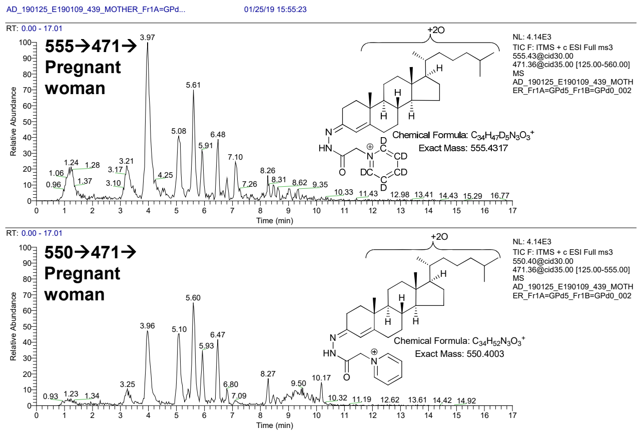


2

**S5C**

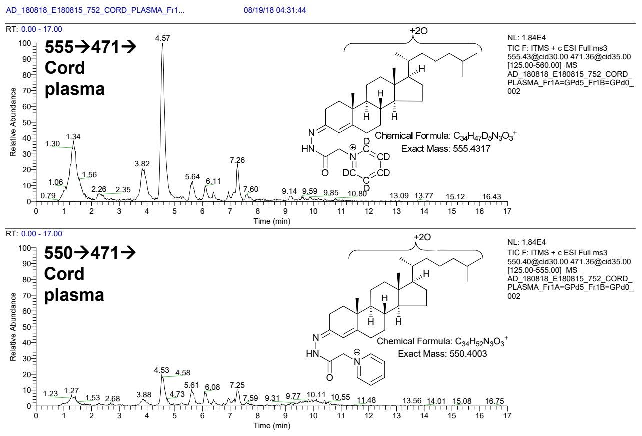


3

**S5D**

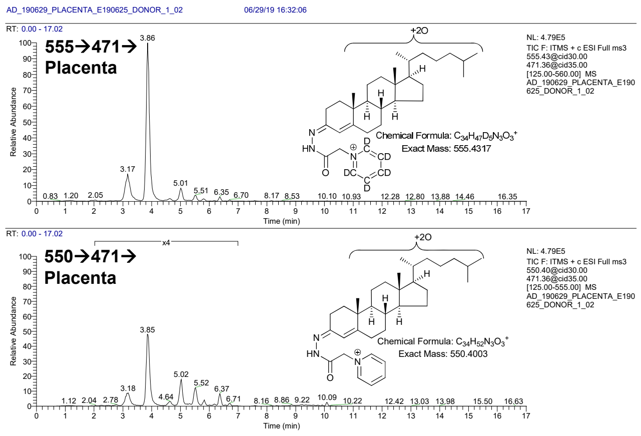


4

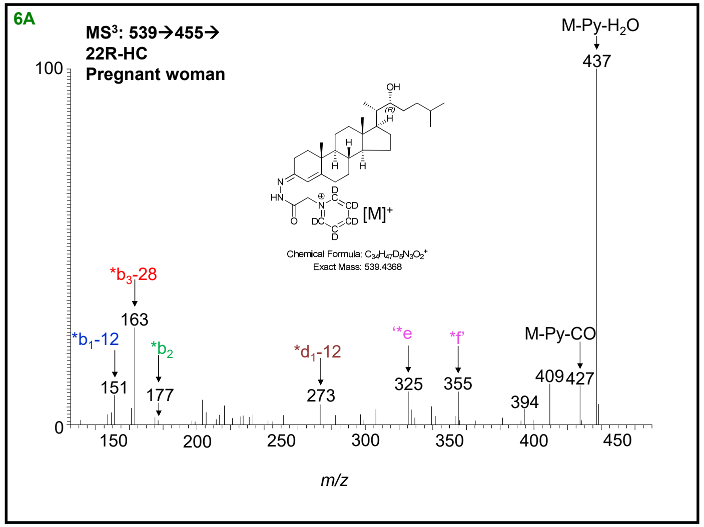
6A
MS³: 539→455→
22R-HC
Pregnant woman
M-Py-H₂O
437
*b₃-28
163
*b₁-12
151
*b₂
177
*d₁-12
273
325
355
M-Py-CO
409
427
394
[M]⁺
m/z

1

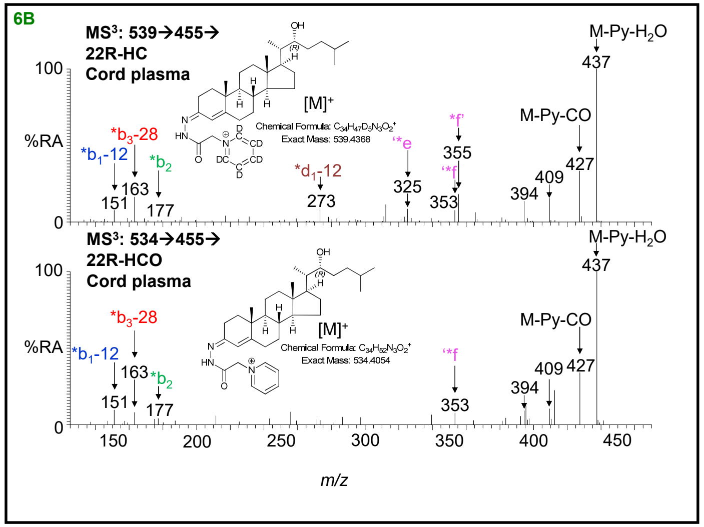
6B
MS³: 539→455→
22R-HC
Cord plasma
MS³: 534→455→
22R-HCO
Cord plasma
M-Py-H₂O
437
M-Py-CO
427
409
394
%RA
*b₃-28
163
*b₁-12
151
*b₂
177
*d₁-12
273
325
355
353
[M]⁺
m/z

2

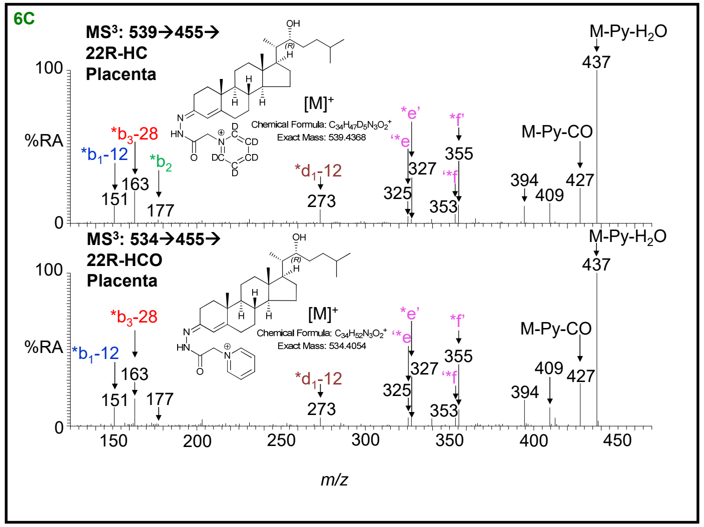
6C
MS³: 539→455→
22R-HC
Placenta
MS³: 534→455→
22R-HCO
Placenta
M-Py-H₂O
437
M-Py-CO
427
409
394
%RA
*b₃-28
163
*b₁-12
151
*b₂
177
*d₁-12
273
327
325
355
353
[M]⁺
m/z

3

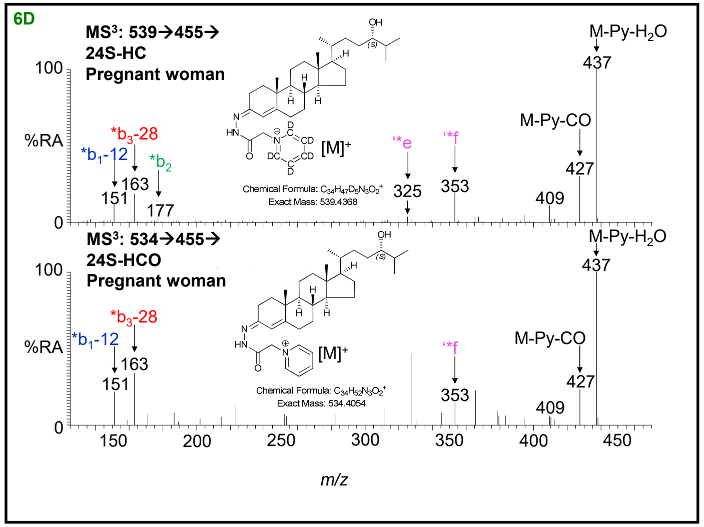
6D
MS³: 539→455→
24S-HC
Pregnant woman
MS³: 534→455→
24S-HCO
Pregnant woman
M-Py-H₂O
437
M-Py-CO
427
409
%RA
*b₃-28
163
*b₁-12
151
*b₂
177
325
353
[M]⁺
m/z

4

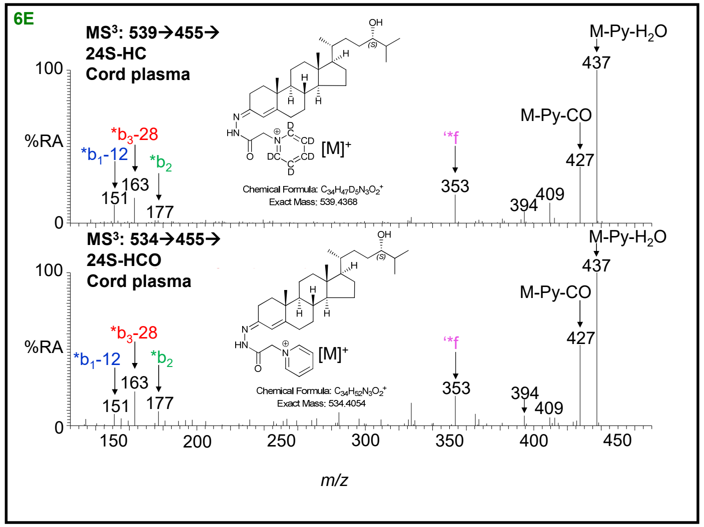
6E
MS³: 539→455→
24S-HC
Cord plasma
MS³: 534→455→
24S-HCO
Cord plasma
M-Py-H2O
M-Py-CO
[M]+
%RA
m/z

5

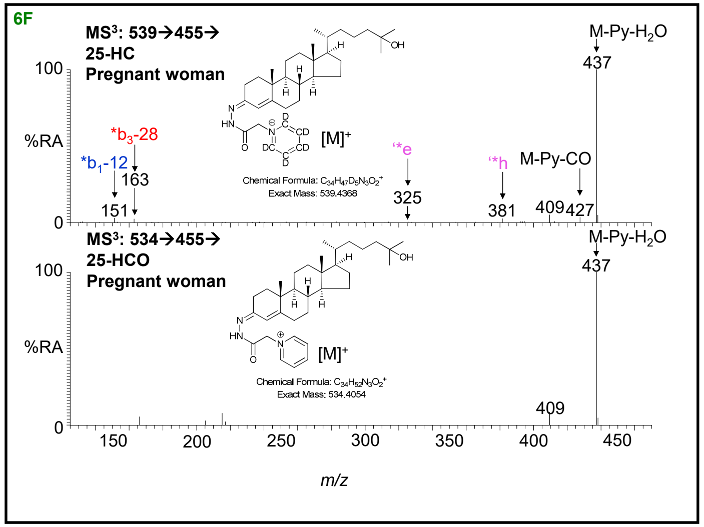
6F
MS³: 539→455→
25-HC
Pregnant woman
MS³: 534→455→
25-HCO
Pregnant woman
M-Py-H2O
M-Py-CO
[M]+
%RA
m/z

6

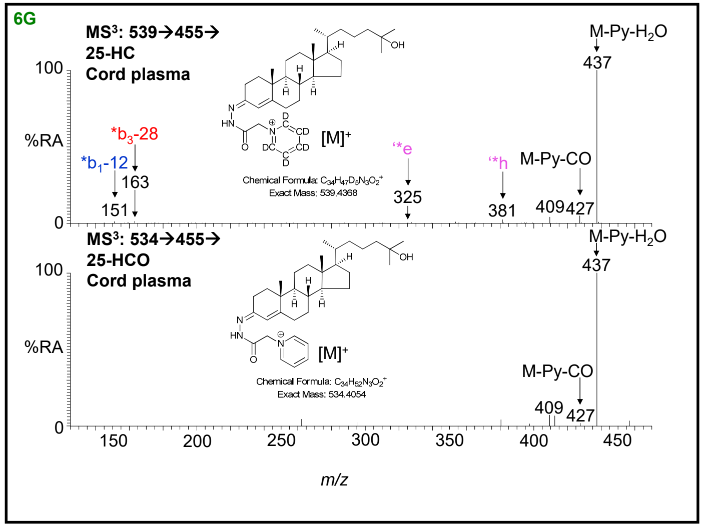
6G
MS³: 539→455→
25-HC
Cord plasma
MS³: 534→455→
25-HCO
Cord plasma
M-Py-H2O
M-Py-CO
[M]+
%RA
m/z

7

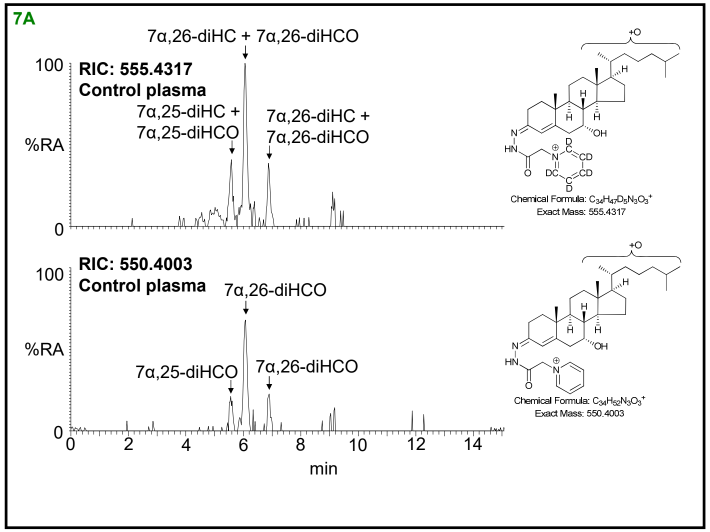

7A
7α,26-diHC + 7α,26-diHCO
RIC: 555.4317
Control plasma
7α,25-diHC +
7α,25-diHCO
7α,26-diHC +
7α,26-diHCO
%RA
100
0
Chemical Formula: C34H47D5N3O3+
Exact Mass: 555.4317
RIC: 550.4003
Control plasma
7α,26-diHCO
7α,25-diHCO
7α,26-diHCO
Chemical Formula: C34H52N3O3+
Exact Mass: 550.4003
0 2 4 6 8 10 12 14
min


1

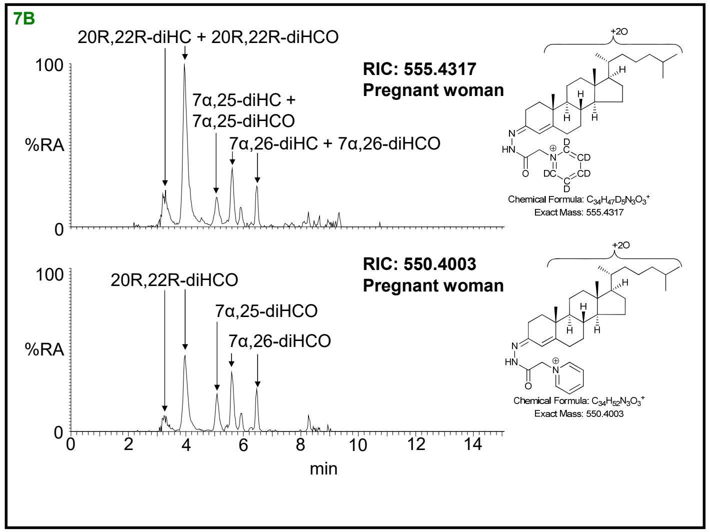

7B
20R,22R-diHC + 20R,22R-diHCO
RIC: 555.4317
Pregnant woman
7α,25-diHC +
7α,25-diHCO
7α,26-diHC + 7α,26-diHCO
%RA
100
0
Chemical Formula: C34H47D5N3O3+
Exact Mass: 555.4317
RIC: 550.4003
Pregnant woman
20R,22R-diHCO
7α,25-diHCO
7α,26-diHCO
Chemical Formula: C34H52N3O3+
Exact Mass: 550.4003
0 2 4 6 8 10 12 14
min


2

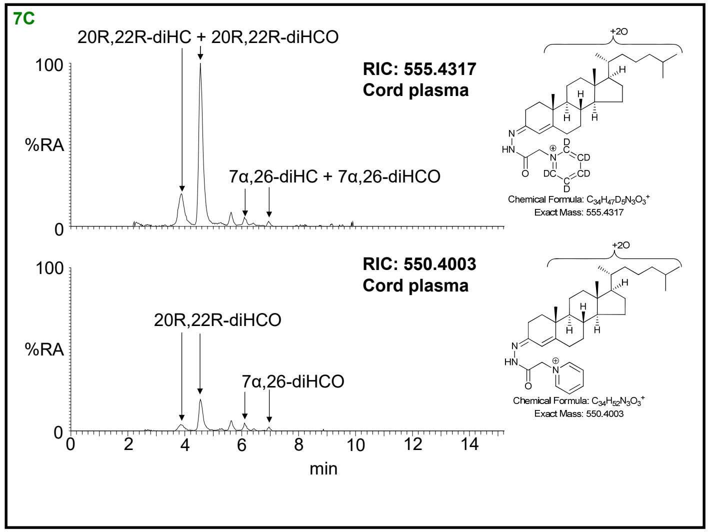

7C
20R,22R-diHC + 20R,22R-diHCO
RIC: 555.4317
Cord plasma
7α,26-diHC + 7α,26-diHCO
%RA
100
0
Chemical Formula: C34H47D5N3O3+
Exact Mass: 555.4317
RIC: 550.4003
Cord plasma
20R,22R-diHCO
7α,26-diHCO
Chemical Formula: C34H52N3O3+
Exact Mass: 550.4003
0 2 4 6 8 10 12 14
min


3

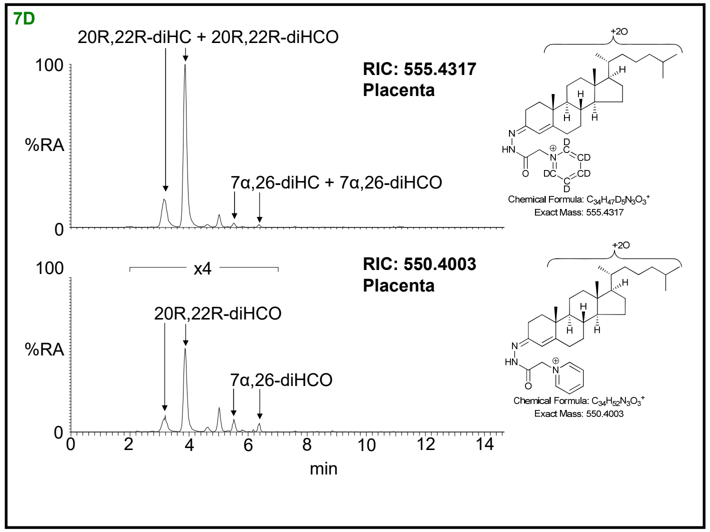

7D
20R,22R-diHC + 20R,22R-diHCO
RIC: 555.4317
Placenta
7α,26-diHC + 7α,26-diHCO
%RA
100
0
Chemical Formula: C34H47D5N3O3+
Exact Mass: 555.4317
x4
RIC: 550.4003
Placenta
20R,22R-diHCO
7α,26-diHCO
Chemical Formula: C34H52N3O3+
Exact Mass: 550.4003
0 2 4 6 8 10 12 14
min


4

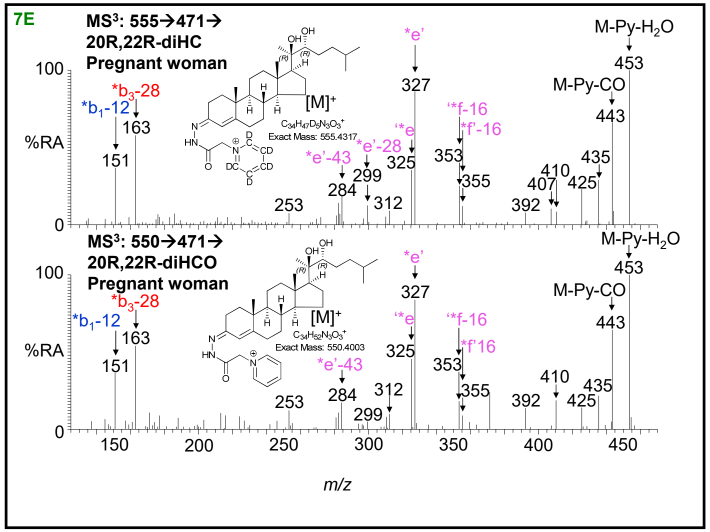
7E
MS³: 555→471→
20R,22R-diHC
Pregnant woman
MS³: 550→471→
20R,22R-diHCO
Pregnant woman
M-Py-H2O
M-Py-CO
[M]+
C34H47D5N3O3+
Exact Mass: 555.4317
C34H52N3O3+
Exact Mass: 550.4003
*b3-28
*b1-12
*e'
*e'-28
*e'-43
*f-16
*f'-16
%RA
m/z

5

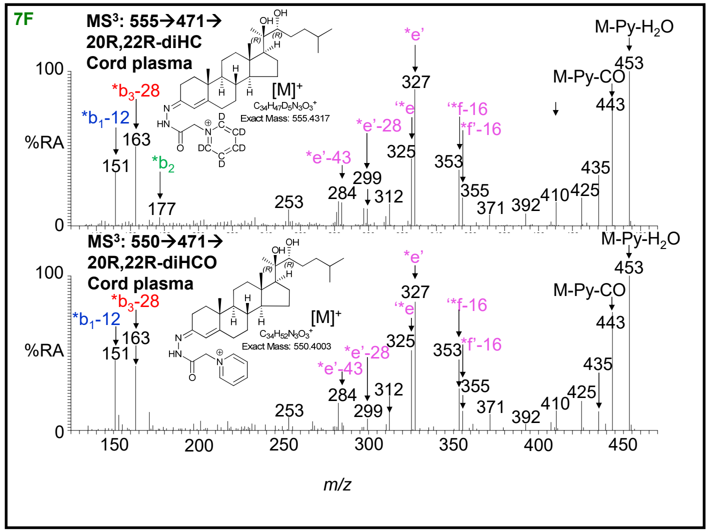
7F
MS³: 555→471→
20R,22R-diHC
Cord plasma
MS³: 550→471→
20R,22R-diHCO
Cord plasma
M-Py-H2O
M-Py-CO
[M]+
C34H47D5N3O3+
Exact Mass: 555.4317
C34H52N3O3+
Exact Mass: 550.4003
*b3-28
*b1-12
*b2
*e'
*e'-28
*e'-43
*f-16
*f'-16
%RA
m/z

6

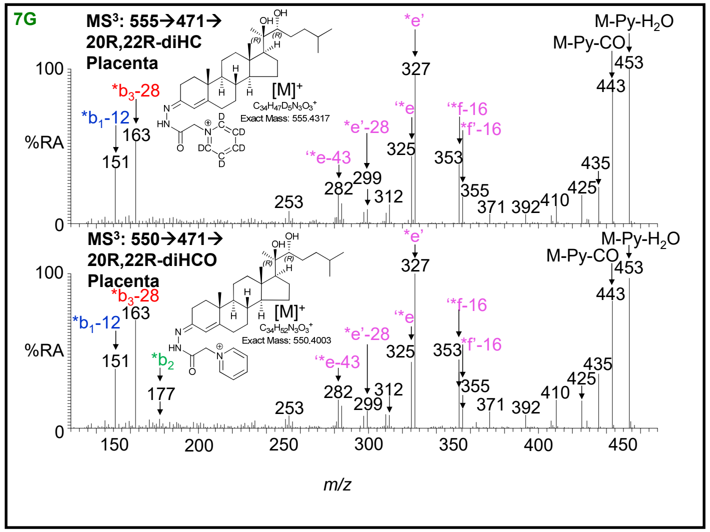
7G
MS³: 555→471→
20R,22R-diHC
Placenta
MS³: 550→471→
20R,22R-diHCO
Placenta
M-Py-H2O
M-Py-CO
[M]+
C34H47D5N3O3+
Exact Mass: 555.4317
C34H52N3O3+
Exact Mass: 550.4003
*b3-28
*b1-12
*b2
*e'
*e'-28
*e-43
*f-16
*f'-16
%RA
m/z

7

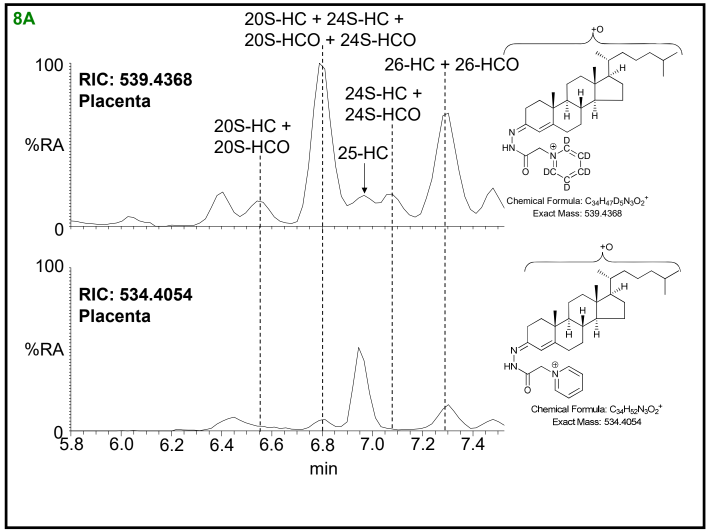
8A
20S-HC + 24S-HC +
20S-HCO + 24S-HCO
26-HC + 26-HCO
RIC: 539.4368
Placenta
24S-HC +
24S-HCO
20S-HC +
20S-HCO
25-HC
%RA
RIC: 534.4054
Placenta
Chemical Formula: C34H47D5N3O2+
Exact Mass: 539.4368
Chemical Formula: C34H52N3O2+
Exact Mass: 534.4054
min

1

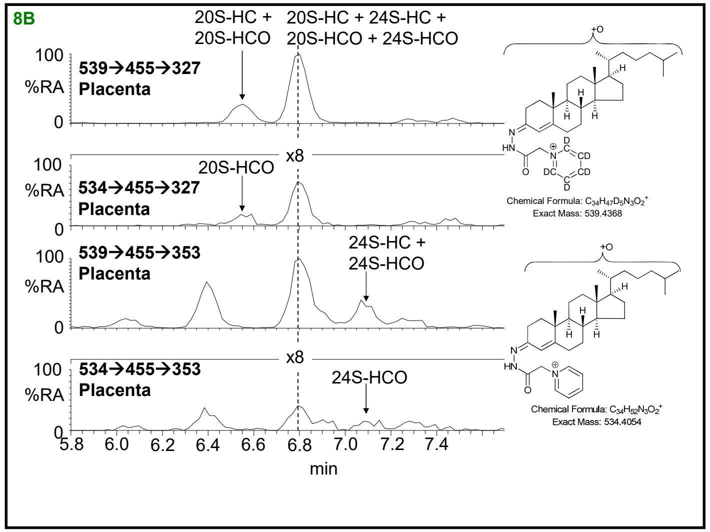
8B
20S-HC +
20S-HCO
20S-HC + 24S-HC +
20S-HCO + 24S-HCO
539→455→327
Placenta
534→455→327
Placenta
20S-HCO
x8
539→455→353
Placenta
24S-HC +
24S-HCO
534→455→353
Placenta
24S-HCO
%RA
min
Chemical Formula: C34H47D5N3O2+
Exact Mass: 539.4368
Chemical Formula: C34H52N3O2+
Exact Mass: 534.4054

2

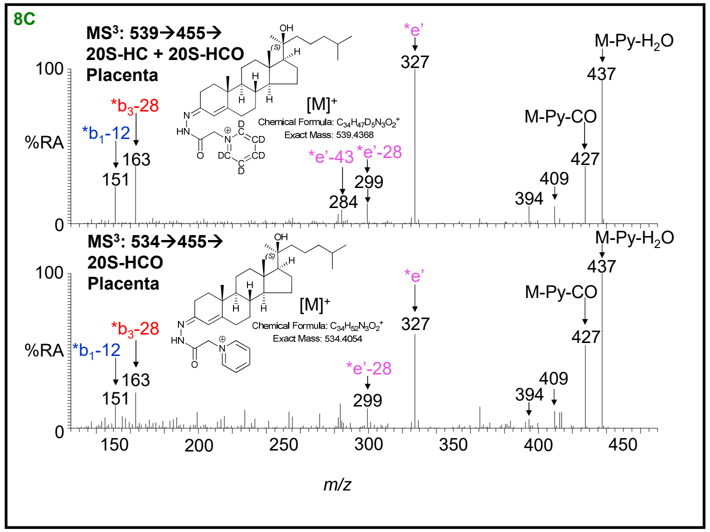
8C
MS3: 539→455→
20S-HC + 20S-HCO
Placenta
*e'
M-Py-H2O
[M]+
Chemical Formula: C34H47D5N3O2+
Exact Mass: 539.4368
*b3-28
*b1-12
M-Py-CO
*e'-43
*e'-28
MS3: 534→455→
20S-HCO
Placenta
Chemical Formula: C34H52N3O2+
Exact Mass: 534.4054
%RA
m/z

3

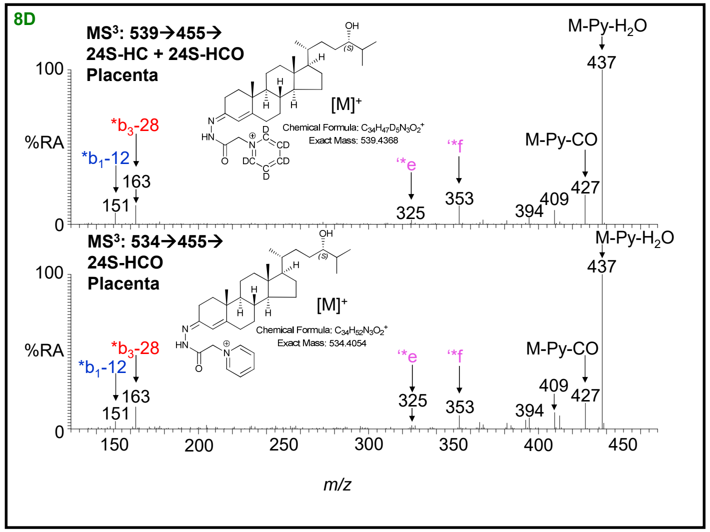
8D
MS3: 539→455→
24S-HC + 24S-HCO
Placenta
M-Py-H2O
[M]+
Chemical Formula: C34H47D5N3O2+
Exact Mass: 539.4368
*b3-28
*b1-12
'*e
'*f
M-Py-CO
MS3: 534→455→
24S-HCO
Placenta
Chemical Formula: C34H52N3O2+
Exact Mass: 534.4054
%RA
m/z

4

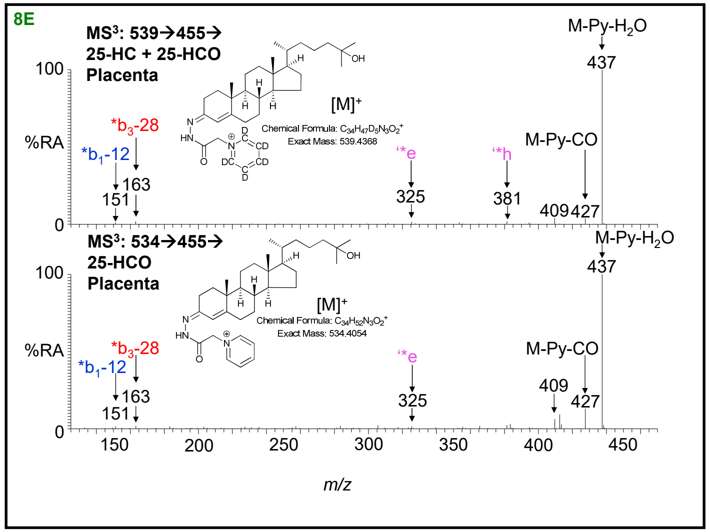
8E
MS3: 539→455→
25-HC + 25-HCO
Placenta
M-Py-H2O
437
100
%RA
0
[M]+
*b3-28
*b1-12
163
151
‘*e
325
‘*h
381
M-Py-CO
409
427
MS3: 534→455→
25-HCO
Placenta
M-Py-H2O
437
100
%RA
0
[M]+
*b3-28
*b1-12
163
151
‘*e
325
M-Py-CO
409
427
150
200
250
300
350
400
450
m/z

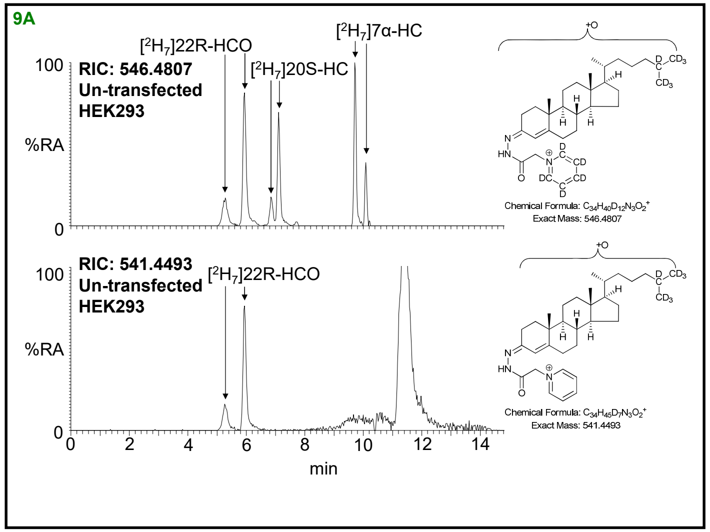
9A
RIC: 546.4807
Un-transfected
HEK293
[$^2H_7$]22R-HCO
[$^2H_7$]20S-HC
[$^2H_7$]7α-HC
%RA
100
0
RIC: 541.4493
Un-transfected
HEK293
[$^2H_7$]22R-HCO
0
2
4
6
8
10
12
14
min
Chemical Formula: $C_{34}H_{40}D_{12}N_3O_2^+$
Exact Mass: 546.4807
Chemical Formula: $C_{34}H_{45}D_7N_3O_2^+$
Exact Mass: 541.4493

1

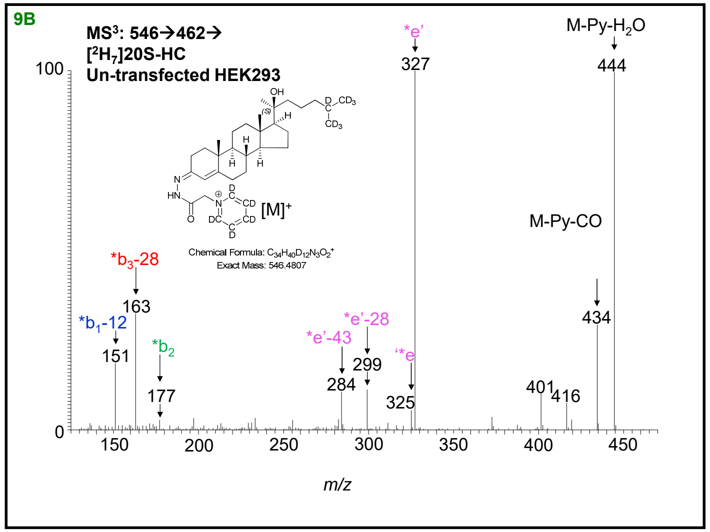
9B
MS$^3$: 546→462→
[$^2H_7$]20S-HC
Un-transfected HEK293
[M]$^+$
Chemical Formula: $C_{34}H_{40}D_{12}N_3O_2^+$
Exact Mass: 546.4807
*e'
327
M-Py-$H_2O$
444
M-Py-CO
434
*$b_3$-28
163
*$b_1$-12
151
*$b_2$
177
*e'-43
284
*e'-28
299
*e
325
401
416
100
0
150
200
250
300
350
400
450
m/z

2

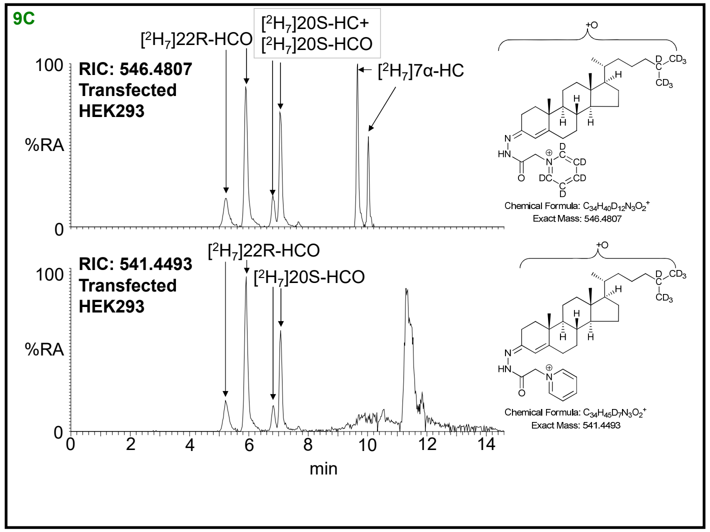
9C
RIC: 546.4807
Transfected
HEK293
[$^2H_7$]22R-HCO
[$^2H_7$]20S-HC+
[$^2H_7$]20S-HCO
[$^2H_7$]7α-HC
%RA
100
0
RIC: 541.4493
Transfected
HEK293
[$^2H_7$]22R-HCO
[$^2H_7$]20S-HCO
0
2
4
6
8
10
12
14
min
Chemical Formula: $C_{34}H_{40}D_{12}N_3O_2^+$
Exact Mass: 546.4807
Chemical Formula: $C_{34}H_{45}D_7N_3O_2^+$
Exact Mass: 541.4493

3

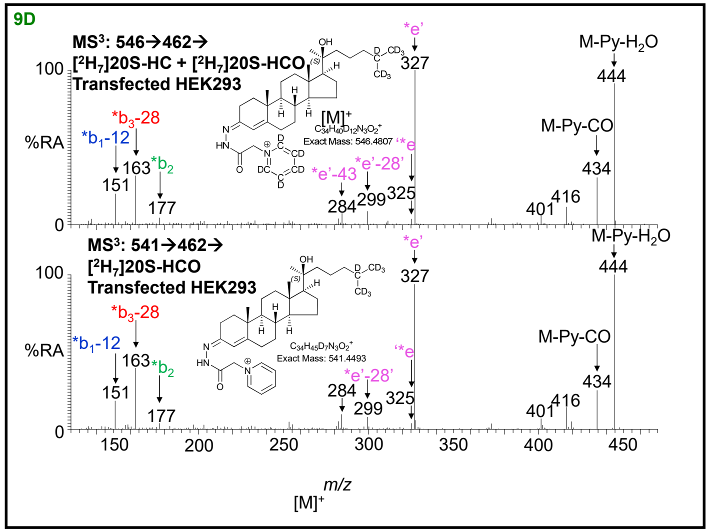
9D
MS$^3$: 546→462→
[$^2H_7$]20S-HC + [$^2H_7$]20S-HCO
Transfected HEK293
[M]$^+$
$C_{34}H_{40}D_{12}N_3O_2^+$
Exact Mass: 546.4807
*e'
327
M-Py-$H_2O$
444
M-Py-CO
434
*$b_3$-28
163
*$b_1$-12
151
*$b_2$
177
*e'-43
284
*e'-28'
299
*e
325
401
416
%RA
100
0
MS$^3$: 541→462→
[$^2H_7$]20S-HCO
Transfected HEK293
$C_{34}H_{45}D_7N_3O_2^+$
Exact Mass: 541.4493
*e'
327
M-Py-$H_2O$
444
M-Py-CO
434
*$b_3$-28
163
*$b_1$-12
151
*$b_2$
177
*e'-28'
284
299
*e
325
401
416
150
200
250
300
350
400
450
m/z
[M]$^+$

4

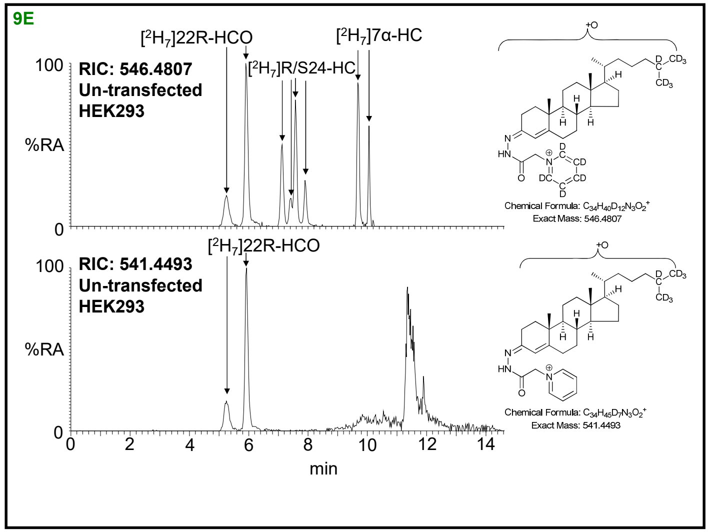
9E
[2H7]22R-HCO
[2H7]R/S24-HC
[2H7]7α-HC
RIC: 546.4807
Un-transfected
HEK293
%RA
RIC: 541.4493
Un-transfected
HEK293
min
Chemical Formula: C34H40D12N3O2+
Exact Mass: 546.4807
Chemical Formula: C34H45D7N3O2+
Exact Mass: 541.4493

5

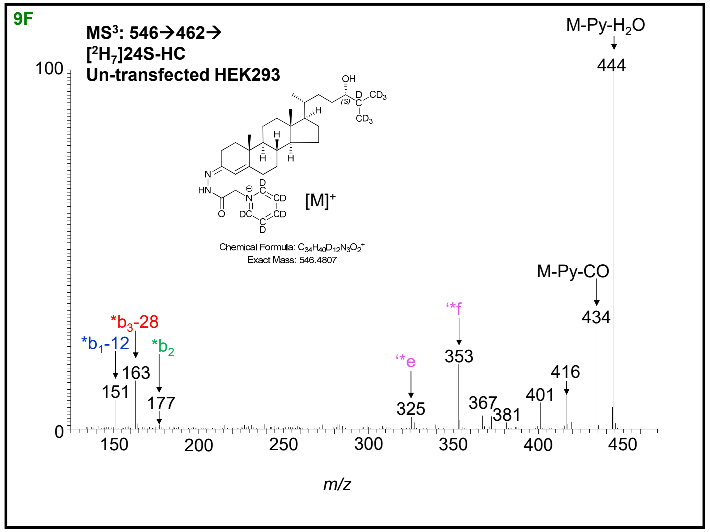
9F
MS3: 546→462→
[2H7]24S-HC
Un-transfected HEK293
M-Py-H2O
444
M-Py-CO
434
[M]+
Chemical Formula: C34H40D12N3O2+
Exact Mass: 546.4807
*b3-28
*b1-12
*b2
151
163
177
325
353
367
381
401
416
m/z

6

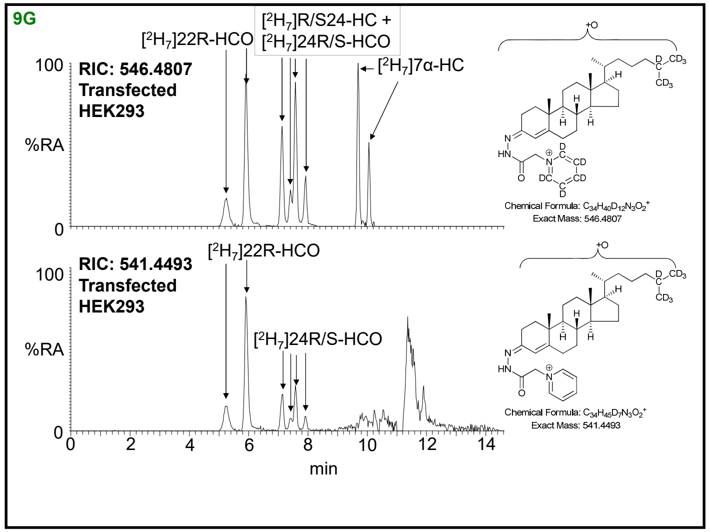
9G
[2H7]22R-HCO
[2H7]R/S24-HC +
[2H7]24R/S-HCO
[2H7]7α-HC
RIC: 546.4807
Transfected
HEK293
%RA
RIC: 541.4493
Transfected
HEK293
[2H7]24R/S-HCO
min
Chemical Formula: C34H40D12N3O2+
Exact Mass: 546.4807
Chemical Formula: C34H45D7N3O2+
Exact Mass: 541.4493

7

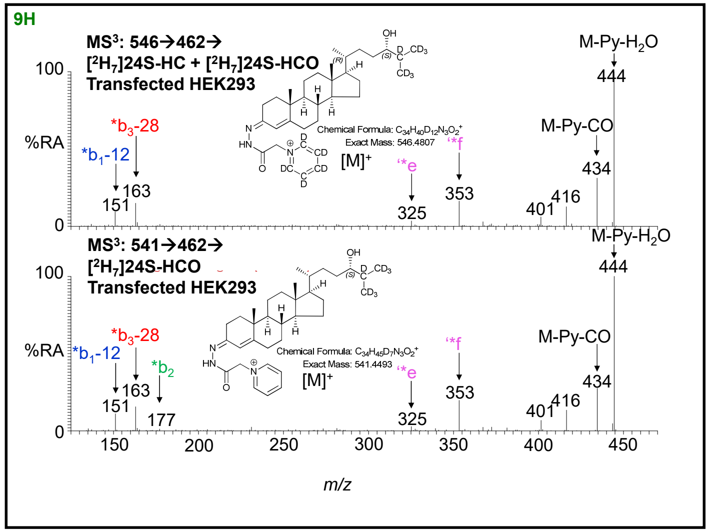
9H
MS3: 546→462→
[2H7]24S-HC + [2H7]24S-HCO
Transfected HEK293
M-Py-H2O
444
M-Py-CO
434
[M]+
Chemical Formula: C34H40D12N3O2+
Exact Mass: 546.4807
*b3-28
*b1-12
151
163
325
353
401
416
MS3: 541→462→
[2H7]24S-HCO
Transfected HEK293
Chemical Formula: C34H45D7N3O2+
Exact Mass: 541.4493
*b2
177
%RA
m/z

8

10

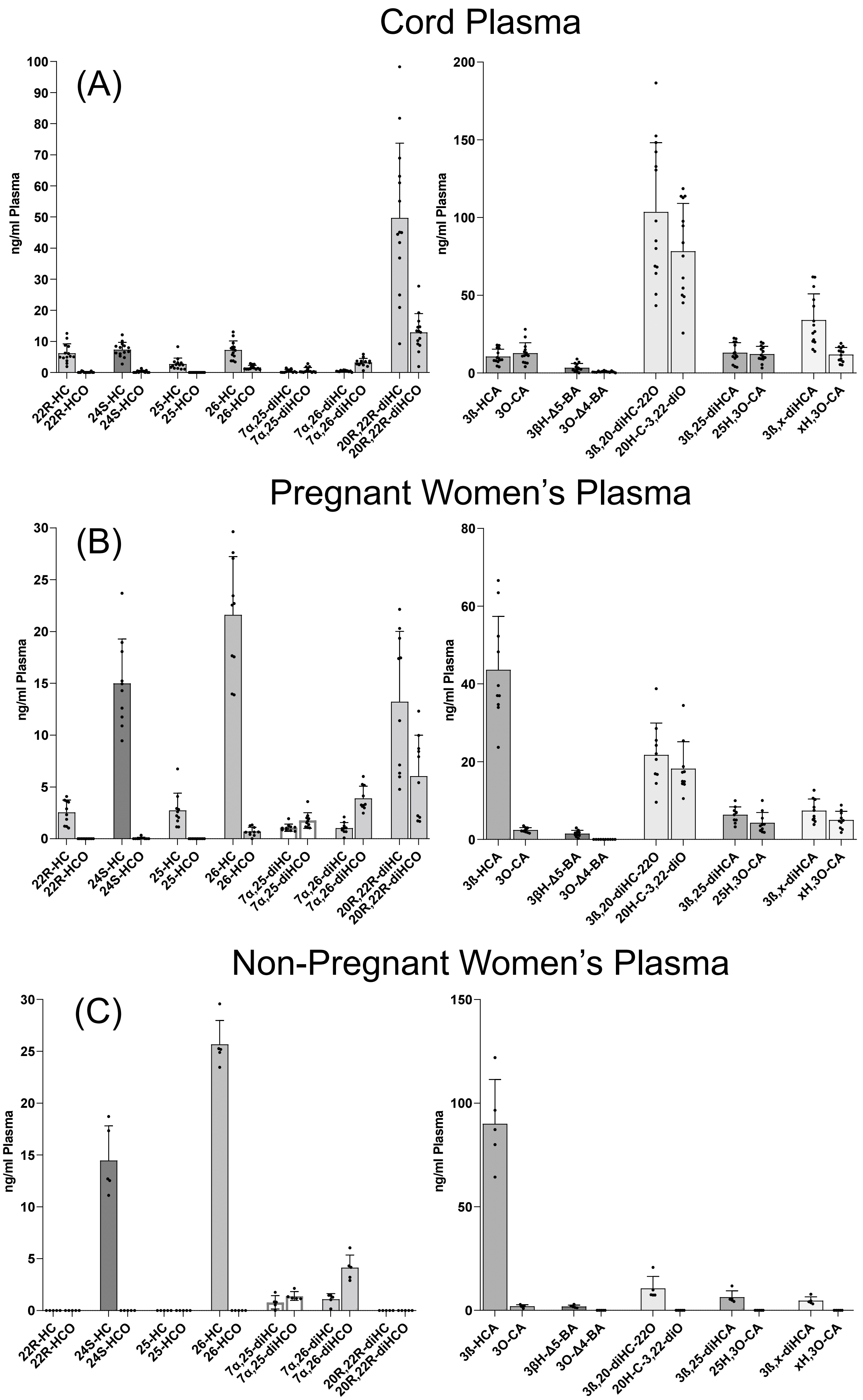
Cord Plasma
(A)
ng/ml Plasma
22R-HC
22R-HCO
24S-HC
24S-HCO
25-HC
25-HCO
26-HC
26-HCO
7α,25-diHC
7α,25-diHCO
7α,26-diHC
7α,26-diHCO
20R,22R-diHC
20R,22R-diHCO
3ß-HCA
3O-CA
3ßH-Δ5-BA
3O-Δ4-BA
3ß,20-diHC-22O
20H-C-3,22-diO
3ß,25-diHCA
25H,3O-CA
3ß,x-diHCA
xH,3O-CA
Pregnant Women's Plasma
(B)
Non-Pregnant Women's Plasma
(C)

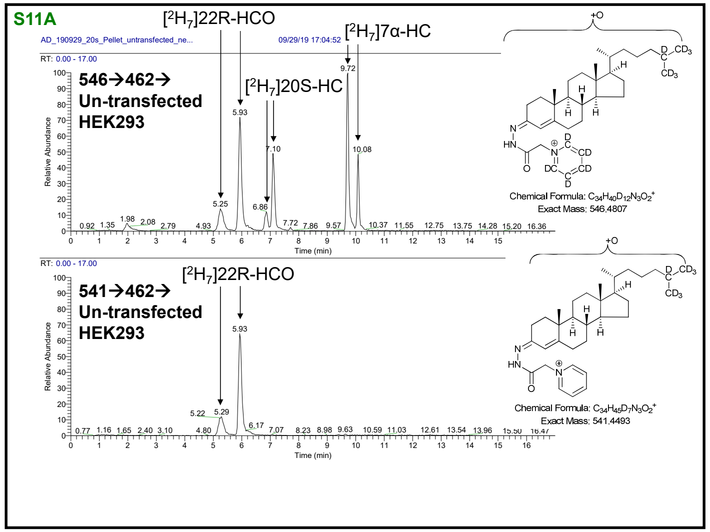
S11A
[2H7]22R-HCO
[2H7]7α-HC
[2H7]20S-HC
546→462→
Un-transfected
HEK293
541→462→
Un-transfected
HEK293
Chemical Formula: C34H40D12N3O2+
Exact Mass: 546.4807
Chemical Formula: C34H45D7N3O2+
Exact Mass: 541.4493
Relative Abundance
Time (min)

1

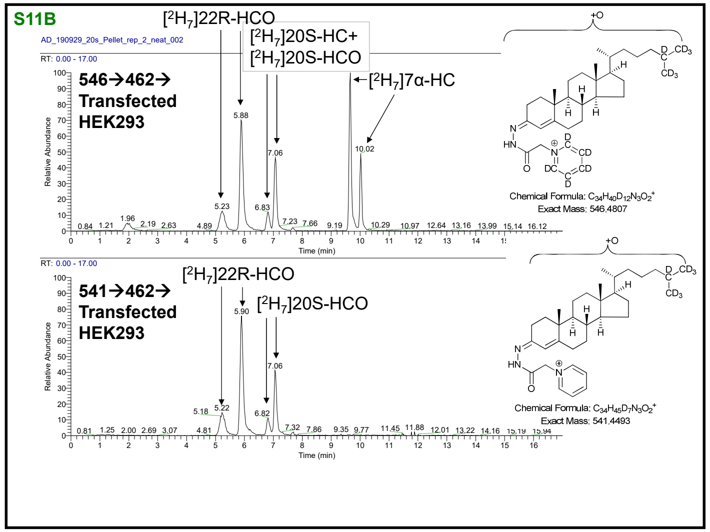
S11B
[2H7]22R-HCO
[2H7]20S-HC+
[2H7]20S-HCO
[2H7]7α-HC
546→462→
Transfected
HEK293
541→462→
Transfected
HEK293
[2H7]20S-HCO
Chemical Formula: C34H40D12N3O2+
Exact Mass: 546.4807
Chemical Formula: C34H45D7N3O2+
Exact Mass: 541.4493
Relative Abundance
Time (min)

2

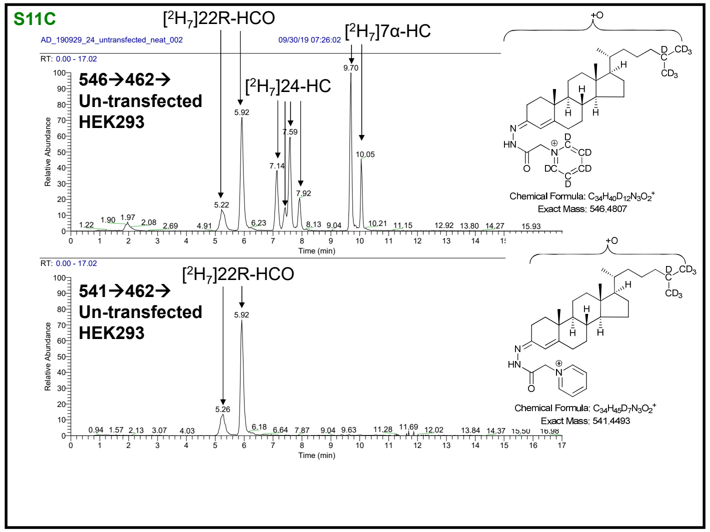
S11C
[2H7]22R-HCO
[2H7]7α-HC
[2H7]24-HC
546→462→
Un-transfected
HEK293
541→462→
Un-transfected
HEK293
Chemical Formula: C34H40D12N3O2+
Exact Mass: 546.4807
Chemical Formula: C34H45D7N3O2+
Exact Mass: 541.4493
Relative Abundance
Time (min)

3

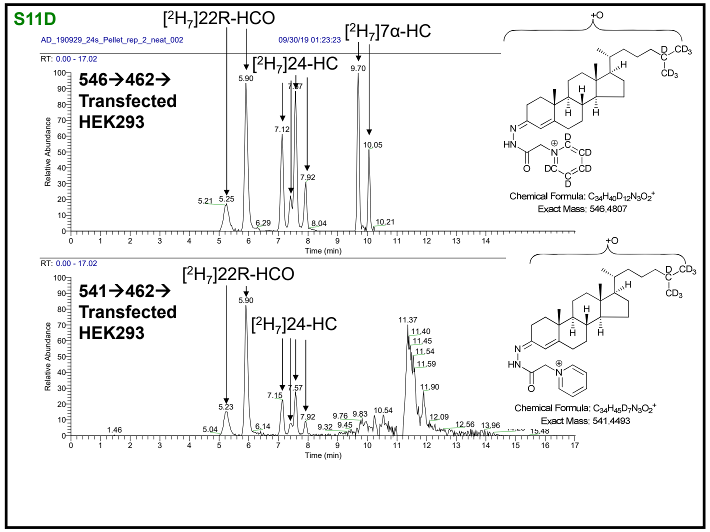
S11D
[2H7]22R-HCO
[2H7]7α-HC
[2H7]24-HC
546→462→
Transfected
HEK293
541→462→
Transfected
HEK293
Chemical Formula: C34H40D12N3O2+
Exact Mass: 546.4807
Chemical Formula: C34H45D7N3O2+
Exact Mass: 541.4493
Relative Abundance
Time (min)

4
